# Cancers relax selection on charge at the mitochondrial complex I quinone cavity

**DOI:** 10.64898/2026.10.24.761499

**Authors:** Ji-Yong Sung, Jae-Ho Cheong

## Abstract

Mitochondrial complex I translocates protons through hydrated channels whose connectivity and electrostatics are both defined by structure, but whether they are distinctly constrained is unknown. From a 2.39 Å cryo-EM structure modelling 2,945 ordered waters, we built the proton wire of the seven mtDNA-encoded ND subunits and scored all 40,166 missense substitutions on two near-orthogonal axes (r = 0.17): electrostatic shift at proton-loading sites, and topological disruption of the wire. Across 858 germline variant sites, loading sites (0.51) and cut vertices (0.67) are both depleted. In tumours only loading sites are enriched (1.42 TCGA, 1.67 PCAWG, 1.29 MSK-IMPACT), a 3.06-fold difference tested as a wire class × tissue-type interaction (P = 3.6 × 10LL). It survives adjustment for conservation, burial and coverage, and is concentrated in ND1 within 15 Å of the quinone cavity, not spread along the proton path. Selection separates charge from connectivity, without establishing a fitness benefit.

**Significance:** Mitochondrial complex I is essential for cellular energy production, yet how cancer alters evolutionary constraints on its proton-translocating machinery remains unclear. By reconstructing hydrogen-bonded proton networks from high-resolution structural data, we distinguish two fundamental properties: electrostatic regulation of proton loading and structural connectivity of proton-transfer pathways. Across three independent cancer cohorts, mutations affecting proton-loading electrostatics are enriched, whereas mutations disrupting network connectivity are not. This divergence is concentrated near the quinone-binding cavity rather than distributed along the proton pathways. Our findings reveal that cancer-associated mitochondrial evolution selectively relaxes constraints on charge regulation while preserving constraints on proton-network integrity, providing a biophysical framework for understanding how mitochondrial mutations differ between human populations and tumors.

## Introduction

Respiratory complex I couples the oxidation of NADH to the translocation of four protons across the inner mitochondrial membrane, over distances of up to 200 Å from the site where electrons enter. (1, 2) Proton translocation does not occur through a single open channel. (3) (4) Cryo-EM structures resolved to better than 2.5 Å show instead a series of hydrated cavities lining the three antiporter-like modules and the E-channel at the ND1–ND3–ND4L–ND6 junction, in which ordered water molecules and the side chains of titratable residues form hydrogen-bonded networks. (3, 5) Two physically distinct properties of such a network determine whether it can carry a proton. (2, 6) The first is electrostatic: the protonation free energy of the titratable groups that load and release the proton, which has been computed for the E-channel as an ensemble of coupled charge states rather than as independent pKa values. (7) The second is topological: whether the hydrogen-bonded chain is continuous at all, since a Grotthuss relay is interrupted by the loss of a single bridging group. (4, 8, 9) Both properties are specified by the structure, and both are in principle subject to natural selection. (3, 6)

Seven of the subunits that build this machinery — ND1 to ND6 and ND4L — are encoded by mitochondrial DNA. (10, 11) Population-scale studies have established strong purifying selection across the mitochondrial genome, with functionally constrained residues clustering in three dimensions within respiratory proteins. (10) In tumours, by contrast, mtDNA mutational patterns differ sharply between complexes: truncating variants are enriched at least two-fold in complex I and concentrate in renal, thyroid and colorectal cancers, whereas missense variation predominates in complex III. (11) We recently showed that cytochrome b missense variants predicted to perturb the electrostatics of the haem centres are purged from the germline but tolerated in tumours at low heteroplasmy, consistent with relaxed purifying selection rather than positive selection. (12–15) That analysis also exposed a methodological limit: the predicted electron-transfer perturbation was almost collinear with the electrostatic shift at the nearest redox centre (r = 0.94), so no physical property could be isolated from local electrostatics, and the question of whether a specific biophysical determinant is under selection was left open. (14, 15)

Complex I offers a way past that limit, for two reasons. (16) First, the physics is protonic rather than electronic: the mtDNA-encoded ND subunits form the membrane arm and its proton pathways, while the FMN–iron–sulfur chain that carries electrons lies almost entirely in nuclear-encoded subunits. (17) Second, and more importantly, the two properties that define a proton wire can be measured independently. (18) The electrostatic shift a substitution imposes on a loading site is computed from its charge change and its distance; the topological disruption it causes is computed from the connectivity of the hydrogen-bonded graph. (18, 19) A charge-neutral substitution can sever a wire, and a distant charge change can shift a loading site without touching one, so the two quantities are separable by construction rather than by statistical adjustment after the fact. (20)

Here we build the hydrogen-bonded proton wire of the seven mtDNA-encoded ND subunits from a 2.39 Å structure of mouse complex I that models 2,945 ordered water molecules, and score every possible missense substitution on these two axes, which prove near-orthogonal (r = 0.17). (3) Comparing 858 germline variant sites with somatic variants from three tumour cohorts, we find that the germline purges substitutions in both classes, whereas tumours enrich only those that change charge at a proton-loading site; substitutions that sever the wire are never enriched in any cohort. (10) The enrichment survives adjustment for sequence conservation, burial and side-chain physicochemistry, does not depend on heteroplasmy, and leaves no detectable oxidative signature in the nuclear genome. (12) Selection on the complex I proton wire therefore distinguishes charge from connectivity, and the constraint that tumours relax is specific to proton-loading electrostatics. (3)

## Results

### A hydrated proton-wire map of the mtDNA-encoded complex I subunits

We built the proton wire from the cryo-EM structure of mouse complex I in the active state (PDB 8OM1, 2.39 Å), which models 2,945 ordered water molecules and resolves all seven mtDNA-encoded subunits: ND1 (318 residues), ND2 (345), ND3 (115), ND4 (459), ND4L (98), ND5 (607) and ND6 (172), 2,114 residues in total (**Fig. 1b**).

**Fig. 1.**
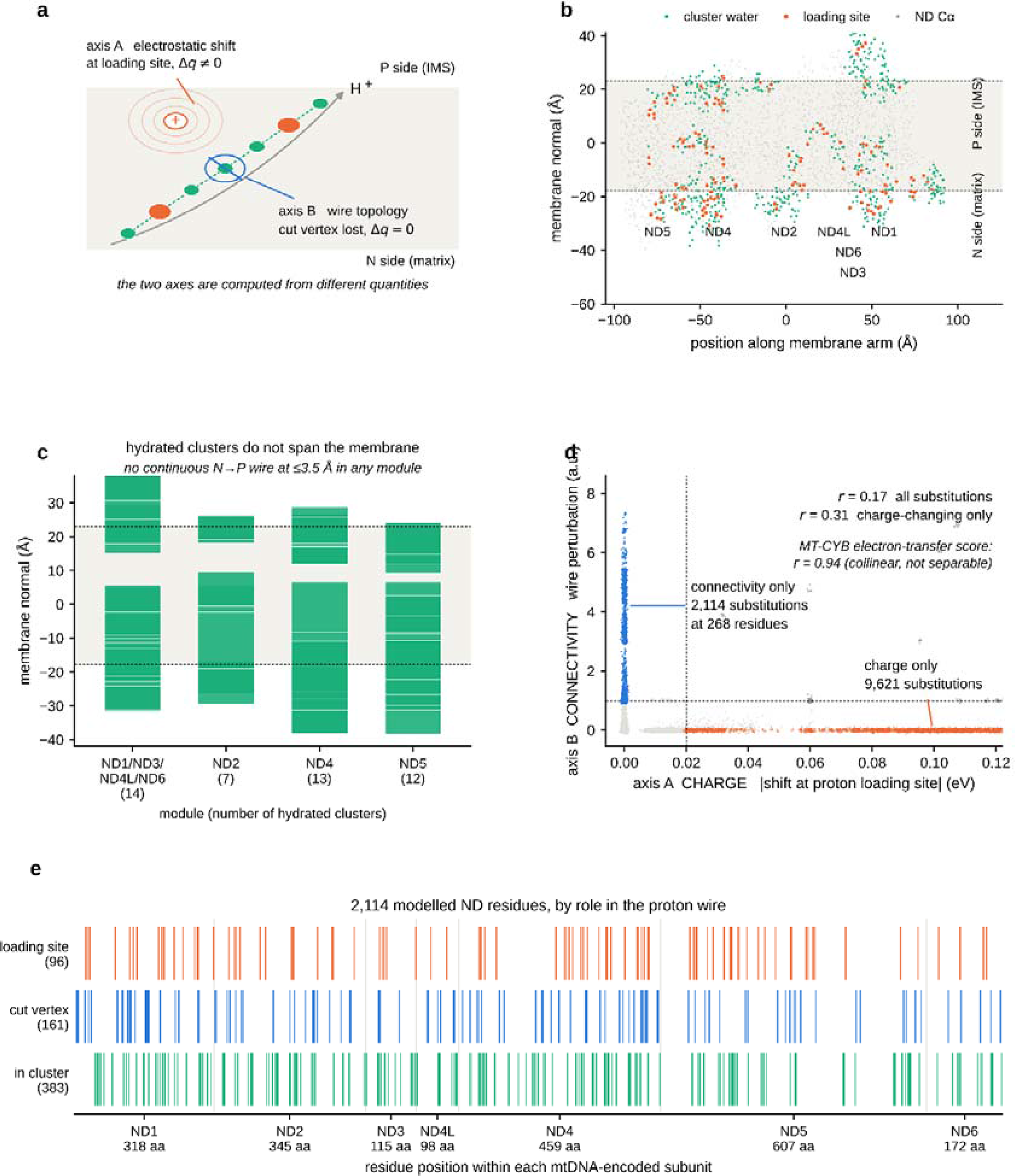
| A separable two-axis framework for mtDNA-encoded complex I variants. a Concept. A substitution at a residue lining a hydrated proton wire can perturb it in two independent ways: by shifting the electrostatic potential at a proton loading site without touching the wire (axis A, charge), or by removing a cut vertex of the hydrogen-bonded network without changing residue charge (axis B, connectivity). b Proton-wire architecture of the complex I membrane arm, from the cryo-EM structure of mouse complex I in the active state (PDB 8OM1, 2.39 Å), projected onto the membrane normal and the principal in-membrane axis. Grey, Cα atoms of the seven mtDNA-encoded ND subunits (n = 2,114 modelled residues); green, water molecules belonging to a major hydrated cluster; orange, proton loading sites. Dashed lines, bilayer boundaries derived from the deposited lipids and detergents (−17.9 to +23.0 Å). c Extent along the membrane normal of each major hydrated cluster, by module; the number of clusters is given in parentheses. No module carries a continuous hydrogen-bonded path from the N side to the P side at a donor–acceptor cutoff of 3.5 Å. d All 40,166 possible missense substitutions at the 2,114 modelled ND residues, scored on both axes. Pearson r = 0.17 across all substitutions and r = 0.31 among charge-changing substitutions (n = 13,307), against r = 0.94 for the collinear electron-transfer score of our cytochrome b analysis. Dashed lines, thresholds defining the off-diagonal classes (0.02 eV on axis A, 1.0 on axis B); points are jittered for display. e Classification of the 2,114 modelled ND residues by role in the proton wire, plotted along each subunit. Source data are provided as a Source Data file.

Nodes of the graph are the exchangeable side-chain heavy atoms of Ser, Thr, Tyr, Cys, Asp, Glu, Asn, Gln, His, Lys, Arg and Trp, together with ordered waters. Backbone amide and carbonyl atoms were excluded: they are not Grotthuss-competent, and including them makes the network continuous across the membrane at 2.85 Å in every module, an artefact of α-helical hydrogen bonding rather than a proton pathway. The restricted graph has 2,948 nodes (1,329 waters, 1,185 ND side-chain atoms, 434 from neighbouring subunits) and 2,006 edges at a donor–acceptor cutoff of 3.5 Å.

Connected components of this graph recover the known architecture. Clusters spanning 25–27 Å along the membrane normal sit in ND2, in ND4, and at the ND1–ND3–ND4L–ND6 junction that forms the E-channel (**Fig. 1b, c**). From these we defined 96 proton-loading sites (titratable residues in a major cluster), 111 wire cut vertices (residues whose removal disconnects their cluster) and 382 hydrated-cluster members in total (**Fig. 1e**). The classification is stable across cutoffs of 3.2, 3.5 and 4.0 Å, which place 240, 382 and 497 ND residues in a cluster.

No module carries a continuous hydrogen-bonded path from the N side to the P side at any cutoff up to 3.5 Å (**Fig. 1c**). The clusters are interrupted near the middle of the bilayer, as expected for gated, transiently hydrated channels. We therefore score path integrity rather than proton-transfer rates throughout.

Human complex I structures of comparable resolution exist (PDB 9TI4, 2.66 Å; 9I4I, 2.63 Å) but model one water molecule between them. Building the wire on human side chains alone collapses it to 292 edges and 48 clustered residues, against 2,006 edges and 382 residues for mouse. Superposing mouse onto human over 2,106 Cα atoms gives an RMSD of 1.65 Å — the fold is the same — but 2,042 of the 2,945 waters then clash with a human heavy atom below 2.2 Å, so transferred waters are not admissible. We therefore kept the mouse geometry and mapped the classification onto human codons by alignment. Identity ranges from 52.9% (ND6) to 77.0% (ND1), and wire residues are more conserved than the rest (75.9% versus 62.3%). All analyses below are restricted to the 1,367 positions at which the mouse and human residues are identical, retaining 78 of 96 loading sites and 85 of 111 cut vertices. The alignment, the indel positions and the wire residues that cannot be transferred are given in **Supplementary Fig. 4**, and the comparison of structural strategies in **Supplementary Fig. 5**.

Sequence identity is a proxy for the quantity that actually matters, which is whether a residue occupies the same position in the two structures, so we measured that directly. For each of the 2,104 mapped residues the Cα atoms within 10 Å were superposed locally onto their human counterparts, and two quantities recorded: the root-mean-square deviation of that local environment, and the distance the side-chain centroid moves once it is superposed. Local agreement is close — median Cα RMSD 0.46 Å and median side-chain deviation 0.79 Å — and 1,756 of 2,104 residues (83.5%) agree within 1.0 Å and 2.0 Å respectively, including 82 of 96 loading sites and 96 of 111 cut vertices (**Supplementary Fig. 7a**). Repeating the selection analysis on the 1,250 codons that are both structurally concordant and identical in sequence leaves every estimate within sampling error of the primary analysis: loading sites 0.47 in the germline and 1.37, 1.71 and 1.23 in the three tumour cohorts, all P ≤ 0.041 (**Supplementary Fig. 7b**). The cross-species transfer is not carrying the result.

### Charge and connectivity are separable properties of the wire

We scored all 40,166 possible missense substitutions at the 2,114 modelled residues on two axes: the screened-Coulomb shift imposed on proton-loading sites (charge), and the topological disruption of the hydrogen-bonded network (connectivity).

The axes are close to orthogonal (Pearson r = 0.17, Spearman 0.07; r = 0.31 among the 13,307 charge-changing substitutions). For comparison, the electron-transfer perturbation score in our cytochrome b analysis correlated with its electrostatic component at r = 0.94, which prevented any property-specific claim there. (14, 15) Both off-diagonal classes are well populated: 2,106 substitutions at 267 residues sever the wire without changing charge, and 9,628 at 2,022 residues shift a loading site without touching the wire (Fig. 1d). After the identity restriction these become 1,556 substitutions at 197 residues and 6,293 at 1,298, with correlations unchanged (0.17 and 0.29).

Residual coupling is geometric rather than definitional: wire residues lie closer to loading sites than other residues do (median 6.9 versus 11.8 Å), so distance to the nearest loading site is carried as a covariate throughout.

### The germline purges both charge-changing and wire-severing substitutions

Among 858 germline ND missense variants at the analysed positions (HelixMTdb), every wire class is depleted relative to each cohort’s own strand-aware trinucleotide spectrum: loading sites 0.51 (P = 9.2 × 10LL), cut vertices 0.67 (P = 7.4 × 10L³) and other cluster members 0.67 (P = 1.8 × 10L³), against 1.10 for the remaining ND residues (P = 0.012; **Fig. 2a**). The null model is calibrated. Synonymous ND variants show observed/expected ratios of 0.87–1.01 in the germline and deviate from expectation in no class or cohort (minimum P = 0.17; **Fig. 3a**).

**Fig. 2.**
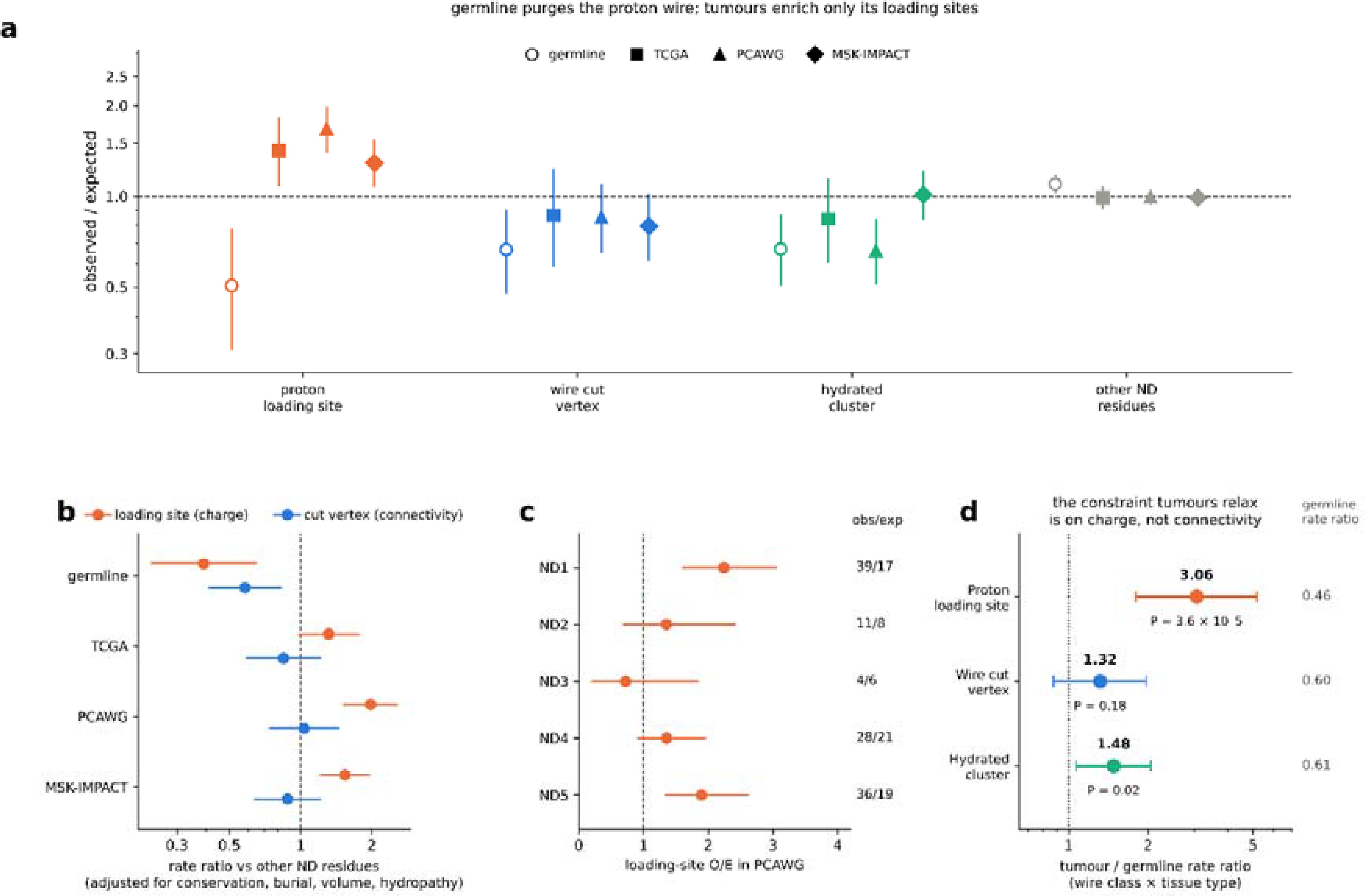
| Proton-loading sites are purged from the germline and enriched in tumours, whereas wire cut vertices are not. a Observed/expected ratios for ND missense variants by proton-wire class in one germline and three tumour cohorts, restricted to the 1,367 positions identical in mouse and human. Observed variants: HelixMTdb n = 858, TCGA n = 664, PCAWG n = 1,329, MSK-IMPACT n = 1,595. Expected counts derive from each cohort’s own strand-aware 192-channel trinucleotide spectrum, estimated from mtDNA variants outside ND coding sequence. Loading sites are depleted in the germline (O/E = 0.505, P = 9.2 × 10□□) and enriched in every tumour cohort (TCGA 1.417, P = 0.014; PCAWG 1.674, P = 1.1 × 10□□; MSK-IMPACT 1.291, P = 7.5 × 10□³), while cut vertices are depleted in the germline (0.665, P = 7.4 × 10□³) and never enriched in tumours (0.865, 0.852, 0.796; all P > 0.07). Error bars, 95% exact Poisson confidence intervals; two-sided exact Poisson tests. b Rate ratios against the reference class of other ND residues, from quasi-Poisson regression with the trinucleotide spectrum as offset, adjusted for PhastCons conservation, burial, side-chain volume change and hydropathy change (n = 6,825–6,999 substitutions per cohort). Points, rate ratios; error bars, 95% confidence intervals; dashed line, no difference from the reference class. c Observed and expected loading-site missense counts by ND subunit in PCAWG. The enrichment is carried by ND1 (39 observed against 17.4 expected) and ND5 (36 against 19.0), with weaker effects in ND4 (28 against 20.6) and ND2 (11 against 8.1) and none in ND3. d Wire class by tissue-type interaction, the direct test of relaxed constraint. For each wire class, the factor by which its germline rate ratio against other ND residues changes in tumours, from a single quasi-Poisson model of all four cohorts with the spectrum-weighted expectation as offset and standard errors clustered on the substitution; the germline rate ratio itself is printed under each point. Points, interaction ratios; error bars, 95% confidence intervals. The interaction is 3.06 for loading sites (95% CI 1.80–5.19, P = 3.6 × 10LL), 1.32 for cut vertices (0.88–1.97, P = 0.18) and 1.48 for other cluster members (1.07–2.05, P = 0.018). Dashed line, no interaction. Source data are provided as a Source Data file.

**Fig. 3.**
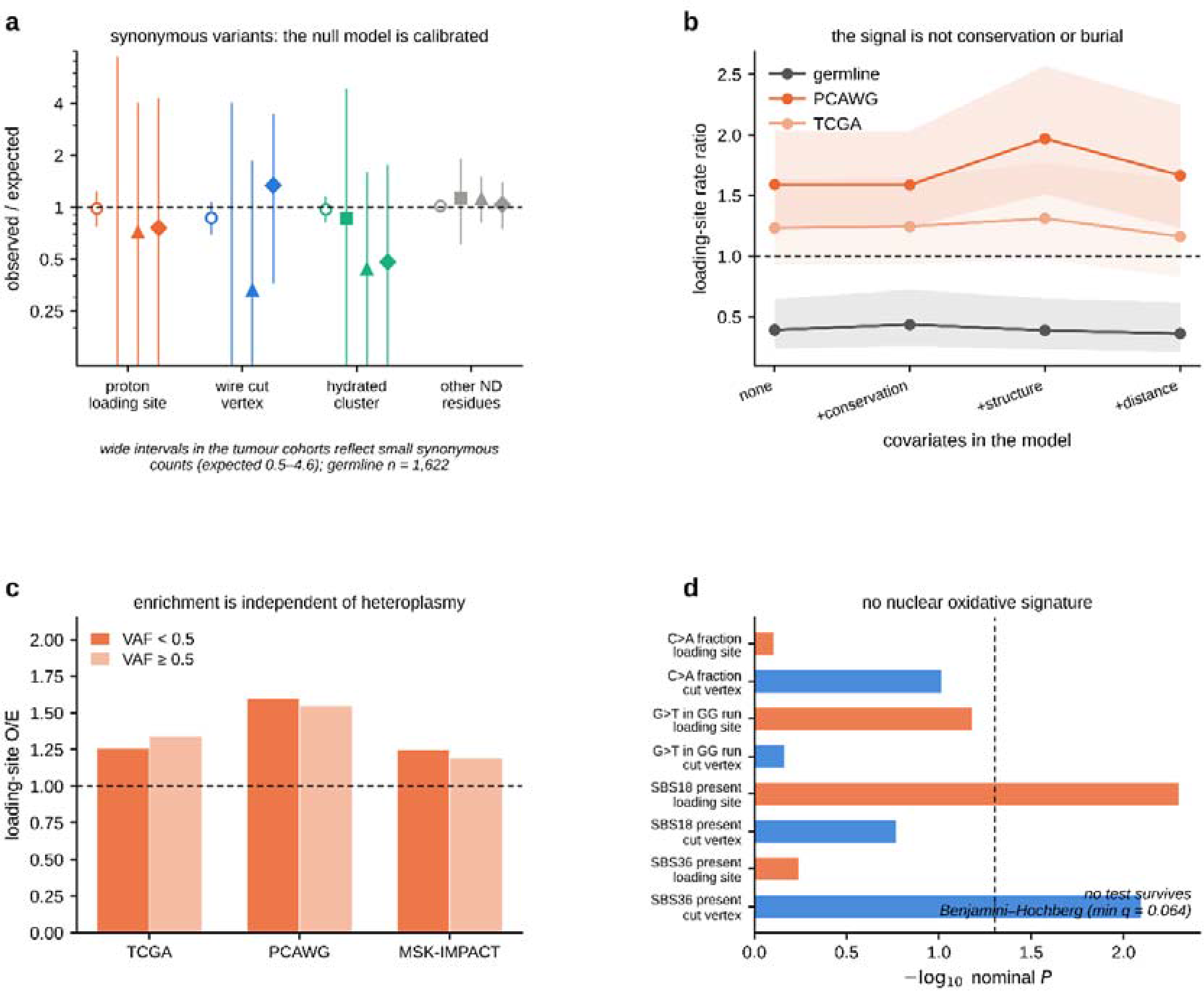
| The loading-site signal is robust to the null model, to covariates, to heteroplasmy and to nuclear mutational processes. a Observed/expected ratios for synonymous ND variants, which should be neutral, in the same four cohorts and classes as Fig. 2a. No class deviates from expectation in any cohort (minimum two-sided P = 0.17). Germline counts are large (n = 1,952 synonymous variants); tumour counts are small (expected 0.5–4.6), which accounts for the wide intervals. Error bars, 95% exact Poisson confidence intervals. b Loading-site rate ratio as covariates are added in a fixed ladder: unadjusted, plus PhastCons, plus burial, volume change and hydropathy change, plus distance to the nearest loading site. Shaded bands, 95% confidence intervals. The ratio is not attenuated by adjustment (PCAWG 1.59 → 1.97 → 1.67; germline 0.39 → 0.36). c Loading-site observed/expected ratio stratified at a variant allele fraction of 0.5 in each tumour cohort. Enrichment is present on both sides of the threshold, unlike the heteroplasmy-confined signal we reported for cytochrome b. d Nuclear mutational readouts in 2,207 TCGA tumours with both a nuclear substitution spectrum and mtDNA coverage, comparing tumours carrying loading-site (n = 51) or cut-vertex (n = 28) ND missense variants with tumours carrying no ND missense variant (n = 1,630). Bars show −logLL of the nominal two-sided P value; dashed line, P = 0.05. C>A fraction and G>T in guanine runs were tested by permutation within cancer type (10,000 permutations); SBS18 and SBS36 presence by Cochran–Mantel–Haenszel tests stratified by cancer type. No test survives Benjamini–Hochberg correction across the 16 comparisons (minimum q = 0.064). Source data are provided as a Source Data file.

### Tumours enrich loading-site substitutions but not wire cut vertices

In tumours the two classes separate. Loading sites are enriched in all three cohorts — 1.42 in TCGA (P = 0.014), 1.67 in PCAWG (P = 1.1 × 10LL) and 1.29 in MSK-IMPACT (P = 7.5 × 10L³) — while cut vertices are never enriched, sitting at or below expectation in each (0.87, 0.85 and 0.80; all P > 0.07; Fig. 2a). Other cluster members are unchanged in TCGA and MSK-IMPACT and depleted in PCAWG (0.66, P = 4.0 × 10LL). Residues outside the wire sit at expectation in every cohort (0.99 in every cohort). Within PCAWG the loading-site enrichment is carried by ND1 (observed 39, expected 17.4) and ND5 (36 versus 19.0), with weaker effects in ND4 (28 versus 20.6) and ND2 (11 versus 8.1) and none in ND3 (**Fig. 2c**). That MSK-IMPACT reproduces the signal is notable: our cytochrome b analysis failed to replicate in this cohort, which we attributed to its heteroplasmy distribution under-sampling the low allele fractions where that signal resided. (14, 15) The present signal is not confined to low heteroplasmy, which accounts for the difference. Comparing a germline ratio with a tumour ratio is not a test of the difference between them, so we fitted both arms in a single model. Counts for every substitution in all four cohorts were regressed on wire class, cohort and a wire class by tissue-type interaction, with the spectrum-weighted expectation as offset and standard errors clustered on the substitution. The loading-site rate ratio of 0.46 in the germline is multiplied by 3.06 in tumours (95% CI 1.80 to 5.19, P = 3.6 × 10^-5^). The same interaction is not significant for cut vertices (1.32, 95% CI 0.88 to 1.97, P = 0.18), and is weaker but present for the remaining cluster members (1.48, 95% CI 1.07 to 2.05, P = 0.018). Adding the conservation and structural covariates, or excluding germline polymorphisms from both arms, moves the loading-site interaction between 3.06 and 4.61 and never above P = 8.8 × 10^-4^ (**Fig. 2d and Supplementary Table 1**). The dissociation this paper reports is therefore a tested difference rather than a juxtaposition of two ratios. Cluster members that are neither loading sites nor cut vertices show a weaker relaxation of their own (interaction 1.48, P = 0.018), which adjusting for distance to the nearest loading site does not remove (1.69). The relaxation is therefore graded across the hydrated cluster rather than confined to its titratable sites, and only the cut vertices are spared.

### The dissociation survives conservation, burial and heteroplasmy

Adjusting for cross-species conservation (PhastCons), burial, side-chain volume change and hydropathy change does not attenuate the effect; if anything it sharpens it. Loading-site rate ratios are 0.39 (95% CI 0.23–0.65, P = 3.3 × 10LL) in the germline and 1.97 (1.51–2.57, P = 5.7 × 10LL) in PCAWG, 1.54 (1.20–1.96, P = 6.2 × 10LL) in MSK-IMPACT and 1.31 (0.97–1.77, P = 0.076) in TCGA. Cut vertices remain depleted in the germline (0.58, 0.41–0.83, P = 2.9 × 10L³) and null in all tumour cohorts (0.85, 1.03, 0.88; **Fig. 2b**). Adding distance to the nearest loading site, which is partly collinear with loading-site membership, gives a conservative 1.67 in PCAWG (**Fig. 3b**). The PhyloP column of the annotation release used is a missing-value sentinel for all but 0.8% of entries and could not be used; conservation adjustment therefore rests on PhastCons alone. Loading-site enrichment is present on both sides of a heteroplasmy threshold of 0.5 — 1.26 and 1.34 in TCGA (1.34 and 1.27 after purity correction), 1.60 and 1.55 in PCAWG, 1.25 and 1.19 in MSK-IMPACT (**Fig. 3c**) — and the heteroplasmy of loading-site missense variants is indistinguishable from that of truncating ND variants (TCGA median 0.607 versus 0.609, P = 0.52). This differs from the heteroplasmy-confined pattern we reported for cytochrome b, where enrichment collapsed above 0.5. (14, 15)

### Recurrence without clonal advantage

Cross-sectional allele fractions cannot separate tolerance from positive selection, but the distribution of calls over sites bears on it. Under tolerance, substitutions accumulate wherever mutation puts them; under selection, particular substitutions should recur across patients more often than mutational opportunity allows. Holding the number of calls fixed and drawing them over the eligible sites with the spectrum weights (10,000 multinomial draws), loading sites carry more recurrent substitutions than expected in PCAWG (32 sites seen in two or more patients against 16.9 expected, P = 1 × 10^-4^) and in TCGA (13 against 7.3, P = 0.024), but not in MSK-IMPACT (25 against 22.9, P = 0.32). The excess is not specific to the class in TCGA, where residues outside the wire show one as well (115 against 89.3, P = 3 × 10^-4^); in PCAWG it is (244 against 237.4, P = 0.26). The most recurrent single substitution is m.3380G>A, ND1 R25Q, in 46 patients across all three cohorts, followed by ND5 D179N, R161Q, R262H and E145K in 10 to 12 each (**Supplementary Table 2**).

A variant under positive selection within a tumour should also reach fixation more often than a tolerated one. It does not. Loading-site missense variants are no more clonal than other ND missense variants in any cohort (median allele fraction 0.61 versus 0.55 in TCGA, 0.04 versus 0.04 in PCAWG, 0.49 versus 0.50 in MSK-IMPACT; all P ≥ 0.23), and the proportion reaching an allele fraction of 0.9 is no higher — lower, in MSK-IMPACT, at 3.2% against 8.7% (P = 0.039). Truncating ND variants, by contrast, are more clonal than missense variants in TCGA (P = 0.015). Recurrence without a clonal advantage is what tolerance at mutationally favoured positions produces, and we therefore do not claim within-tumour positive selection. One substitution is the exception: m.3380G>A reaches an allele fraction above 0.9 in 7 of its 11 TCGA tumours, against 127 of 653 ND missense calls overall (P = 0.0019). Class-level tolerance and variant-level selection are not mutually exclusive, and this variant behaves as the second. The classification depends on one free parameter, the donor–acceptor distance at which two polar groups are counted as hydrogen bonded. Repeating the whole analysis at 3.2 Å and 4.0 Å leaves the loading-site result intact at 4.0 Å (germline 0.46, TCGA 1.34, PCAWG 1.52, MSK-IMPACT 1.21; all P ≤ 0.012) but not at 3.2 Å, where only 32 loading sites survive and the comparison loses power. Cut vertices are never enriched at any cutoff; at 4.0 Å they become depleted in tumours (0.68, 0.64 and 0.49) rather than neutral. The dissociation between charge and connectivity therefore holds throughout, while the precise behaviour of the connectivity class does not: what is robust is that tumours do not accumulate wire-severing substitutions, not that they are indifferent to them (**Supplementary Fig. 1**). Two further filters were tested by removal. Without the restriction to positions identical in mouse and human, loading sites remain depleted in the germline (0.64) and enriched in every tumour cohort (1.40, 1.57, 1.27; all P ≤ 0.014). Without the germline-polymorphism filter the somatic estimates are essentially unchanged (1.34, 1.62, 1.24). Neither filter generates the signal (**Supplementary Figs 2, 3**).

### Coverage, cancer type and subunit do not produce the signal

A position sequenced in more samples has more opportunity to be called mutated, and the wire classes are not spread evenly over the genes. The per-base coverage matrices distributed with the calls settle this directly. Loading-site codons are covered slightly less often than ND positions outside the wire, not more: mean coverage 0.460 against 0.474 in TCGA (P = 5.3 × 10^-4^) and 0.123 against 0.144 in MSK-IMPACT (P = 1.2 × 10^-9^). Entering log coverage as a covariate accordingly raises rather than lowers the loading-site rate ratio, from 1.43 to 1.50 in TCGA and from 1.30 to 1.54 in MSK-IMPACT (**Supplementary Fig. 8a**); PCAWG is whole-genome and uniformly covered. Coverage works against the enrichment reported here, not for it. Nor is the enrichment produced by the few lineages that carry most mitochondrial mutations. Scoring every cancer type with at least twenty calls against the same null puts 5 of 8 types above one in TCGA, 12 of 16 in PCAWG and 13 of 21 in MSK-IMPACT, with no detectable heterogeneity in any cohort (Cochran’s Q, P = 0.38, 0.55 and 0.62). Removing each type in turn moves the pooled ratio between 1.20 and 1.57 in TCGA, 1.46 and 1.75 in PCAWG, and 1.16 and 1.30 in MSK-IMPACT, so no single lineage carries it (**Supplementary Fig. 8b**).

Removing each ND gene in turn leaves the germline depletion (0.43 to 0.57) and the tumour enrichment (TCGA 1.11 to 1.52, PCAWG 1.50 to 1.79, MSK-IMPACT 1.21 to 1.36) in place, so no single subunit produces the effect either (Supplementary Fig. 8c). ND1 is nonetheless the largest contributor: without it the enrichment loses significance in TCGA (1.11, P = 0.61) and in MSK-IMPACT (1.21, P = 0.12) while holding in PCAWG (1.50, P = 4.3 × 10^-4^). Scored on its own, ND6 runs the other way in tumours (0.34 to 0.40); it is also the subunit for which the cross-species transfer is weakest. The two free parameters of axis A do not carry the separability. The effective dielectric constant scales every score by one factor and so cannot change a correlation or a ranking; across distance floors of 2, 3 and 5 Å the correlation between the axes runs from 0.20 to 0.12, and at a seventy-fifth-percentile threshold the depletion of loading-site charge change among common germline variants is 6.7% against 25% at every one of the nine parameter settings (Supplementary Fig. 8d). Finally, of the 48 primary tests reported here, eight survive a 5% false-discovery rate over the whole set: loading-site depletion in the germline and enrichment in PCAWG and MSK-IMPACT, the tissue-type interaction, the PCAWG recurrence excess, cluster-member depletion in the germline and in PCAWG, and cut-vertex depletion in the germline. TCGA loading-site enrichment survives correction within its own family (q = 0.023) but not over the whole paper (q = 0.073), which is the limitation of that cohort’s sample size rather than of the effect (**Supplementary Table 3**).

### The tumour signal is concentrated at the quinone cavity

If relaxed constraint acted on proton translocation as such, it should be spread along the path. It is not. Splitting the loading sites by position in the structure, those of ND1 that lie within 15 Å of the quinone ligand modelled in 8OM1 are enriched 4.84-fold in TCGA (P = 1.5 × 10^-8^), 3.85-fold in PCAWG (P = 2.4 × 10^-9^) and 2.86-fold in MSK-IMPACT (P = 8.8 × 10^-7^). ND1 loading sites further from the cavity are not enriched in any cohort (0.64 to 1.17, all P ≥ 0.62), and loading sites outside ND1 are enriched only in PCAWG (1.47, P = 7.4 × 10^-4^; 1.07 in TCGA and in MSK-IMPACT). The germline behaves differently again: depletion there is carried by the loading sites outside ND1 (0.40, P = 1.4 × 10^-4^), the positions the tumours leave alone. Constraint in the population is distributed along the wire; its relaxation in tumours is local (**Fig. 4a, b**). One substitution carries much of this. m.3380G>A, which replaces ND1 Arg25 with glutamine, is the most recurrent proton-wire substitution in the study, seen in 46 patients across all three cohorts and reaching a median allele fraction of 0.95 in TCGA. It is absent from HelixMTdb, defines no branch of PhyloTree, and is listed in MITOMAP as a reported variant in MELAS. Excluding it, the quinone-proximal enrichment falls but persists in TCGA (2.35, P = 0.025) and PCAWG (2.07, P = 0.012) and disappears in MSK-IMPACT (0.84, P = 0.75), so the signal in that cohort rests on this one substitution (**Fig. 4c**). Coverage does not explain the position: the quinone-proximal codons are covered slightly less than the rest of ND1 (0.465 against 0.482 in TCGA). Five further recurrent substitutions lie in the same region, among them ND1 Glu204Lys and Asp199Asn, both within 13 Å of the ligand.

**Fig. 4.**
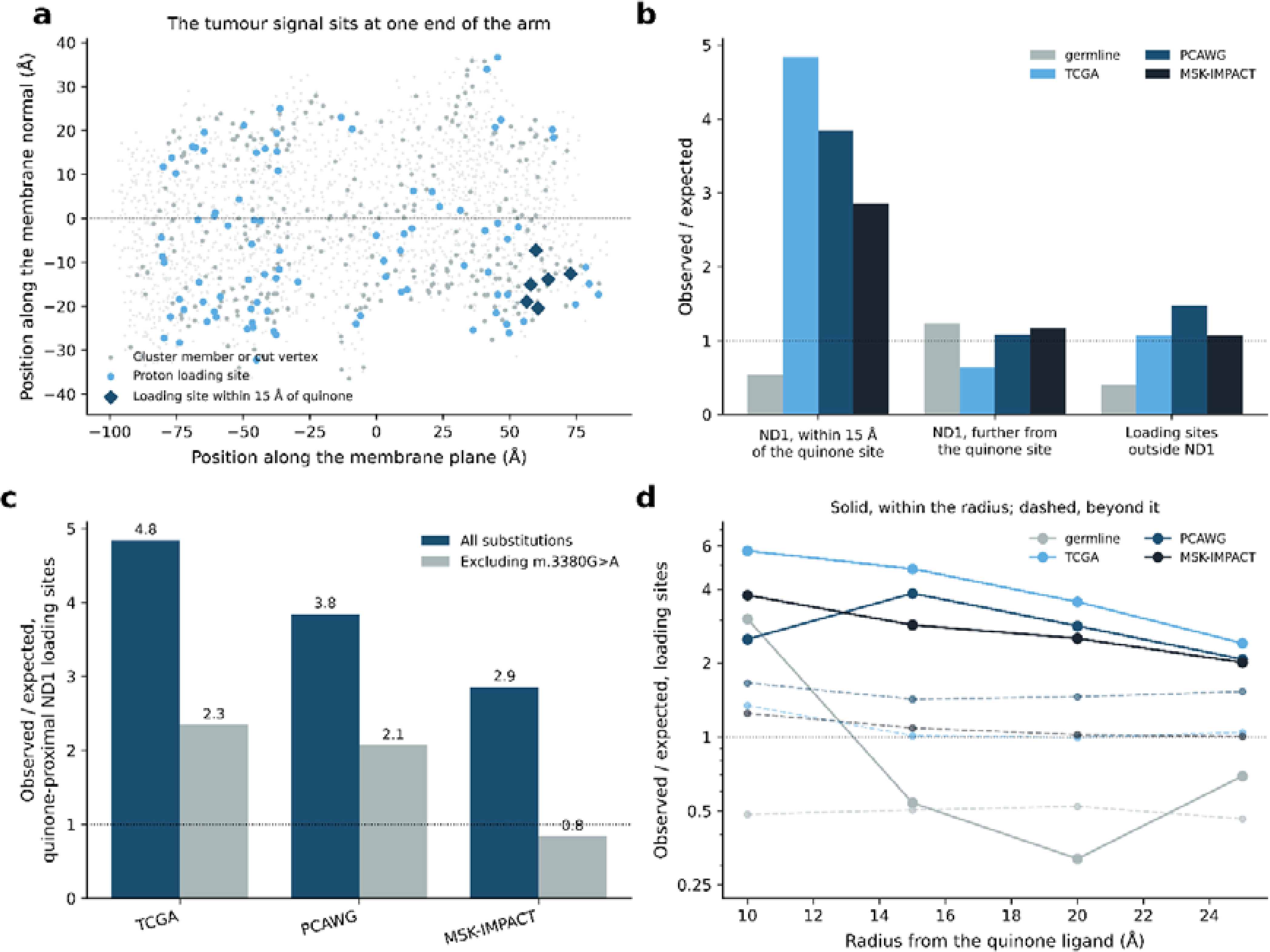
| The tumour signal is concentrated at the quinone cavity of ND1. a All modelled ND residues in the membrane frame, with loading sites within 15 Å of the quinone ligand modelled in PDB 8OM1 marked as diamonds. b Observed/expected for loading-site missense substitutions split by position relative to the quinone cavity — ND1 within 15 Å, ND1 beyond 15 Å, and loading sites outside ND1 — in the germline and the three tumour cohorts. Quinone-proximal ND1 loading sites are enriched 4.84-fold in TCGA (P = 1.5 × 10□□), 3.85-fold in PCAWG (P = 2.4 × 10□□) and 2.86-fold in MSK-IMPACT (P = 8.8 × 10□□), whereas ND1 loading sites further from the cavity are not enriched in any cohort (0.64 to 1.17, all P ≥ 0.62). Germline depletion is carried instead by the loading sites outside ND1 (0.40, P = 1.4 × 10□□). c The quinone-proximal estimate with and without the single most recurrent substitution, m.3380G>A (ND1 Arg25Gln). Without it the enrichment persists in TCGA (2.35, P = 0.025) and PCAWG (2.07, P = 0.012) and disappears in MSK-IMPACT (0.84, P = 0.75). d Observed/expected for loading-site missense substitutions inside (solid) and outside (dashed) a radius from the quinone ligand, as the radius is swept from 10 to 25 Å; the germline shows no such gradient. Error bars, 95% exact Poisson confidence intervals; two-sided exact Poisson tests. Source data are provided as a Source Data file.

Because so much rests on this one substitution, we checked it against the matched normals. In TCGA all eleven calls have a matched normal at a median depth of 122 reads and not one alt read in any of them; in PCAWG all fourteen have a normal at a median depth of 2,260 reads and, again, no alt reads. That is not the background: among the other wire-class calls in PCAWG, where the normals are deep enough to detect low-level inherited heteroplasmy, 193 of 236 show at least one alt read in the normal, against 0 of 14 for m.3380G>A (P = 2.6 × 10^-10^). The substitution is not annotated in a homopolymer, and the tumour calls are deep (median 357 reads, 274 of them alt). It also reaches fixation far more often than its peers: 7 of 11 TCGA calls exceed an allele fraction of 0.9, against 20 of 117 other wire-class calls (P = 0.0017) and 127 of 653 ND missense calls overall (P = 0.0019). m.3380G>A therefore arises somatically and expands clonally. The MSK-IMPACT normals are too thin for this test (median 4 reads), so the same cannot be shown there. Away from the quinone cavity, the graph does recover the arrangement the structural literature describes. Eight charged loading sites lie within 6 Å of the bilayer midplane in the antiporter-like subunits, forming a chain that runs 37 Å along the membrane plane with a median neighbour spacing of 4.8 Å: ND5 His332, Lys223, Glu145, Asp179 and Arg176, and ND4 Glu378, His293 and His319. Of the 23 charged residues in that slab, 15 sit in a hydrated cluster (Supplementary Fig. 9d). The classification also picks out both glutamates of ND4L, Glu34 and Glu70, which are the only two the subunit has; both recur in tumours. In absolute terms, 2.3% of TCGA patients, 5.5% of PCAWG patients and 2.3% of MSK-IMPACT patients carry a loading-site missense variant, and 0.6% to 1.4% carry one of the quinone-proximal ND1 substitutions.

The 15 Å radius is ours, so we removed it. Sweeping the radius from 10 to 25 Å leaves loading sites inside it enriched in every tumour cohort (TCGA 5.73 to 2.41, PCAWG 2.50 to 2.07, MSK-IMPACT 3.78 to 2.01) and those outside it close to expectation (0.99 to 1.66), with the germline showing no such gradient (**Fig. 4d**). Treating distance as a continuous variable removes the threshold altogether. Regressing the counts on proximity to the cavity, with the spectrum-weighted expectation as offset, gives a rate ratio per e-fold decrease in distance of 2.81 in TCGA, 1.43 in PCAWG and 1.55 in MSK-IMPACT after adjustment for conservation, burial, volume change and hydropathy change (all P ≤ 0.002). Because distance to the cavity is largely a property of the subunit, we added subunit fixed effects; the gradient does not weaken but strengthens, to 4.57, 3.12 and 3.20 (all P ≤ 0.003), and within ND1 alone it is 10.06 in TCGA (P = 0.004) and 4.05 in PCAWG (P = 0.039), with MSK-IMPACT underpowered at that scale (3.99, P = 0.31). Proximity to the quinone cavity therefore predicts tumour enrichment within a subunit, not only between subunits (**Supplementary Fig. 9a**).

The gradient is specific to the tumours. Fitted on its own with subunit effects, the germline shows none (0.68, P = 0.54). Fitting both arms together and allowing the subunit effects to differ between them, the tumour gradient exceeds the germline gradient 7.9-fold (95% CI 1.3 to 47.6, P = 0.024; 10.1-fold with the covariates, P = 0.019). The confidence interval is wide, as it must be with twenty germline calls at these positions, but the direction is the one the enrichment implies (**Supplementary Fig. 9b**). A blunter statistic, the mean distance of all observed calls to the cavity, is shifted toward it in TCGA (P = 0.003) and MSK-IMPACT (P = 0.043) but not in PCAWG (P = 0.36), where the calls far from the cavity dominate the mean. The other half of the dissociation is a null, which deserves a bound rather than a P value alone. Cut vertices give observed/expected 0.865 in TCGA, 0.852 in PCAWG and 0.796 in MSK-IMPACT, with upper confidence limits of 1.24, 1.10 and 1.02. The data therefore exclude enrichment above those values, and the expected counts are large enough to have detected a true ratio of 1.55, 1.38 and 1.36 four times in five (**Supplementary Fig. 9c**). An effect the size of the loading-site enrichment would not have been missed.

### No detectable effect on nuclear oxidative mutagenesis

Increased proton leak or altered quinone chemistry could raise mitochondrial superoxide and leave an oxidative mark on the nuclear genome. We tested this in 2,207 TCGA tumours with both a nuclear substitution spectrum and mtDNA coverage, comparing those carrying loading-site (n = 51) or cut-vertex (n = 28) ND missense variants with those carrying none (n = 1,630). Single-base substitution profiles were fitted to COSMIC v3.4 exome signatures (8,058 tumours, median reconstruction cosine 0.91).

None of four readouts — C>A fraction, G>T at guanines within GG/GGG runs, and the presence of SBS18 or SBS36 — survives multiple-testing correction across the sixteen comparisons (minimum q = 0.064; Fig. 3d). The strongest nominal associations, SBS18 with loading-site variants (odds ratio 2.50, P = 0.005) and SBS36 with cut-vertex variants (3.08, P = 0.008), rest on 51 and 28 tumours respectively and should not be interpreted. The full set of readouts, with Benjamini–Hochberg q values, is shown in **Supplementary Fig. 6**

### Mitochondrial disease variants do not occupy a distinct niche

We asked whether pathogenic ND variants concentrate on the axis that tumours leave alone. They do not. Among 21 MITOMAP-confirmed pathogenic substitutions within the analysed set, no class is enriched (loading sites 1.03, cut vertices 2.04, P = 0.37), but expected counts are 1.0–2.0 per class, so the comparison is underpowered rather than negative. A broader reported-pathogenic set (n = 115) trends toward loading sites (1.88, P = 0.089) — the same direction as tumours, not the opposite one (**Supplementary Fig. 10a**). Disease and tumour variants also overlap strongly: 49 of 115 reported-pathogenic substitutions are observed in at least one tumour cohort, against 10.2% of all other substitutions (odds ratio 6.4, P = 9.7 × 10L¹L; **Supplementary Fig. 10c**). One contrast at the variant level is sharp. Substitutions common in the germline — carried by more than ten HelixMTdb individuals — shift a loading site by at least 0.02 eV in only 6.7% of cases, against 24.7% of all possible substitutions and 24.3% of tumour-observed ones (**Supplementary Fig. 10b**). What the population tolerates at high frequency avoids loading-site charge change specifically.

### Substitutions that fixed over evolutionary time conserve charge

The germline cohort reports depletion in the present population. Variants that define branches of the human mitochondrial phylogeny report a longer interval: substitutions that reached fixation along a lineage and have been carried since. Of 4,950 branch-defining substitutions in PhyloTree build 17, 173 are ND missense substitutions at the analysed positions. Loading sites carry five of them against 8.0 expected (0.63) and cut vertices eight against 12.1 (0.66) — the same direction as the germline, though neither reaches significance at this sample size (P = 0.39 and 0.29) — while the synonymous set is unbiased (0.93 to 1.27, all P ≥ 0.12). Placed beside the cohorts, the loading-site ratio runs 0.63 on the phylogeny and 0.51 in the present germline against 1.29 to 1.67 in tumours (**Supplementary Fig. 7c**).

What the five fixed substitutions are is more informative than how many. All five exchange one titratable group for another: ND1 Y30H on ten branches, ND5 Y159H on five, ND4 Y409H on four, ND5 H67Y on two and ND1 Y30C on one. The electrostatic shift they impose at a loading site is accordingly small, 0.048 eV on average, against 0.827 eV for the loading-site substitutions that recur in four or more tumours and 0.431 eV across all possible loading-site substitutions (P = 6.3 × 10^-4^, **Supplementary Fig. 7d**). What evolution has avoided at these positions is not the position but the charge change. Tumours recur at the same positions with the unit-charge substitutions — arginine to glutamine, glutamate to lysine, aspartate to asparagine — that the phylogeny does not carry.

## Discussion

We find that selection on the proton-translocating machinery of complex I distinguishes two physically separable properties of the same hydrated network. (21) Substitutions that change charge at a proton-loading site and substitutions that sever the hydrogen-bonded wire are both purged from the human germline. In tumours the two part company: loading-site substitutions are enriched in three independent cohorts, while wire-severing substitutions are never enriched in any of them. (11) The dissociation survives adjustment for sequence conservation, burial and side-chain physicochemistry, and is not an artefact of the mutational null, which is calibrated on synonymous variants. (11) Fitted in one model, the germline-to-tumour change at loading sites is a 3.06-fold relaxation, while at cut vertices it is not significant.

The distinction is interpretable in terms of what each class of substitution does to the enzyme. (21) A charge change near a loading site shifts the protonation free energy of a titratable group without removing it, which can alter coupling efficiency or pumping stoichiometry while leaving the complex assembled and catalytically active. Severing a wire is closer to loss of function. That tumours do not accumulate the second class may reflect the availability of a cheaper route to the same end: truncating variants are enriched at least two-fold in complex I relative to other respiratory complexes and reach high heteroplasmy, with a substantial share arising at homopolymer indel hotspots. If complex I loss is what is selected, a frameshift achieves it more reliably than a missense substitution that happens to remove a bridging side chain. (11) On this reading, the missense signal we detect is not about losing complex I at all, but about tuning the electrostatics of proton loading in complexes that remain intact.

Why charge, and why there? The localisation argues against the interpretation we first entertained, that a charge change degrades pumping efficiency while leaving catalysis intact. Mutagenesis of the charged residues of the membrane arm does not produce that phenotype: in Paracoccus denitrificans, neutralising the central-axis residues of ND4 impairs NADH: quinone oxidoreduction, and every variant that remains active still pumps four protons per NADH, with no uncoupling and no evidence of slip. (11) In Escherichia coli the equivalent lysines of ND2 are required for electron transfer and proton pumping alike. Charge changes in the membrane arm therefore impair the enzyme as a whole rather than decouple it, and it is precisely there that we find no tumour enrichment. The signal sits instead at the quinone cavity, where ND1 substitutions associated with LHON and MELAS are known to act on ubiquinone binding, and where a charge change alters the chemistry and the first protonation events of quinone reduction rather than the stoichiometry of the pump. (22–24)

What the data establish is therefore narrower than a general loosening of constraint on complex I. (25) The relaxation is not a property of the proton path along its length but of one end of it. (11) This differs from what we found for cytochrome b, in a way that is internally consistent. (26) There, enrichment of charge-changing variants near the haem centres was confined below a heteroplasmy threshold of 0.5 and collapsed above it, which we read as relaxed purifying selection with wild-type mtDNA buffering the defect. (27) Here the enrichment is present on both sides of that threshold, and the heteroplasmy distribution of loading-site variants is indistinguishable from that of truncating variants, so buffering does not describe this pattern. Whether tumours gain from the relaxation, tolerate it, or merely fail to purge it, our data cannot say: loading-site substitutions recur across patients more than mutational opportunity allows, but reach fixation no more often than other missense substitutions. (25) We make no claim of a fitness benefit. The exception is m.3380G>A, verified somatic against deep matched normals and fixed more often than its peers; whatever holds for the class, that substitution behaves as if it is selected within the tumour.

One result sits outside this account. In PCAWG, hydrated-cluster members that are neither loading sites nor cut vertices are depleted rather than neutral (0.66, P = 4.0 × 10LL), an effect absent in the other two cohorts. We report it without an explanation; it may reflect residual structure in the PCAWG spectrum or a genuine constraint on the cluster interior that our two axes do not capture. (28) Stratifying by allele fraction does not localise it, so it is not an artefact of low-allele-fraction calling.

Our initial expectation that mitochondrial disease variants would occupy the axis tumours leave alone was not supported. (29) The confirmed MITOMAP set is too small to place, the broader reported set trends toward loading sites — the same direction as tumours — and the two sets overlap strongly. (29) Rather than distinct niches, this points to a shared set of positions at which the protein is sensitive, reached by germline transmission in one case and somatic mutation in the other. Purifying selection is nonetheless visible at the level of individual variants: substitutions that have become common in the population avoid loading-site charge change specifically, and those that fixed on branches of the human mtDNA tree exchange one titratable group for another. What has been avoided over evolutionary time is the charge change, not the position.

The nuclear genomes of the same tumours carry no oxidative mark: no readout among C>A burden, G>T at guanine runs, SBS18 and SBS36 survives multiple-testing correction. (30, 31) A charge change at the quinone cavity need not raise superoxide production, and the negative result is consistent with that; but it is also what limited power would produce, and it mirrors our negative result for complex III Q^(o)-site variants. (4, 30) It does not exclude localised mitochondrial reactive oxygen species signalling, and the phenotype most relevant to a proton wire — membrane potential and ΔpH — is not something a nuclear mutational signature can report. (32)

Several limitations warrant consideration. The wire geometry comes from a mouse structure, because no human structure of comparable resolution models hydration; we verified that human side chains alone cannot support the analysis and that the local environment of a mapped residue agrees between the species, but the transfer still costs us a quarter of the wire residues, including ND6 positions that carry LHON variants. (3) The graph is built from a single static structure and its ordered waters, with no conformational ensemble, so we score path integrity and make no rate predictions. (33, 34) Axis B is ordinal rather than an energy and its steric term is heuristic, but the residue classes used for the selection analyses are derived from the graph alone, so that heuristic does not reach the main result. The hydrogen-bond cutoff, by contrast, does affect the connectivity class. Conservation adjustment rests on PhastCons alone, because the PhyloP column of the annotation release used is a missing-value sentinel; mitochondrial local constraint scores would strengthen the tumour arm, though they cannot be applied to the germline arm without circularity. TCGA does not reach significance on its own, and the enrichment survives removal of any single subunit except that TCGA and MSK-IMPACT depend on ND1 for it. Two further limitations follow from the localisation. The structure is the active state, and complex I interconverts with a deactive state whose quinone-cavity loops and hydration differ, so the wire we build describes one conformer of a cycling enzyme. (33) And the quinone-proximal enrichment depends substantially on m.3380G>A, whose somatic status cannot be confirmed in MSK-IMPACT, the cohort where it is most frequent, because those normals reach a median of four reads. Nothing here is functionally validated.

The decisive experiment is available. Charge and connectivity make different predictions for the kinetic isotope effect: a classical electrostatic shift at a loading s1ite does not change the H/D ratio of proton pumping, whereas a change in the geometry of the transfer barrier does. (35, 36) Measuring pumping stoichiometry in HLO and DLO in cybrid lines carrying variants selected from the off-diagonal classes would separate the two mechanisms directly, and would test whether the property tumours relax is electrostatic in the ordinary sense or something more. (37)

Finally, the scope of this result is a single enzyme. Complex I is not the only proton-handling machine encoded by mitochondrial DNA: COX1 to COX3 carry the D- and K-channels of complex IV, and ATP6 the half-channels of the ATP synthase, where the essential charges are catalytically indispensable rather than efficiency-determining and the account proposed above therefore predicts no relaxation at all. (38, 39) Whether the dissociation reported here is particular to complex I or a general feature of selection on proton-handling machinery is an open question, directly testable with the same data and the same two axes, and one we do not settle.

## Methods

### Structure and construction of the proton-wire graph

#### We used the cryo-EM structure of mouse heart respiratory complex

I in the active state (PDB 8OM1, 2.39 Å), which models 2,945 ordered water molecules. (3) All seven mtDNA-encoded subunits are resolved: ND1 (318 residues), ND2 (345), ND3 (115), ND4 (459), ND4L (98), ND5 (607) and ND6 (172), giving 2,114 modelled residues in total. Alternative conformations were removed and the first conformer retained. The membrane frame was defined from the deposited lipid and detergent molecules (3PE, CDL, PC1, LMT, MYR): the membrane normal is the least-variance principal axis of their atoms and the in-membrane axis the greatest, placing the bilayer between −17.9 Å and +23.0 Å on the normal. The matrix (N) side was identified as the side carrying the FMN cofactor. Nodes of the proton-wire graph are the Grotthuss-competent heavy atoms: the hydroxyl, carboxyl, amide, imidazole, amine, guanidinium, thiol and indole atoms of Ser, Thr, Tyr, Cys, Asp, Glu, Asn, Gln, His, Lys, Arg and Trp side chains, together with the oxygen of every modelled water within 12 Å of an ND subunit, and the equivalent side-chain atoms of nuclear-encoded subunits lining the ND surface within 6 Å. (18) Backbone amide and carbonyl atoms were excluded. (20) They are not Grotthuss-competent — the amide proton does not exchange on a catalytic timescale —and including them saturates the network: with backbone atoms the graph is continuous across the membrane at 2.85 Å in every module, which is an artefact of α-helical hydrogen bonding rather than a proton pathway. Restricting to exchangeable side chains and waters gave 2,948 nodes (1,329 waters, 1,185 ND side-chain atoms, 434 from neighbouring subunits). Edges connect node pairs at heavy-atom distances between 2.2 Å and 3.5 Å, excluding pairs within one residue. Because the cryo-EM model carries no hydrogens, a distance-only criterion was used, as is standard for structures without modelled protons. Connected components of four or more nodes were taken as hydrated clusters. (40) We report a sensitivity sweep over cutoffs of 3.2, 3.5 and 4.0 Å, which places 240, 382 and 497 ND residues respectively in a cluster; 3.5 Å was used throughout. Three residue classes were defined from this graph, in a strict hierarchy. Proton loading sites are titratable residues (Asp, Glu, His, Lys, Arg, Tyr, Cys) belonging to a major hydrated cluster, where major means at least 12 nodes or a span of at least 10 Å along the membrane normal (96 residues). (6) Wire cut vertices are residues whose side-chain nodes are articulation points of their cluster, so that deleting them disconnects it (111 residues). Hydrated-cluster members are the remaining residues contributing nodes to a cluster of at least four (382 residues in total across the three classes). All other modelled ND residues form the reference class.

No module carries a continuous hydrogen-bonded path from the N side to the P side at any cutoff up to 3.5 Å. The hydrated clusters span 25–27 Å along the membrane normal and are interrupted near the middle of the bilayer, consistent with the gated, transiently hydrated channels described for complex I. We therefore make no claim about proton-transfer rates and score only path integrity.

### The two axes

Every possible missense substitution at every modelled ND residue (40,166 substitutions over 2,114 residues) was scored on two axes designed to be statistically separable. (41, 42)

#### Axis A, charge

The screened Coulomb shift that a substitution’s net charge change imposes on each proton loading site, ΔE = 14.4·Δq / (ε·r) eV, with r the distance from the substituted side-chain centroid to the loading-site centroid, floored at 3 Å, and ε = 8. (43, 44) Net charge used Asp and Glu = −1, Lys and Arg = +1, His = +0.1 at pH 7.4. (44) Axis A is the largest absolute shift over all loading sites. This is the same functional form used for haem redox-potential shifts in our previous analysis of cytochrome b.

#### Axis B, connectivity

The perturbation a substitution imposes on wire topology, combining two terms. (45) A substitution that removes a Grotthuss-competent side chain from a hydrated cluster contributes 1 + (fragments created on deletion) + 4·(weighted betweenness within its cluster) + 1 if the residue is a cut vertex. (45) A substitution that increases side-chain volume by more than 20 Å³ at a residue lining cluster waters contributes a steric term proportional to the excess volume and the number of cluster waters within 5 Å. (46) Axis B is ordinal rather than an energy, and we report it in arbitrary units. Axis B is used only for the separability diagnostic; the residue classes entering the selection analyses are defined from the graph alone.

The axes are close to orthogonal. Across all possible substitutions the Pearson correlation between them is 0.17 (Spearman 0.07); restricted to charge-changing substitutions, the more demanding test, it is 0.31. For comparison, the electron-transfer perturbation score in our cytochrome b analysis correlated with its electrostatic component at r = 0.94, which prevented any tunnelling-specific claim there. The separation arises by construction: axis B is computed from graph topology and side-chain geometry, not from the electrostatic term, so a charge-neutral substitution that severs a wire scores on B alone, while a distant charge change scores on A alone.

The off-diagonal cells are well populated. Taking thresholds of 0.02 eV on A and 1.0 on B, 2,114 substitutions at 268 residues perturb connectivity without changing charge, and 9,621 substitutions at 2,021 residues change charge without touching the wire. After restricting to positions identical in mouse and human (below), these become 1,556 substitutions at 197 residues and 6,293 at 1,298 residues, with the correlations essentially unchanged (0.17 and 0.29). Residual coupling between the axes is geometric: loading sites and wire residues lie closer to each other than non-wire residues do (median distance to the nearest loading site 6.9 Å versus 11.8 Å). Distance to the nearest loading site is therefore carried as a covariate in the regressions below.

### Transfer from the mouse structure to human coordinates

The analysis requires human variant data but a structure with modelled hydration, and no single structure provides both. (3) Human complex I structures are available at comparable resolution — PDB 9TI4 (2.66 Å) and 9I4I (2.63 Å) — but they model one water molecule between them, because their depositors were studying complex IV maturation rather than proton pathways. (47) We therefore used the mouse structure for wire geometry and transferred the classification to human coordinates. (3) We verified that this is the only workable option by measuring the alternatives. Building the wire graph on the human 9TI4 side chains alone collapses it: 161 edges and 29 ND residues in a cluster, against 2,687 edges and 331 residues for mouse with its own waters. Superposing mouse onto human on 2,106 matched Cα atoms gives an RMSD of 1.57 Å — the fold is the same — but 2,052 of the 2,945 transferred waters then clash with a human heavy atom at under 2.2 Å (median nearest distance 1.83 Å), so a graph built on borrowed water positions is not physically admissible. (3, 47) The human structure is used instead as a fold control and to confirm that the membrane arm carries no human-specific insertion in the regions analysed. (47)

Human ND protein sequences were obtained by translating the revised Cambridge Reference Sequence (NC_012920.1, 16,569 bp) in the vertebrate mitochondrial code over the coding intervals 3307–4262 (ND1), 4470–5511 (ND2), 10059–10404 (ND3), 10470–10766 (ND4L), 10760–12137 (ND4), 12337–14148 (ND5) and 14149–14673 (ND6, light strand, reverse complemented). (48) All seven translations match their expected lengths exactly (318, 347, 115, 98, 459, 603 and 174 residues); ND1, ND2, ND3 and ND4 terminate in incomplete stop codons completed by polyadenylation. Mouse and human sequences were aligned per subunit with a global Needleman–Wunsch alignment (BLOSUM62, gap open −11, extend −1). (49, 50) Identity ranges from 52.9% (ND6) to 77.0% (ND1), with a mean of 65.0% over 2,106 aligned residues. (48) Wire residues are more conserved than the rest (75.7% versus 62.2% identical), but far from invariant. (40, 51) Primary analyses are restricted to the 1,367 positions at which the mouse and human residues are identical, where the structural inference transfers directly. (52) This retains 78 of 96 loading sites (81.2%) and 85 of 111 cut vertices (76.6%). The 93 wire residues that differ include chemically substantive changes — ND1 His93Asn, ND1 Lys126Asn, ND5 Glu75Gln, ND5 Lys77Ser, ND5 Asp352Asn, ND5 Lys547Asn — at which the human protein lacks the titratable group present in mouse; these positions are excluded rather than modelled. (20) Insertions and deletions are terminal in ND2 and ND5 but internal in ND6, which also has the lowest identity; ND6 results should be read with that in mind.

### Variant data

Germline and somatic mtDNA variant calls were taken from Gorelick et al. (github.com/reznik-lab/mtdna-mutations), which provides harmonised calls across cohorts in rCRS coordinates annotated in the vertebrate mitochondrial code. (11)

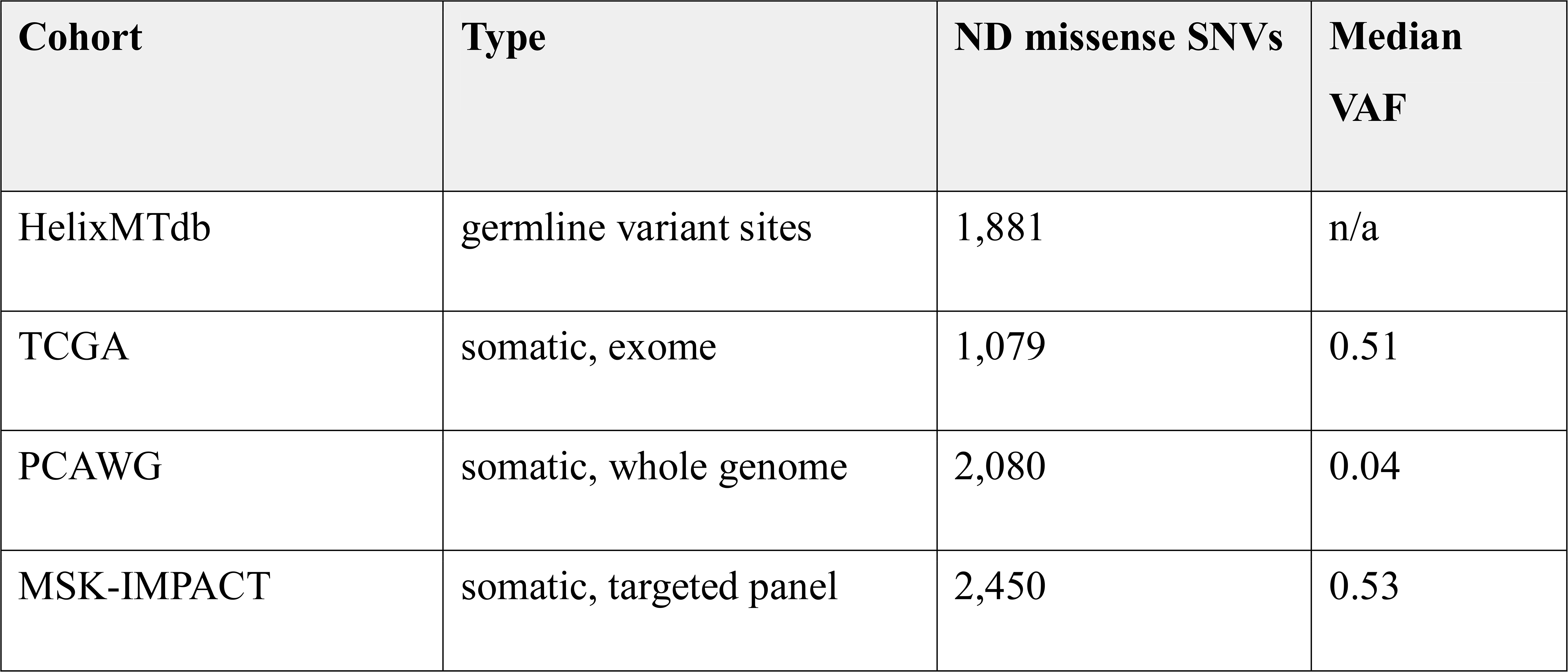

The three tumour cohorts differ markedly in heteroplasmy detection. PCAWG whole genomes reach mtDNA depths of roughly 9,000–11,000 reads and so call variants at much lower allele fractions than TCGA exomes; this is a difference in sensitivity, not in biology, and it motivates reporting all three cohorts separately rather than pooling them. Following the approach used for cytochrome b, somatic calls at sites carried by more than ten HelixMTdb individuals were removed as likely germline polymorphisms (566 of the possible substitutions), and expected counts for the somatic analyses were computed over the same restricted set of sites. (12) The germline analysis retains the full site universe, because the filter is defined by germline carrier counts and would otherwise remove the variants being analysed. Nuclear somatic mutations for the signature analysis were open-access ensemble-masked MAF files from the NCI Genomic Data Commons, covering 3,364,892 single-nucleotide variants in 9,393 patients across 30 TCGA cohorts, with the 11-bp reference context reported per call. Of these patients, 2,207 also have mtDNA coverage in the Gorelick calls and could be assigned a proton-wire status. (53, 54)

### Substitution universe and null model

The universe of possible substitutions was enumerated from first principles rather than taken from an annotation database. For every coding position of every ND gene and every alternate base, the mutant codon was translated in the vertebrate mitochondrial code and the substitution classified as synonymous, missense or nonsense, with ND6 handled on the light strand. This yields 19,026 substitutions over 2,114 codons: 13,447 missense, 4,668 synonymous and 911 nonsense. The missense set agrees with MitImpact v2.9 at 13,426 of 13,439 shared entries. Enumerating the universe directly matters for the calibration check. MitImpact lists only missense rows, so a synonymous set derived from it is biased toward first- and second-codon positions and gives a mis-calibrated null; built from first principles, the synonymous set is 94–100% third-codon positions in every class, as it should be. Expected counts come from each cohort’s own strand-aware trinucleotide spectrum. Each possible substitution is assigned one of 192 channels (the rCRS trinucleotide centred on the mutated base, plus the alternate allele), and channel probabilities are estimated from all mtDNA single-nucleotide variants in that cohort excluding coding variants in the ND genes, so the null is independent of the data being tested. Probabilities were Laplace-smoothed by 0.5. The spectra rest on 7,310 (HelixMTdb), 2,359 (TCGA), 5,170 (PCAWG) and 6,179 (MSK-IMPACT) variants. (54, 55)

The expected count for a residue class is the total observed count in the universe multiplied by that class’s share of summed channel probabilities. Enrichment was tested with two-sided exact Poisson tests, and confidence intervals are exact Poisson intervals.

#### Calibration

The null model was validated on synonymous ND variants, which should be neutral. Observed/expected ratios are 0.87–1.01 in the germline (1,622 variants) and span 0.33–1.34 in the tumour cohorts, where synonymous counts are small (expected 0.5–4.6). No class deviates from expectation in any cohort (minimum P = 0.17).

### Selection statistics

Rate ratios were estimated by quasi-Poisson regression at the level of the individual possible substitution, with the logarithm of its channel probability as an offset and the proton-wire class as a categorical predictor against the reference class of other ND residues. Dispersion was estimated by Pearson chi-squared. Models were fitted separately for each cohort on 6,825–6,999 substitutions.

Covariates were added in a fixed ladder, and the loading-site coefficient reported at each step:

1. unadjusted;
2. plus PhastCons (100 vertebrates), as a measure of cross-species conservation;
3. plus burial (heavy atoms within 8 Å of the side-chain centroid), side-chain volume change and Kyte–Doolittle hydropathy change;
4. plus distance to the nearest proton loading site.

PhastCons was taken from MitImpact v2.9 (82.5% coverage of possible missense substitutions). The PhyloP column of that release is unusable: it is a −20 sentinel for all but 0.8% of entries, and treating it as a score silently reduces the analysis to a few hundred substitutions. Conservation adjustment therefore rests on PhastCons alone, which is a limitation. We deliberately did not adjust the germline analysis for mitochondrial local constraint (MLC). MLC is derived from depletion of variation in human population data, which is the quantity the germline analysis measures; adjusting one by the other would remove the signal by construction rather than because it is spurious. Cross-species conservation is the appropriate covariate for the germline arm. MLC remains a legitimate covariate for the tumour arm, where it is derived from independent data.

Heteroplasmy was analysed by stratifying somatic missense variants at a variant allele fraction of 0.5, as in the cytochrome b analysis. For TCGA, allele fractions were additionally corrected for tumour purity as VAF divided by ABSOLUTE purity, capped at 1, using the published pan-cancer purity table. Loading-site enrichment was compared between strata, and the heteroplasmy distribution of loading-site missense variants was compared with that of truncating ND variants by two-sided Mann–Whitney tests. Per-subunit enrichment was computed with the same machinery restricted to each ND gene, and reported only where the expected count is at least three.

### Sensitivity analyses

Four sensitivity analyses were run. The hydrogen-bond cutoff was varied over 3.2, 3.5 and 4.0 Å, rebuilding the graph, the clusters, the articulation points and the residue classes at each value and repeating the full observed/expected analysis. The restriction to positions identical in mouse and human was removed, extending the analysis to all 2,110 transferable positions. The germline-polymorphism filter was removed from the somatic cohorts. The steric term of axis B was removed, leaving the topological term alone.

### Disease variants and nuclear mutational signatures

#### Disease variants

Pathogenic annotations were taken from MitImpact v2.9, which carries MITOMAP status and phenotype and ClinVar significance. Two sets were defined: variants with MITOMAP status Cfrm (21 within the analysed set) and a broader set combining Cfrm, Reported, point-mutation flags and ClinVar Pathogenic (115). Enrichment by wire class was computed against the germline trinucleotide spectrum as above. The confirmed set is underpowered — expected counts per class are 1.0 to 2.0 — and we report it with exact Poisson intervals rather than drawing conclusions from it. These annotations date from the 2017 ClinVar release carried by MitImpact v2.9. Overlap between disease and tumour variant sets was tested by Fisher’s exact test on the 2×2 table of reported-pathogenic status against observation in any tumour cohort.

#### Nuclear signatures

For each patient, a 96-channel single-base substitution profile was built from the MAF reference context, folding to a pyrimidine reference. Profiles with at least 20 substitutions (8,058 tumours) were fitted to COSMIC v3.4 GRCh38 exome signatures, excluding the possible-artefact subset, by non-negative least squares with backward elimination of components contributing under 3% of the burden; the median reconstruction cosine similarity was 0.91. Four oxidative readouts were tested: the C>A fraction of nuclear substitutions, the fraction of G>T substitutions falling at a guanine within a run of two or more guanines (read on the purine strand from the 11-bp context, a sequence-level proxy for hole-trapping damage), and the presence of SBS18 and SBS36. Patients were grouped by the highest-ranking proton-wire class among their ND missense variants, against a reference of tumours with mtDNA coverage and no ND missense variant. Continuous readouts were centred within cancer type and compared by permutation restricted to cancer types containing both groups (10,000 permutations). Signature exposures are zero-inflated, so SBS18 and SBS36 were tested as presence or absence with a Cochran–Mantel–Haenszel test stratified by cancer type. The sixteen tests were corrected together by the Benjamini–Hochberg procedure.

### Structural concordance, recurrence, phylogeny and the interaction test

#### Structural concordance

For each residue mapped between the two structures, the Cα atoms of all mapped residues within 10 Å were superposed onto their human counterparts by the Kabsch algorithm, requiring at least five residues in the patch. Two quantities were recorded: the root-mean-square deviation of the patch after superposition, and the distance between the mouse and human side-chain centroids under the same transformation. A residue was called concordant at a local RMSD of at most 1.0 Å and a side-chain deviation of at most 2.0 Å. The selection analysis was repeated on the codons that are concordant and identical in sequence, with the null model and all filters unchanged.

#### Recurrence

Among the substitutions eligible for the somatic analysis, the number of patient-level calls was held fixed and redistributed over sites by a multinomial draw with the cohort’s own trinucleotide weights, 10,000 times. The statistic is the number of sites in a wire class carrying calls from two or more patients, and the P value is the fraction of draws reaching the observed count. Duplicate calls of the same substitution in the same patient were collapsed.

#### Clonality

Allele fractions of loading-site, cut-vertex and remaining missense variants were compared by the Mann-Whitney test against the missense variants outside the wire, and the proportion at or above an allele fraction of 0.9 by Fisher’s exact test. Truncating ND variants (nonsense and frameshift) were included as the comparison class most plausibly under selection.

#### Phylogeny

Branch-defining substitutions were taken from PhyloTree build 17 as distributed with HaploGrep. Reversions, insertions, deletions, heteroplasmic calls and positions flagged as uncertain were discarded, and the remaining tokens resolved against the rCRS to give reference and alternate alleles. Enrichment was tested against the germline spectrum by the same observed/expected framework as the cohorts.

#### Interaction test

Counts for every eligible substitution in all four cohorts were stacked into one table and fitted by quasi-Poisson regression on wire class, cohort and the product of wire class with a tumour indicator, with the logarithm of the spectrum-weighted expected count as offset. The tumour indicator was not entered as a main effect because it is a linear combination of the cohort terms. Standard errors were clustered on the substitution, since the same substitution contributes a row to each cohort. The model was fitted with and without the conservation and structural covariates, and with and without the germline-polymorphism filter applied to both arms.

### Coverage, stratification and the jackknife

#### Coverage

The per-sample, per-base coverage matrices distributed with the TCGA and MSK-IMPACT calls were reduced to the fraction of samples covering each base, counting only samples with an entry at that base. Coverage was compared between wire classes by the Mann-Whitney test and entered into the selection model on the log scale. PCAWG has no coverage matrix and is whole-genome.

#### Cancer types

Each cancer type with at least twenty eligible calls was scored against the share of the cohort’s spectrum-weighted opportunity falling on the class, with exact Poisson intervals. Heterogeneity was tested by Cochran’s Q on the log ratios with Poisson weights, and the pooled ratio recomputed with each type removed.

#### Jackknife

The selection analysis was repeated seven times with one ND gene removed, and once for each gene alone, with the spectrum and all filters unchanged.

#### Axis A parameters

Axis A was recomputed over effective dielectric constants of 4, 8 and 20 and distance floors of 2, 3 and 5 Å. Because the dielectric scales all scores by one factor, the variant-level contrasts were repeated with a quantile threshold in place of the fixed 0.02 eV cut.

#### Multiple testing

Primary tests were grouped into seven families and corrected by Benjamini-Hochberg within each family and across the whole set; both q values are reported.

#### Structural context

Residue positions were expressed in the membrane frame as the projection of the side-chain centroid onto the membrane normal and onto the principal in-plane axis. The quinone cavity was located as the centroid of the ligand WYK bound at the ND1 interface in 8OM1, and distances measured to it. Loading sites were split into those of ND1 within 15 Å of that centroid, those of ND1 further away, and those of the other subunits, and each group scored by the same observed/expected framework. The central axis was taken as the charged loading sites lying within 6 Å of the bilayer midplane in ND2, ND4 and ND5.

#### Somatic support

Matched-normal depth and alternate-allele counts were taken from the released calls. A normal was counted as informative at twenty reads or more. Allele fractions at or above 0.9 were called clonal, and compared between substitutions by Fisher’s exact test. Homopolymer context was taken from the annotation accompanying the calls.

#### Robustness of the localisation

The radius defining proximity to the quinone cavity was swept from 10 to 25 Å. Distance was then entered directly as minus the logarithm of the distance in ångström, so that the coefficient is a rate ratio per e-fold decrease, in models with and without subunit fixed effects and the covariates, and in a pooled model over all four cohorts with the subunit effects free to differ between the germline and tumour arms and standard errors clustered on the substitution. The mean distance of observed calls was compared against 10,000 multinomial redistributions under the cohort spectrum. For the cut-vertex null, the smallest true ratio detectable with 80% power at the observed expected count was computed from the Poisson distribution.

## Data and code availability

Structures are available from the Protein Data Bank under accessions 8OM1 (mouse complex I, active state, 2.39 Å), 9TI4 and 9I4I (human complex I). The revised Cambridge Reference Sequence is NCBI NC_012920.1.

Germline and somatic mtDNA variant calls for HelixMTdb, TCGA, PCAWG and MSK-IMPACT, together with the tumour purity tables, were obtained from Gorelick et al. (https://github.com/reznik-lab/mtdna-mutations). Conservation scores and pathogenicity annotations are from MitImpact v2.9, distributed with the same repository. Nuclear somatic variants are open-access ensemble-masked MAF files from the NCI Genomic Data Commons (https://portal.gdc.cancer.gov). COSMIC v3.4 reference signatures were taken from the SigProfilerAssignment distribution.

Custom code for the proton-wire graph, cluster and cut-vertex detection, the two axes, mouse-to-human mapping, universe enumeration, null-model construction, selection statistics and signature fitting will be provided as Supplementary Software.

## Acknowledgements

We thank the TCGA Research Network, the ICGC/TCGA Pan-Cancer Analysis of Whole Genomes (PCAWG) Consortium and the Helix team for making their data publicly available, and the authors of Gorelick et al. for releasing their mtDNA mutation calls. This work was supported by a grant from the Korean ARPA-H Project through the Korea Health Industry Development Institute (KHIDI), funded by the Ministry of Health & Welfare, Republic of Korea (grant number RS-2025-25456722), and by the ANCHOR through the Seoul ANCHOR Center, funded by the Ministry of Education (MOE) and the Seoul Metropolitan Government (2026-ANCHOR-01-022-05), to J.-H.C.

## Author Contributions

**J.-Y.S.** conceived and designed the research, developed the computational framework, performed structural modeling and bioinformatic analyses, collected and curated data, conducted statistical analyses, interpreted the results, prepared the figures, and wrote, reviewed, and revised the manuscript. **J.-H.C**. acquired funding, contributed to the interpretation of the results, and reviewed and revised the manuscript. Both authors read and approved the final manuscript.

## Supplementary Figures

**Supplementary Fig. 1.**
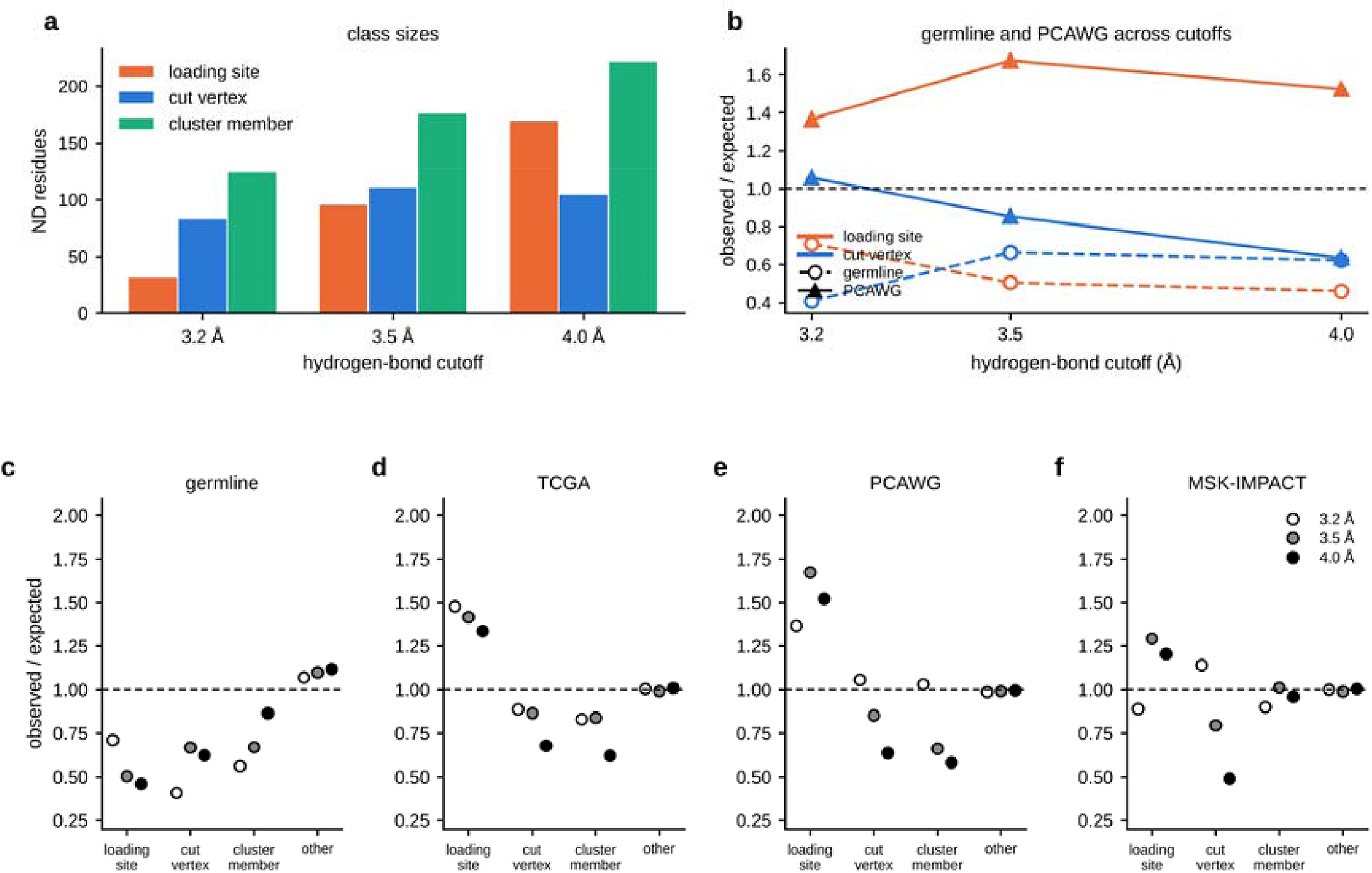
| Hydrogen-bond cutoff sensitivity. a Residue class sizes at donor–acceptor cutoffs of 3.2, 3.5 and 4.0 Å. b Loading-site and cut-vertex observed/expected ratios in the germline (open symbols, dashed) and PCAWG (filled, solid) across the three cutoffs. c–f Observed/expected by proton-wire class at each cutoff, one panel per cohort; open, grey and filled circles are 3.2, 3.5 and 4.0 Å. The loading-site pattern holds at 3.5 and 4.0 Å but not at 3.2 Å, where only 32 loading sites survive and the comparison loses power. Cut vertices are never enriched at any cutoff and become depleted in tumours at 4.0 Å.

**Supplementary Fig. 2.**
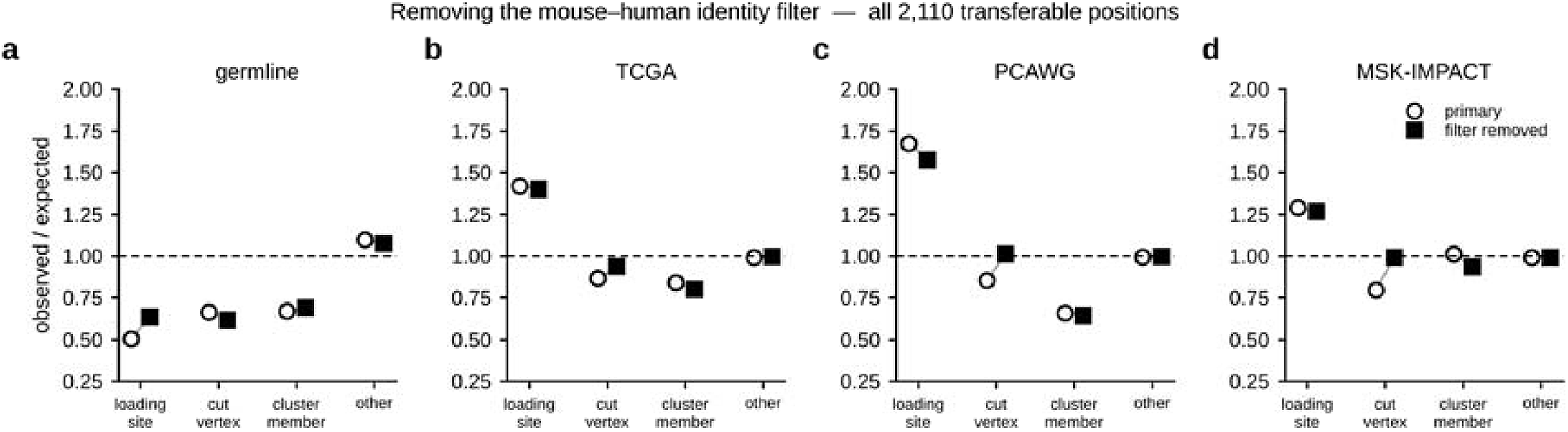
| Removing the mouse–human identity filter. a–d Observed/expected by proton-wire class under the primary restriction to positions identical in mouse and human (open circles) and across all 2,110 transferable positions (filled squares), one panel per cohort. The loading-site result keeps its direction and significance in every cohort.

**Supplementary Fig. 3.**
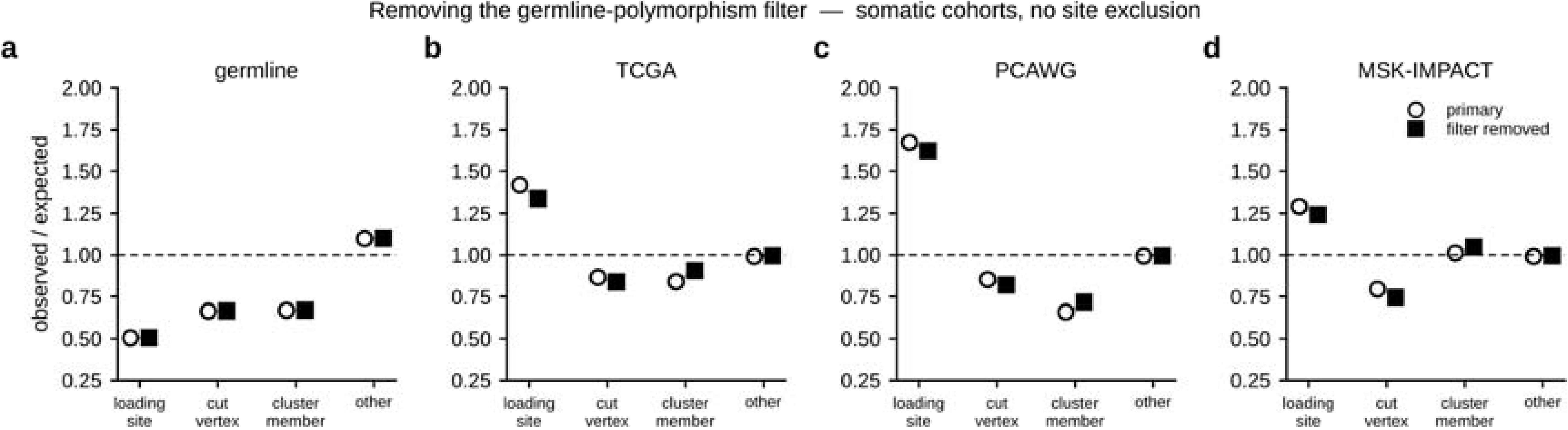
| Removing the germline-polymorphism filter. a–d As Supplementary Fig. 2, with somatic calls at sites carried by more than ten HelixMTdb individuals retained rather than removed. Somatic estimates are essentially unchanged.

**Supplementary Fig. 4.**
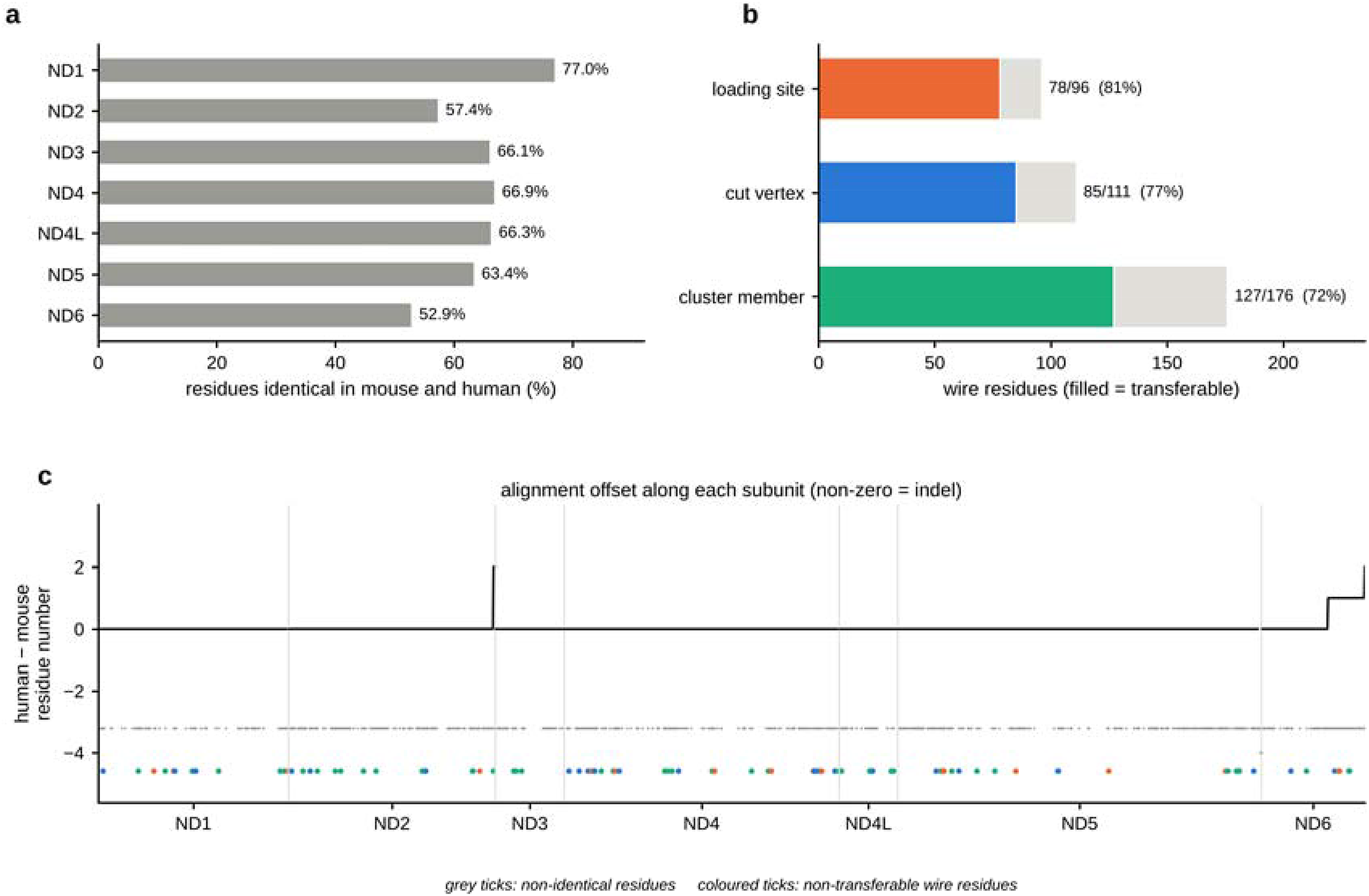
| Mouse-to-human transfer of the proton-wire classification. a Percentage of residues identical in mouse and human, per subunit. b Wire residues retained by the identity filter, by class; the filled part of each bar is transferable. c Alignment offset along each subunit, where a non-zero value marks an insertion or deletion; offsets are terminal in ND2 and ND5 but internal in ND6. Grey ticks mark non-identical residues and coloured ticks the wire residues that cannot be transferred.

**Supplementary Fig. 5.**
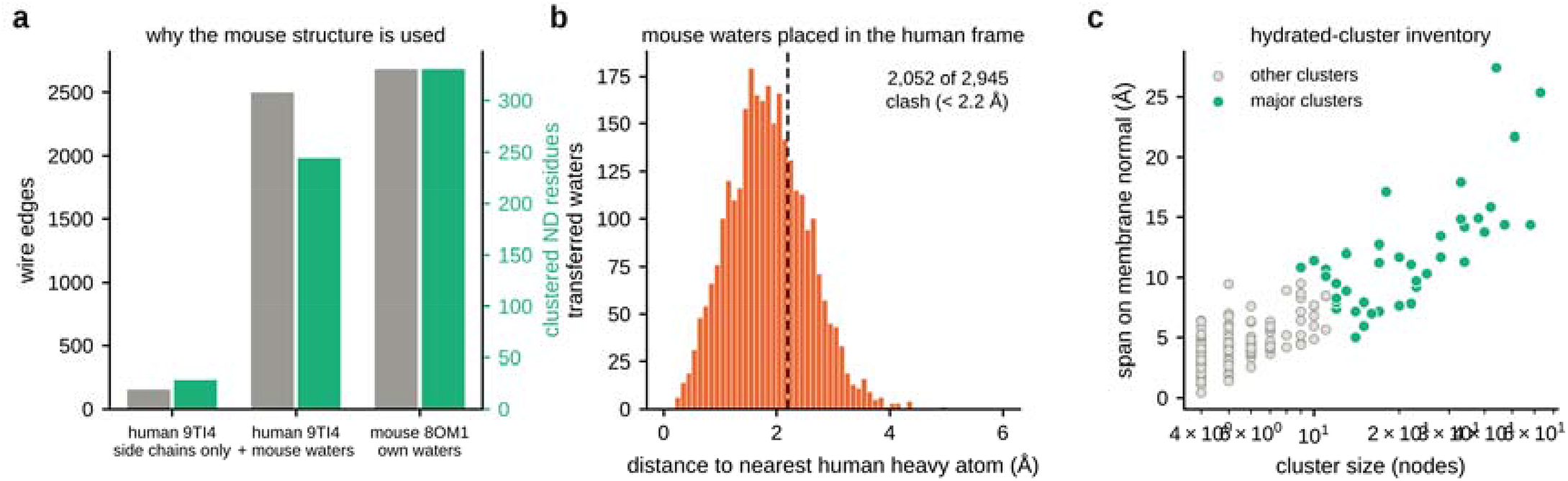
| Why the wire geometry is taken from the mouse structure. a Wire edges (grey, left axis) and clustered ND residues (green, right axis) under three strategies: human 9TI4 side chains alone, human side chains with mouse waters superposed, and mouse 8OM1 with its own waters. b Distance from each mouse water, placed in the human frame by superposition, to the nearest human heavy atom; the dashed line marks 2.2 Å. c Hydrated clusters by size and span along the membrane normal, with the major clusters used to define proton loading sites highlighted.

**Supplementary Fig. 6.**
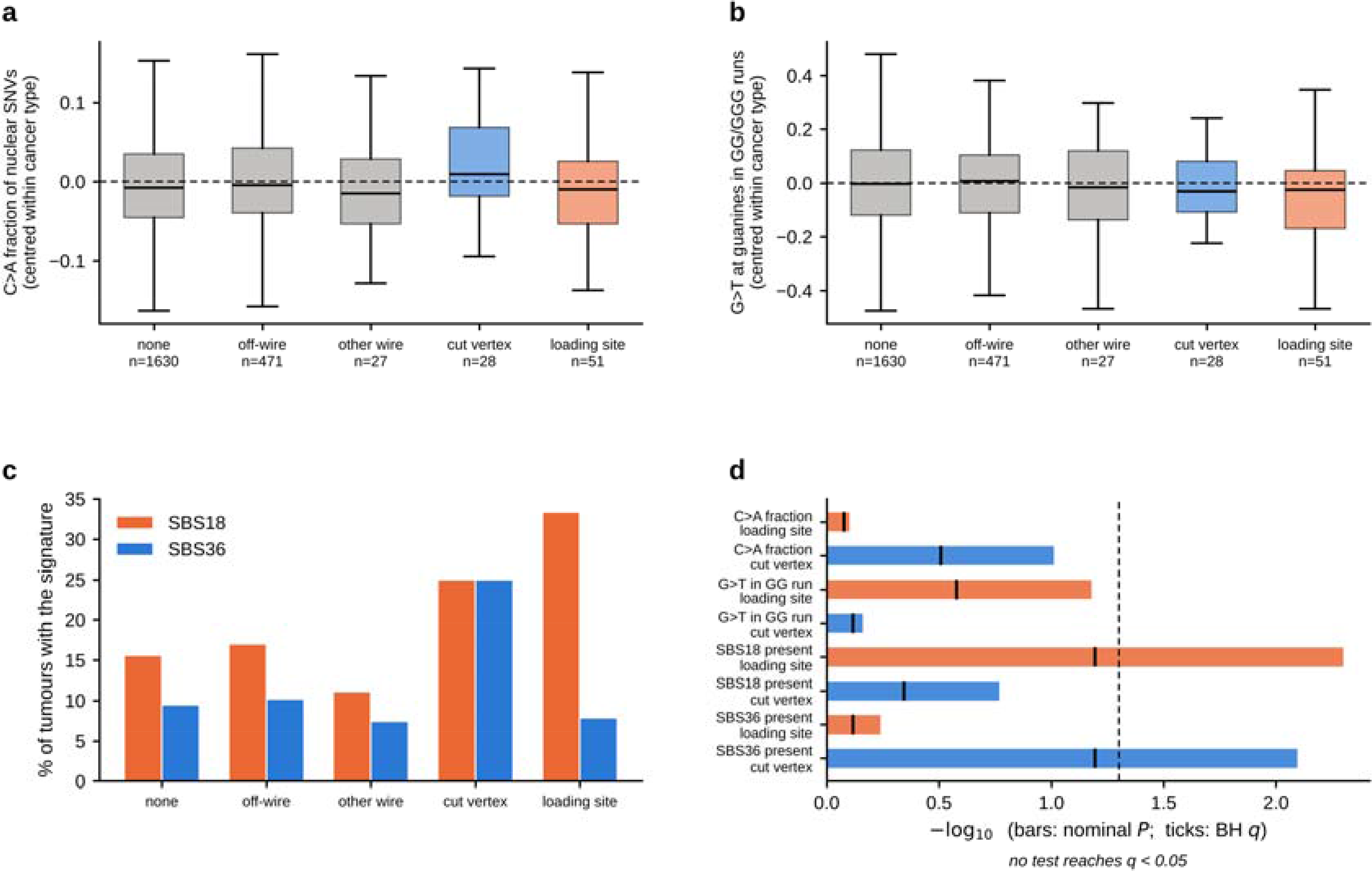
| Nuclear mutational readouts by proton-wire status. a, b C>A fraction of nuclear substitutions and fraction of G>T substitutions at guanines within GG/GGG runs, centred within cancer type, by group. Boxes give the interquartile range and the median; whiskers extend to 1.5 times the interquartile range. c Percentage of tumours carrying SBS18 or SBS36. d All sixteen tests: bars give −log10 of the nominal two-sided P value and vertical ticks the Benjamini–Hochberg q value; the dashed line marks P = 0.05. No test reaches q < 0.05.

**Supplementary Fig. 7.**
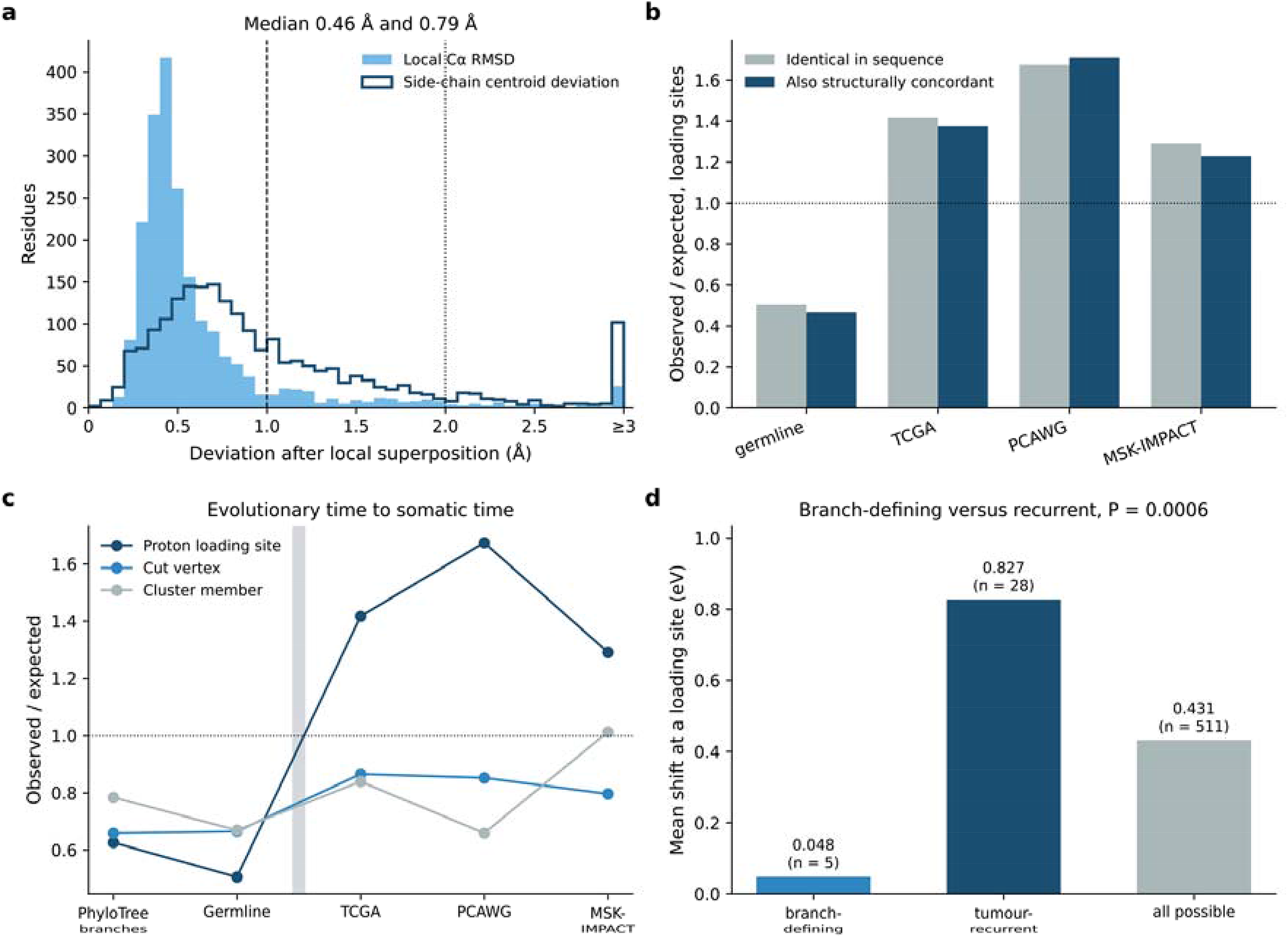
| Structural concordance, evolutionary timescale and the size of the electrostatic shift. a Local agreement between the mouse and human structures: the root-mean-square deviation of the Cα atoms within 10 Å of each mapped residue after local superposition (filled), and the distance the side-chain centroid moves under the same transformation (outline). Dashed and dotted lines mark the 1.0 Å and 2.0 Å thresholds; values above 3 Å are collected in the last bin. **b** Observed/expected for loading-site missense substitutions restricted to codons identical in sequence (grey) and to those that are also structurally concordant (blue). **c** Observed/expected by wire class across three timescales: branch-defining substitutions of PhyloTree build 17, the present-day germline, and the three tumour cohorts. The band separates the evolutionary from the somatic comparisons. **d** Mean electrostatic shift imposed at a proton loading site by the substitutions that fixed on a phylogenetic branch, by those recurring in four or more tumours, and by all possible loading-site substitutions.

**Supplementary Fig. 8.**
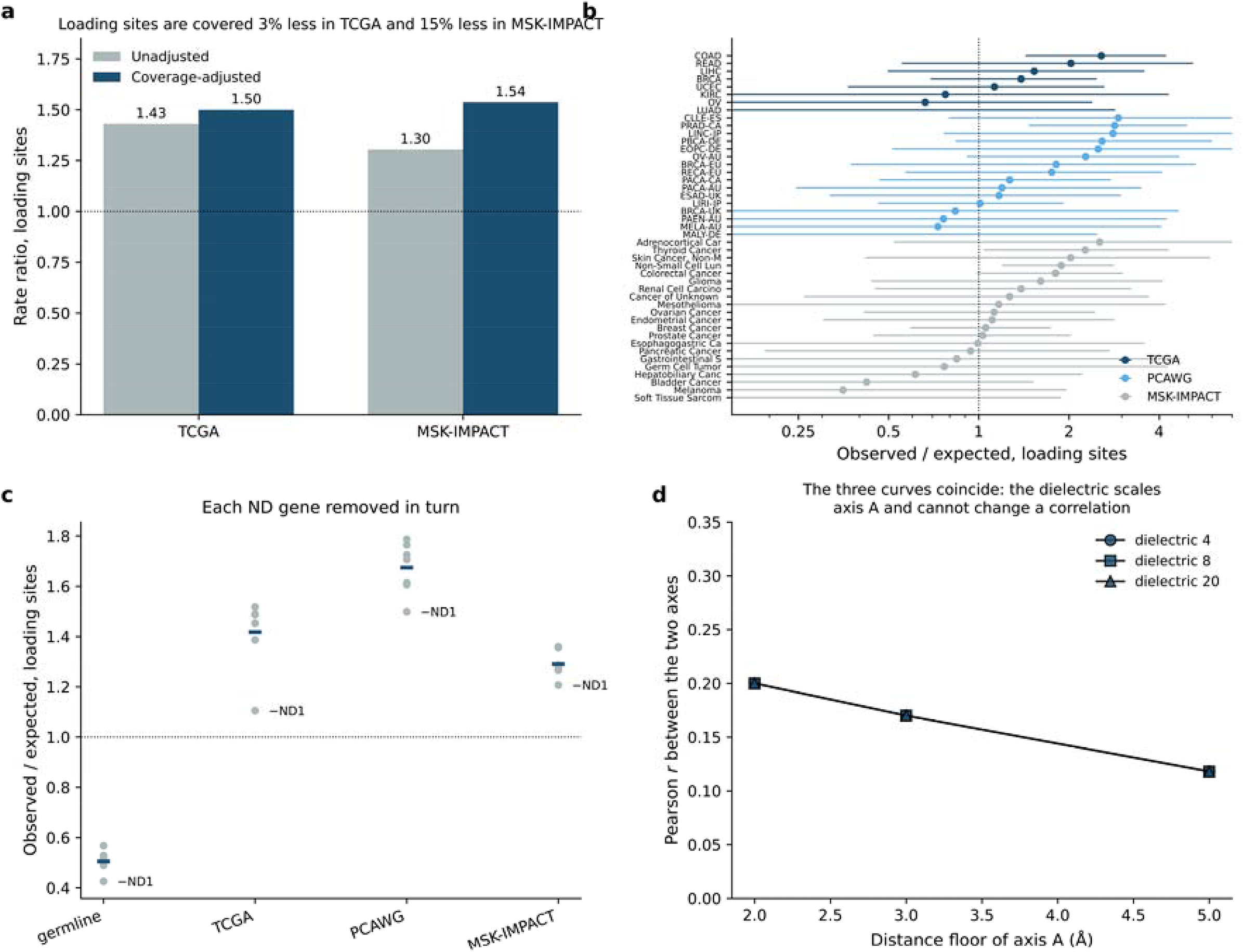
| Technical and compositional controls. a Loading-site rate ratio before and after adjusting for per-base sequencing coverage, in the two cohorts for which a coverage matrix exists. Loading sites are covered less than other ND positions, so the adjustment raises the estimate. **b** Observed/expected for loading-site missense substitutions by cancer type, for every type with at least twenty eligible calls, with exact Poisson intervals. **c** Observed/expected with each ND gene removed in turn (grey) against the estimate on all seven genes (bar). The replicate labelled in each cohort is the lowest. **d** Correlation between the two axes over the parameters of axis A. The three dielectric curves coincide because the dielectric scales axis A and cannot change a correlation.

**Supplementary Fig. 9.**
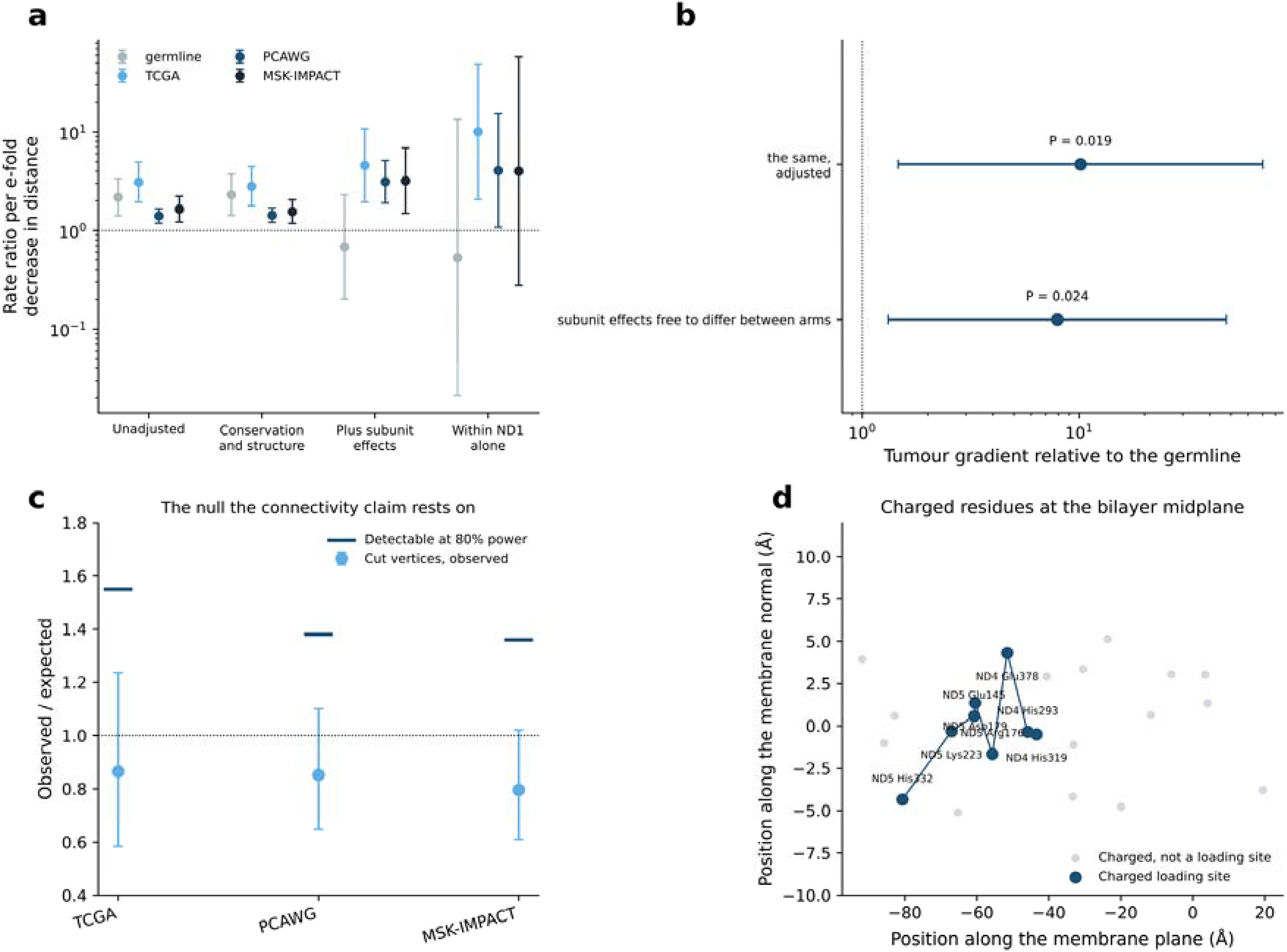
| Robustness of the quinone localisation, and the charge axis of the antiporter-like subunits. a Rate ratio per e-fold decrease in distance to the cavity, through a ladder of adjustments. b The tumour gradient relative to the germline, from a pooled model with subunit effects free to differ between arms. c Cut-vertex observed/expected with exact Poisson intervals, against the smallest true ratio each cohort could have detected with 80% power. d Charged residues within 6 Å of the bilayer midplane in the antiporter-like subunits; those classified as loading sites are joined in order along the membrane plane.

**Supplementary Fig. 10.**
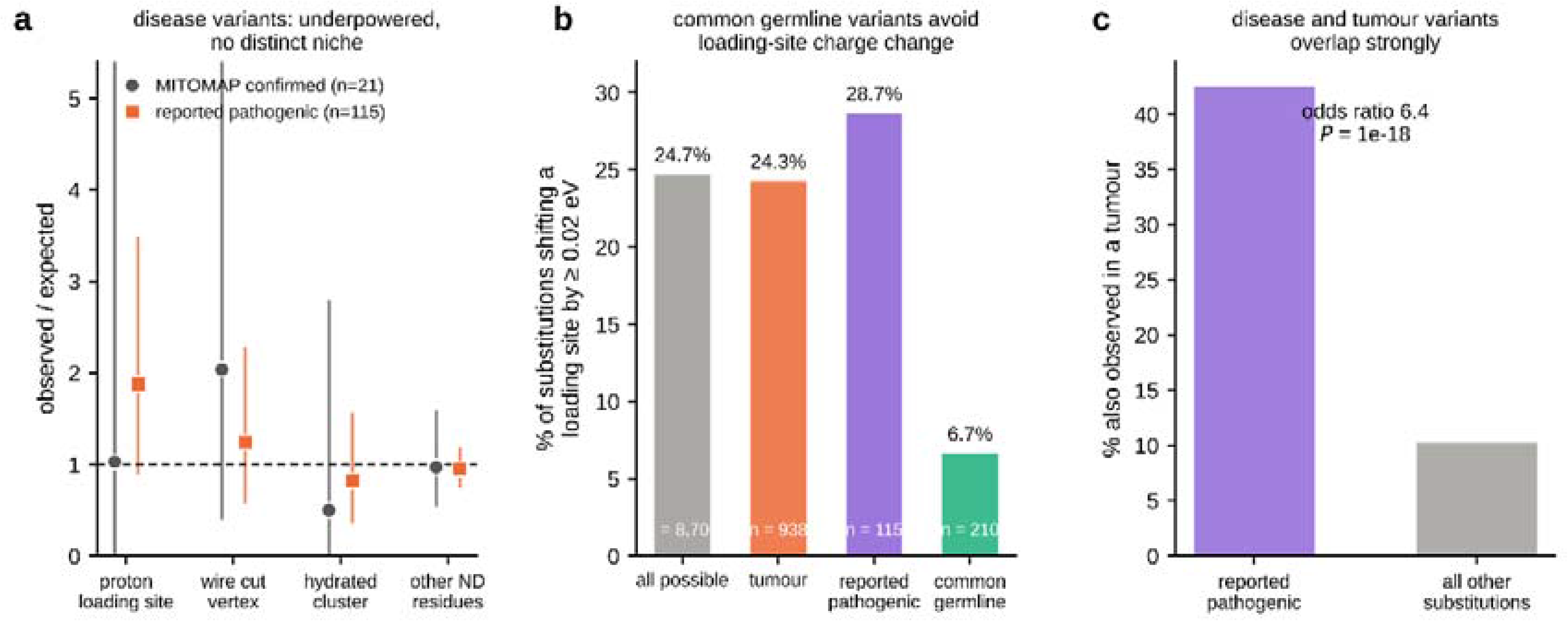
| Disease variants do not occupy a distinct niche, and common germline variants avoid loading-site charge change. a Observed/expected ratios by proton-wire class for MITOMAP-confirmed pathogenic ND missense variants (n = 21) and for a broader reported-pathogenic set (n = 115), against the germline trinucleotide spectrum. Neither set is significantly enriched in any class; expected counts for the confirmed set are 1.0–2.0 per class, so this comparison is underpowered rather than negative. Error bars, 95% exact Poisson confidence intervals (upper bounds clipped at 5.4); two-sided exact Poisson tests. b Percentage of substitutions shifting a proton loading site by at least 0.02 eV, by variant group. Common germline variants (carried by more than ten HelixMTdb individuals) are strongly depleted of such substitutions relative to all possible substitutions (6.7% versus 24.7%), whereas tumour-observed and reported-pathogenic variants are not. c Percentage of substitutions also observed in at least one tumour cohort, for reported-pathogenic variants versus all other substitutions. Disease and tumour variant sets overlap strongly (two-sided Fisher’s exact test, odds ratio 6.4, P = 9.7 × 10□¹□). Source data are provided as a Source Data file.

**Supplementary Table 1.**
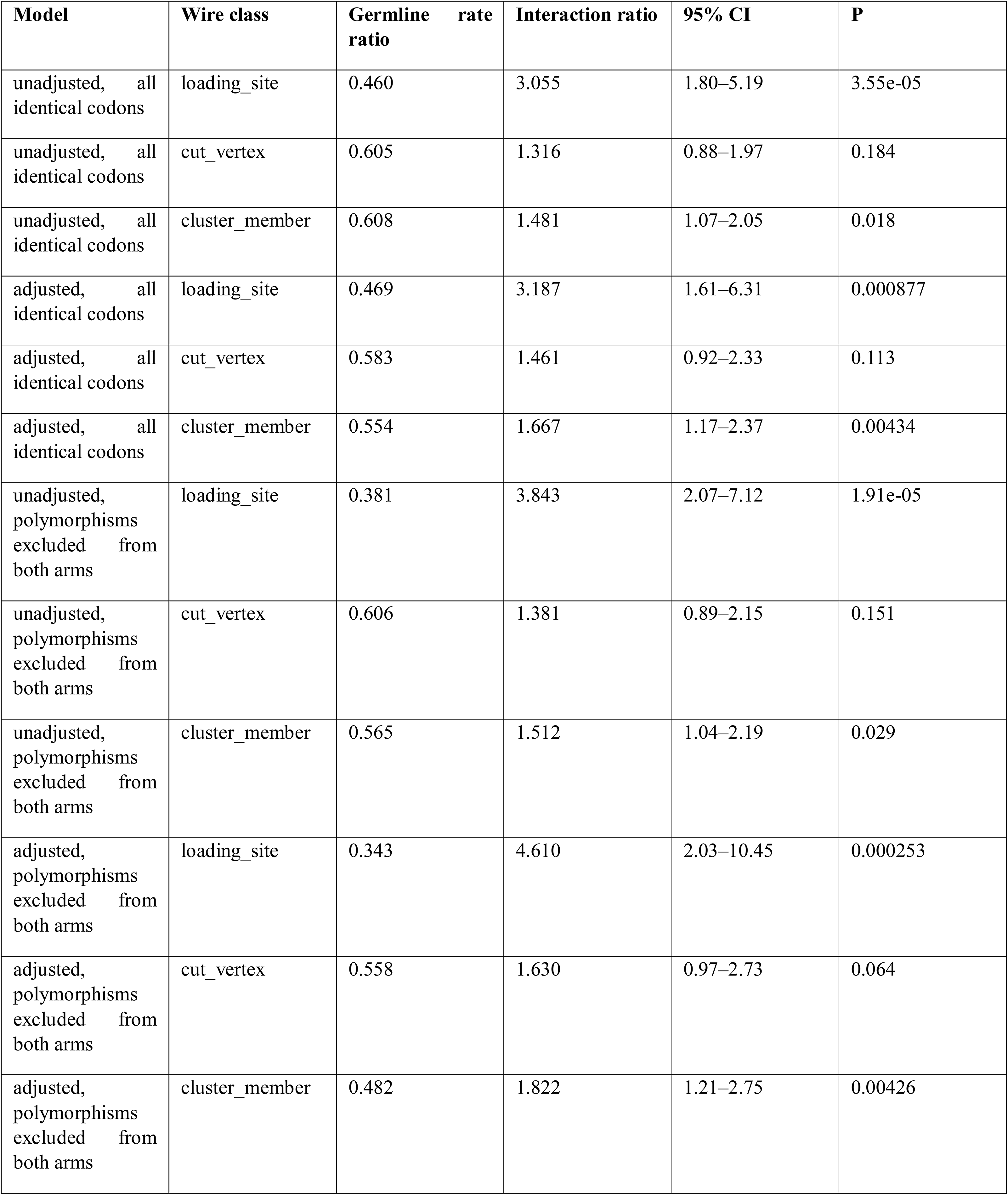
| Wire class by tissue-type interaction. The germline rate ratio compares the class with ND residues outside the wire. The interaction ratio is the factor by which that ratio changes in tumours, and is the test of relaxed constraint. Quasi-Poisson regression, standard errors clustered on the substitution.

**Supplementary Table 2.**
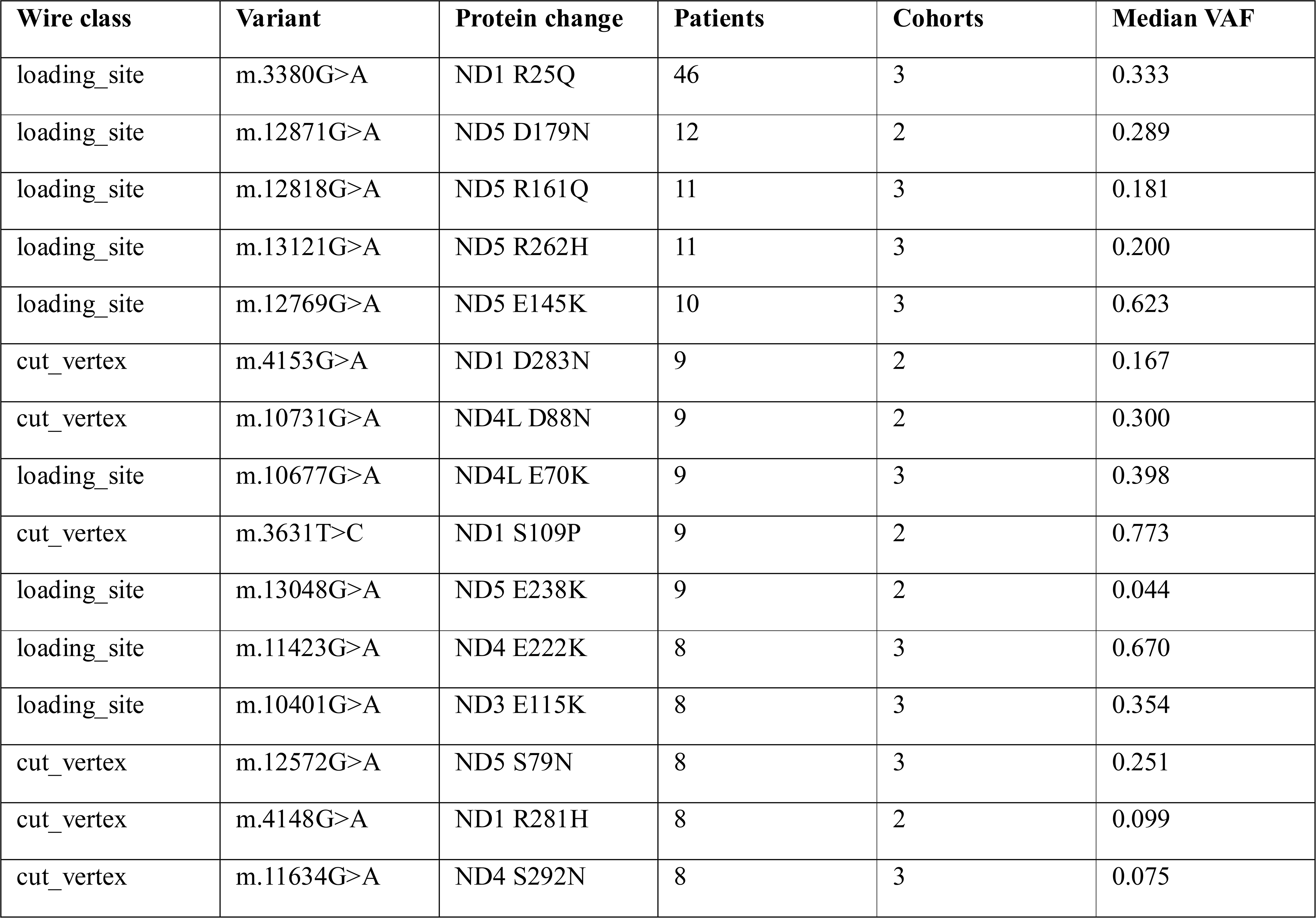
| Recurrent proton-wire substitutions in tumours. Substitutions of the wire classes seen in at least four patients, pooled over TCGA, PCAWG and MSK-IMPACT. Cohorts is the number of the three in which the substitution appears. The full list is given in the Source Data.

**Supplementary Table 3.**
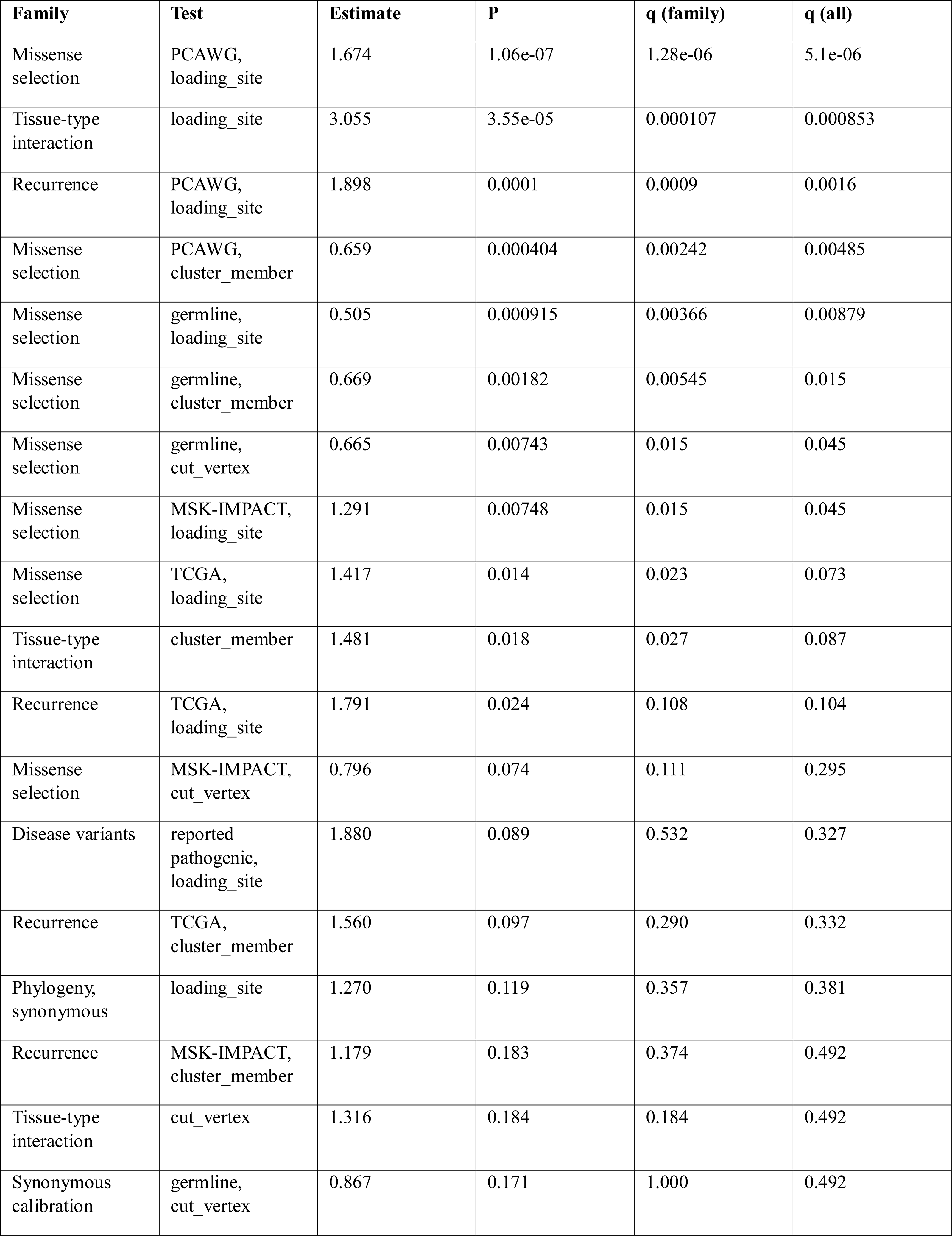
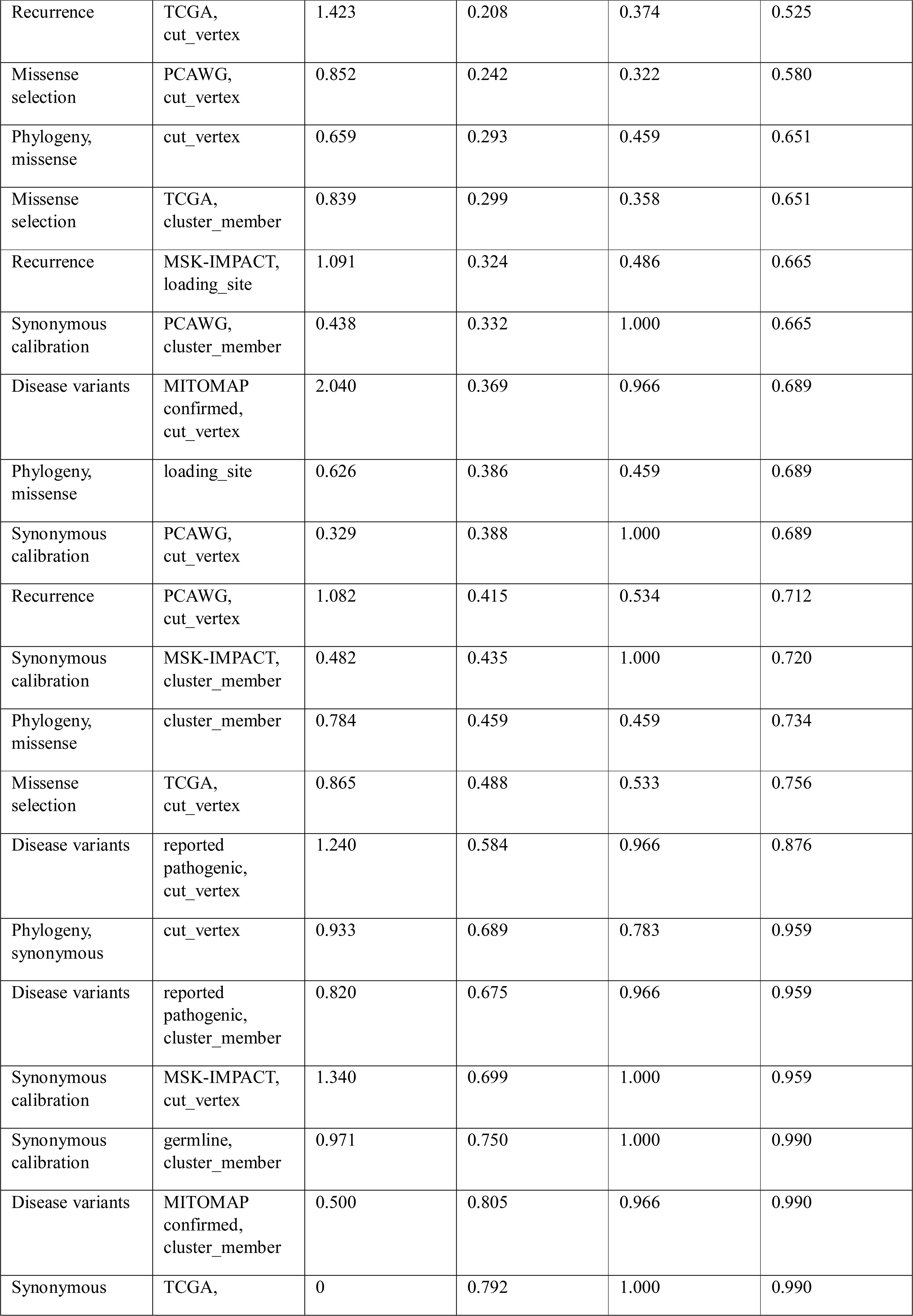
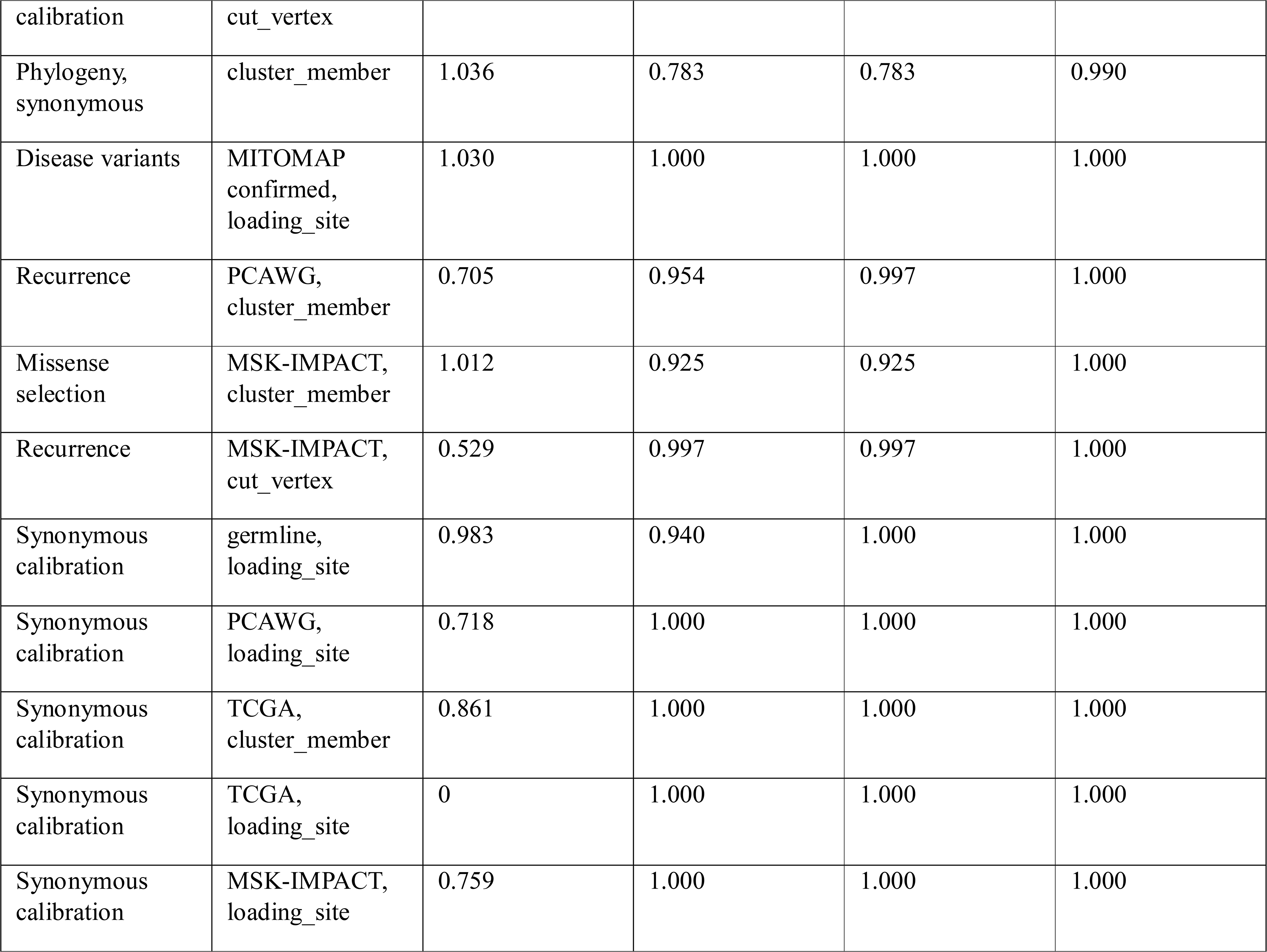
| Multiple testing across the paper. Primary tests grouped into families, corrected by Benjamini-Hochberg within each family and over the whole set. Estimates are the ratio or rate ratio reported in the text.

## Reference

1. Beghiah A, Saura P, Kovalova T, Hoeser F, Friedrich T, Kaila VRI. A carboxylate switch point controls long-range energy transduction in respiratory Complex I. Nature Communications. 2026;17(1).

2. Choi JH, Voth GA. Hydration-Controlled Proton Transport in Respiratory Complex I. Journal of the American Chemical Society. 2026;148(13):13954–63.

3. Grba DN, Chung I, Bridges HR, Agip A-NA, Hirst J. Investigation of hydrated channels and proton pathways in a high-resolution cryo-EM structure of mammalian complex I. Science Advances. 2023;9(31).

4. Chung I, Wright JJ, Bridges HR, Ivanov BS, Biner O, Pereira CS, et al. Cryo-EM structures define ubiquinone-10 binding to mitochondrial complex I and conformational transitions accompanying Q-site occupancy. Nature Communications. 2022;13(1).

5. Grba DN, Hirst J. Mitochondrial complex I structure reveals ordered water molecules for catalysis and proton translocation. Nature Structural & Molecular Biology. 2020;27(10):892–900.

6. Di Luca A, Gamiz-Hernandez AP, Kaila VRI. Symmetry-related proton transfer pathways in respiratory complex I. Proceedings of the National Academy of Sciences. 2017;114(31).

7. Uddin MR, Khaniya U, Gupta C, Mao J, Ranepura GA, Wei RJ, et al. Finding the E-channel proton loading sites by calculating the ensemble of protonation microstates. Biochimica et Biophysica Acta (BBA) - Bioenergetics. 2025;1866(1).

8. Kratochvil HT, Watkins LC, Mravic M, Thomaston JL, Nicoludis JM, Somberg NH, et al. Transient water wires mediate selective proton transport in designed channel proteins. Nature Chemistry. 2023;15(7):1012–21.

9. Pomès R, Roux B. Molecular Mechanism of H+ Conduction in the Single-File Water Chain of the Gramicidin Channel. Biophysical Journal. 2002;82(5):2304–16.

10. Lake NJ, Ma K, Liu W, Battle SL, Laricchia KM, Tiao G, et al. Quantifying constraint in the human mitochondrial genome. Nature. 2024;635(8038):390–7.

11. Gorelick AN, Kim M, Chatila WK, La K, Hakimi AA, Berger MF, et al. Respiratory complex and tissue lineage drive recurrent mutations in tumour mtDNA. Nature Metabolism. 2021;3(4):558–70.

12. Yuan Y, Ju YS, Kim Y, Li J, Wang Y, Yoon CJ, et al. Comprehensive molecular characterization of mitochondrial genomes in human cancers. Nature Genetics. 2020;52(3):342–52.

13. Smith ALM, Whitehall JC, Greaves LC. Mitochondrial DNA mutations in ageing and cancer. Molecular Oncology. 2022;16(18):3276–94.

14. Sung J-Y & C, J.-H. Relaxed evolutionary constraints on mitochondrial electron-transfer-associated variants in human cancers. bioRXiv preprint at doi: https://doiorg/10.64898/20261021760641. 2026.

15. Sung J-Y & C, J.-H. Quantum transport in mitochondrial complex I is governed by a conserved structural bottleneck. bioRxiv https://doiorg/10.64898/20260528728423. 2026.

16. Sazanov LA. A giant molecular proton pump: structure and mechanism of respiratory complex I. Nature Reviews Molecular Cell Biology. 2015;16(6):375–88.

17. Vinothkumar KR, Zhu J, Hirst J. Architecture of mammalian respiratory complex I. Nature. 2014;515(7525):80–4.

18. Kaila VRI, Wikström M, Hummer G. Electrostatics, hydration, and proton transfer dynamics in the membrane domain of respiratory complex I. Proceedings of the National Academy of Sciences. 2014;111(19):6988–93.

19. Khaniya U, Gupta C, Cai X, Mao J, Kaur D, Zhang Y, et al. Hydrogen bond network analysis reveals the pathway for the proton transfer in the E-channel of T. thermophilus Complex I. Biochimica et Biophysica Acta (BBA) - Bioenergetics. 2020;1861(10).

20. Badolato S, Rossmann SC, Pereira J, Kim H, Kaila VRI. Molecular Principles of Gating Proton Transport in the Antiporter Modules of Respiratory Complex I. Journal of the American Chemical Society. 2026;148(20):21056–68.

21. Jarman OD, Hirst J, Dupont C. Membrane-domain mutations in respiratory complex I impede catalysis but do not uncouple proton pumping from ubiquinone reduction. PNAS Nexus. 2022;1(5).

22. Pätsi J, Maliniemi P, Pakanen S, Hinttala R, Uusimaa J, Majamaa K, et al. LHON/MELAS overlap mutation in ND1 subunit of mitochondrial complex I affects ubiquinone binding as revealed by modeling in Escherichia coli NDH-1. Biochimica et Biophysica Acta (BBA) - Bioenergetics. 2012;1817(2):312–8.

23. Yi JT, Wang P, Stuchebrukhov AA. Mutation at the entrance of the quinone cavity severely disrupts quinone binding in respiratory complex I. Scientific Reports. 2023;13(1).

24. Kim H, Saura P, Pöverlein MC, Gamiz-Hernandez AP, Kaila VRI. Quinone Catalysis Modulates Proton Transfer Reactions in the Membrane Domain of Respiratory Complex I. Journal of the American Chemical Society. 2023;145(31):17075–86.

25. Grandhi S, Bosworth C, Maddox W, Sensiba C, Akhavanfard S, Ni Y, et al. Heteroplasmic shifts in tumor mitochondrial genomes reveal tissue-specific signals of relaxed and positive selection. Human Molecular Genetics. 2017;26(15):2912–22.

26. Ju YS, Alexandrov LB, Gerstung M, Martincorena I, Nik-Zainal S, Ramakrishna M, et al. Origins and functional consequences of somatic mitochondrial DNA mutations in human cancer. eLife. 2014;3.

27. Kim M, Mahmood M, Reznik E, Gammage PA. Mitochondrial DNA is a major source of driver mutations in cancer. Trends in Cancer. 2022;8(12):1046–59.

28. Sanchez-Contreras M, Sweetwyne MT, Kohrn BF, Tsantilas Kristine A, Hipp Michael J, Schmidt Elizabeth K, et al. A replication-linked mutational gradient drives somatic mutation accumulation and influences germline polymorphisms and genome composition in mitochondrial DNA. Nucleic Acids Research. 2021;49(19):11103–18.

29. McCormick EM, Lott MT, Dulik MC, Shen L, Attimonelli M, Vitale O, et al. Specifications of the ACMG/AMP standards and guidelines for mitochondrial DNA variant interpretation. Human Mutation. 2020;41(12):2028–57.

30. Jin S-G, Meng Y, Johnson J, Szabó PE, Pfeifer GP. Concordance of hydrogen peroxide–induced 8-oxo-guanine patterns with two cancer mutation signatures of upper GI tract tumors. Science Advances. 2022;8(22).

31. Thibodeau ML, Zhao EY, Reisle C, Ch’ng C, Wong H-L, Shen Y, et al. Base excision repair deficiency signatures implicate germline and somatic MUTYH aberrations in pancreatic ductal adenocarcinoma and breast cancer oncogenesis. Molecular Case Studies. 2019;5(2).

32. Kampjut D, Sazanov LA. Structure of respiratory complex I – An emerging blueprint for the mechanism. Current Opinion in Structural Biology. 2022;74.

33. Röpke M, Riepl D, Saura P, Di Luca A, Mühlbauer ME, Jussupow A, et al. Deactivation blocks proton pathways in the mitochondrial complex I. Proceedings of the National Academy of Sciences. 2021;118(29).

34. Kravchuk V, Petrova O, Kampjut D, Wojciechowska-Bason A, Breese Z, Sazanov L. A universal coupling mechanism of respiratory complex I. Nature. 2022;609(7928):808–14.

35. Salomonsson L, Brändén G, Brzezinski P. Deuterium isotope effect of proton pumping in cytochrome c oxidase. Biochimica et Biophysica Acta (BBA) - Bioenergetics. 2008;1777(4):343–50.

36. Johansson A-L, Chakrabarty S, Berthold CL, Högbom M, Warshel A, Brzezinski P. Proton-transport mechanisms in cytochrome c oxidase revealed by studies of kinetic isotope effects. Biochimica et Biophysica Acta (BBA) - Bioenergetics. 2011;1807(9):1083–94.

37. Salomonsson L, Faxén K, Ädelroth P, Brzezinski P. The timing of proton migration in membrane-reconstituted cytochrome c oxidase. Proceedings of the National Academy of Sciences. 2005;102(49):17624–9.

38. Wikstro m M, Krab K, Sharma V. Oxygen Activation and Energy Conservation by Cytochrome c Oxidase. Chemical Reviews. 2018;118(5):2469–90.

39. Wong JE, Zíková A, Gahura O, Ozkan B. The Ancestral Shape of the Access Proton Path of Mitochondrial ATP Synthases Revealed by a Split Subunit-a. Molecular Biology and Evolution. 2023;40(6).

40. Ro pke M, Saura P, Riepl D, Po verlein MC, Kaila VRI. Functional Water Wires Catalyze Long-Range Proton Pumping in the Mammalian Respiratory Complex I. Journal of the American Chemical Society. 2020;142(52):21758–66.

41. Fowler DM, Fields S. Deep mutational scanning: a new style of protein science. Nature Methods. 2014;11(8):801–7.

42. Hopf TA, Ingraham JB, Poelwijk FJ, Schärfe CPI, Springer M, Sander C, et al. Mutation effects predicted from sequence co-variation. Nature Biotechnology. 2017;35(2):128–35.

43. Warshel A, Russell ST. Calculations of electrostatic interactions in biological systems and in solutions. Quarterly Reviews of Biophysics. 2009;17(3):283–422.

44. Bystroff C, Oatley SJ, Kraut J. Crystal structures of Escherichia coli dihydrofolate reductase: the NADP+ holoenzyme and the folate .cnt NADP+ ternary complex. substrate binding and a model for the transition state. Biochemistry. 2002;29(13):3263–77.

45. del Sol A, Fujihashi H, Amoros D, Nussinov R. Residues crucial for maintaining short paths in network communication mediate signaling in proteins. Molecular Systems Biology. 2006;2(1).

46. Zamyatnin AA. Protein volume in solution. Progress in Biophysics and Molecular Biology. 1972;24:107–23.

47. Nguyen MD, Sierra-Magro A, Singh V, Khawaja A, Timón-Gómez A, Barrientos A, et al. Structural basis for late maturation steps of mitochondrial respiratory chain complex IV within the human respirasome. Nature Communications. 2026;17(1).

48. Andrews RM, Kubacka I, Chinnery PF, Lightowlers RN, Turnbull DM, Howell N. Reanalysis and revision of the Cambridge reference sequence for human mitochondrial DNA. Nature Genetics. 1999;23(2):147-.

49. Needleman SB, Wunsch CD. A general method applicable to the search for similarities in the amino acid sequence of two proteins. Journal of Molecular Biology. 1970;48(3):443–53.

50. Henikoff S, Henikoff JG. Amino acid substitution matrices from protein blocks. Proceedings of the National Academy of Sciences. 1992;89(22):10915–9.

51. Zdorevskyi O, Djurabekova A, Lasham J, Sharma V. Horizontal proton transfer across the antiporter-like subunits in mitochondrial respiratory complex I. Chemical Science. 2023;14(23):6309–18.

52. Parey K, Lasham J, Mills DJ, Djurabekova A, Haapanen O, Yoga EG, et al. High-resolution structure and dynamics of mitochondrial complex I—Insights into the proton pumping mechanism. Science Advances. 2021;7(46).

53. Aaltonen LA, Abascal F, Abeshouse A, Aburatani H, Adams DJ, Agrawal N, et al. Pan-cancer analysis of whole genomes. Nature. 2020;578(7793):82–93.

54. Cheng DT, Mitchell TN, Zehir A, Shah RH, Benayed R, Syed A, et al. Memorial Sloan Kettering-Integrated Mutation Profiling of Actionable Cancer Targets (MSK-IMPACT). The Journal of Molecular Diagnostics. 2015;17(3):251–64.

55. Zehir A, Benayed R, Shah RH, Syed A, Middha S, Kim HR, et al. Mutational landscape of metastatic cancer revealed from prospective clinical sequencing of 10,000 patients. Nature Medicine. 2017;23(6):703–13.

