## Supplementary code for "Cancers relax selection on charge at the mitochondrial complex I quinone cavity": config.py

```
"""Paths and analysis parameters."""
from pathlib import Path

ROOT = Path(__file__).resolve().parent
DATA = ROOT / "data"
RESULTS = ROOT / "results"
FIGURES = ROOT / "figures"
RESULTS.mkdir(exist_ok=True)
FIGURES.mkdir(exist_ok=True)

# --- input files ---------------------------------------------------------
MOUSE_CIF = DATA / "8OM1.cif"
HUMAN_CIF = DATA / "9TI4.cif"
RCRS_FASTA = DATA / "rCRS.fasta"
MTDNA = DATA / "mtdna-mutations" / "data"
NUCLEAR_MAFS = DATA / "tcga_nuclear_mafs"
COSMIC = DATA / "COSMIC_v3.4_SBS_GRCh38_exome.txt"

# --- structure -----------------------------------------------------------
MOUSE_CHAINS = {"H": "ND1", "N": "ND2", "A": "ND3", "M": "ND4",
                "K": "ND4L", "L": "ND5", "J": "ND6"}
HUMAN_CHAINS = {"s": "ND1", "i": "ND2", "j": "ND3", "r": "ND4",
                "k": "ND4L", "l": "ND5", "m": "ND6"}
SUBUNITS = ("ND1", "ND2", "ND3", "ND4", "ND4L", "ND5", "ND6")
LIPIDS = {"3PE", "CDL", "PC1", "LMT", "MYR", "PEE", "PLC"}

# --- proton-wire graph ---------------------------------------------------
# Grotthuss-competent side-chain atoms. Backbone N and O are excluded by
# construction: the amide proton does not exchange on a catalytic timescale,
# and including backbone atoms saturates the network through alpha-helical
# hydrogen bonding.
WIRE_ATOMS = {
    "SER": {"OG"}, "THR": {"OG1"}, "TYR": {"OH"}, "CYS": {"SG"},
    "ASP": {"OD1", "OD2"}, "GLU": {"OE1", "OE2"},
    "ASN": {"OD1", "ND2"}, "GLN": {"OE1", "NE2"},
    "HIS": {"ND1", "NE2"}, "LYS": {"NZ"},
    "ARG": {"NE", "NH1", "NH2"}, "TRP": {"NE1"},
}
TITRATABLE = {"ASP", "GLU", "HIS", "LYS", "ARG", "TYR", "CYS"}

HBOND_MIN = 2.2            # A, shortest admissible donor-acceptor distance
HBOND_CUTOFF = 3.5         # A, primary cutoff
HBOND_SWEEP = (3.2, 3.5, 4.0)
WATER_SHELL = 12.0         # A, waters kept within this distance of an ND atom
LINING_SHELL = 6.0         # A, non-ND side chains kept within this distance
MIN_CLUSTER = 4            # nodes, smallest hydrated cluster considered
MAJOR_MIN_NODES = 12
MAJOR_MIN_SPAN = 10.0      # A along the membrane normal

# --- axis A, electrostatics ---------------------------------------------
DIELECTRIC = 8.0
R_FLOOR = 3.0              # A
CHARGE = {"ASP": -1.0, "GLU": -1.0, "LYS": 1.0, "ARG": 1.0, "HIS": 0.1}

# --- axis B, connectivity ------------------------------------------------
STERIC_VOLUME_THRESHOLD = 20.0   # A^3
STERIC_WATER_SHELL = 5.0         # A

# --- thresholds used to define the off-diagonal classes ------------------
AXIS_A_THRESHOLD = 0.02    # eV
AXIS_B_THRESHOLD = 1.0

# --- variant analysis ----------------------------------------------------
# Coding intervals on the rCRS (1-based inclusive); ND6 is on the light strand.
ND_GENES = {
    "ND1":  (3307, 4262, 1),
    "ND2":  (4470, 5511, 1),
    "ND3":  (10059, 10404, 1),
    "ND4L": (10470, 10766, 1),
    "ND4":  (10760, 12137, 1),
    "ND5":  (12337, 14148, 1),
    "ND6":  (14149, 14673, -1),
}
ND_PROTEIN_LENGTHS = {"ND1": 318, "ND2": 347, "ND3": 115, "ND4": 459,
                      "ND4L": 98, "ND5": 603, "ND6": 174}
MITO_CODE = 2              # vertebrate mitochondrial genetic code
POLYMORPHISM_CARRIERS = 10  # somatic calls above this HelixMTdb count are dropped
SPECTRUM_PSEUDOCOUNT = 0.5
N_CHANNELS = 192           # strand-aware trinucleotide channels
VAF_SPLIT = 0.5
N_PERMUTATIONS = 10_000
RANDOM_SEED = 0

COHORTS = ("germline", "TCGA", "PCAWG", "MSK-IMPACT")
WIRE_CLASSES = ("loading_site", "cut_vertex", "cluster_member", "other")
```
