## Supplementary code for "Cancers relax selection on charge at the mitochondrial complex I quinone cavity": README.md

# Selection on the complex I proton wire

Analysis code for *Cancers relax selection on charge at the mitochondrial
complex I quinone cavity*.

The pipeline builds the hydrogen-bonded proton wire of the seven mtDNA-encoded
subunits of respiratory complex I from a cryo-EM structure, scores every
possible missense substitution on two physically separable axes, and tests
whether the resulting residue classes are under differential selection in the
human germline and in tumours.

## Requirements

Python 3.10 or later.

```
pip install -r requirements.txt
```

Analyses were run with Python 3.13, numpy 2.3, pandas 2.3, scipy 1.16,
networkx 3.5, statsmodels 0.14, biopython 1.88 and gemmi 0.7.

## Input data

Place the following under `data/` (see `config.py` to change the location).

| File | Source |
|---|---|
| `8OM1.cif` | RCSB PDB — mouse complex I, active state, 2.39 Å |
| `9TI4.cif` | RCSB PDB — human complex I (used only as a fold control) |
| `rCRS.fasta` | NCBI NC_012920.1, the revised Cambridge Reference Sequence |
| `mtdna-mutations/` | Gorelick et al., <https://github.com/reznik-lab/mtdna-mutations> |
| `tcga_nuclear_mafs/` | Open-access ensemble-masked MAF files, NCI Genomic Data Commons |
| `COSMIC_v3.4_SBS_GRCh38_exome.txt` | COSMIC v3.4 reference signatures |

The Gorelick repository supplies the germline (HelixMTdb), TCGA, PCAWG and
MSK-IMPACT mtDNA variant calls, the tumour purity tables and MitImpact v2.9.

## Running

```
bash run_all.sh
```

or step by step:

| Script | Produces |
|---|---|
| `scripts/01_build_wire.py` | proton-wire graph, hydrated clusters, residue classes |
| `scripts/02_map_to_human.py` | rCRS translation, mouse–human alignment, codon map |
| `scripts/03_score_axes.py` | axis A (charge) and axis B (connectivity) per substitution |
| `scripts/04_universe.py` | complete substitution universe and mutational null model |
| `scripts/05_selection.py` | observed/expected ratios, quasi-Poisson regressions, enrichment by subunit |
| `scripts/06_heteroplasmy_disease.py` | heteroplasmy strata, disease-variant enrichment and overlap |
| `scripts/07_nuclear.py` | nuclear SBS96 profiles, COSMIC fitting, oxidative readouts |
| `scripts/08_sensitivity.py` | cutoff, filter and axis-B sensitivity analyses |
| `scripts/09_figures.py` | Figure 1, Figure 3 and Supplementary Figures 1-6 |
| `scripts/10_recurrence.py` | recurrence of individual substitutions, clonality |
| `scripts/11_haplogroup.py` | PhyloTree branch-defining variants, the three timescales |
| `scripts/12_local_structure.py` | per-residue mouse–human structural concordance |
| `scripts/13_interaction.py` | the wire class by tissue-type interaction test |
| `scripts/14_figure_s7.py` | Supplementary Figure 7 |
| `scripts/15_coverage.py` | sequencing-coverage control |
| `scripts/16_strata.py` | cancer-type stratification, PCAWG allele-fraction bands |
| `scripts/17_jackknife.py` | leave-one-gene-out |
| `scripts/18_axis_sensitivity.py` | axis A dielectric and distance floor |
| `scripts/19_multiple_testing.py` | one Benjamini-Hochberg table for the paper |
| `scripts/20_figure_s8.py` | Supplementary Figure 8 |
| `scripts/21_structural_context.py` | position in the enzyme: quinone cavity, central axis |
| `scripts/22_figure_s9.py` | Supplementary Figure 9 |
| `scripts/23_somatic_support.py` | matched-normal support and clonality of recurrent calls |
| `scripts/24_quinone_robustness.py` | radius sweep, distance gradient, cut-vertex bound |
| `scripts/25_figure_s10.py` | Supplementary Figure 10 |
| `scripts/26_main_figures.py` | Figure 2 and Figure 4 |

Intermediates are written to `results/`, figures to `figures/` and the console
output of each step to `logs/`. The scripts are ordered; later steps read what
earlier ones write. `run_all.sh` covers steps 01-09; the remaining steps are
run in order afterwards. Figures 2 and 4 are built last, by
`scripts/26_main_figures.py`, because they use the interaction test of step 13
and the quinone analyses of steps 21 and 24.

The tables behind the reported numbers are

| File in `results/` | Contents |
|---|---|
| `residue_classes.csv` | every modelled ND residue with its wire class and burial |
| `hydrated_clusters.csv` | size, water count and membrane span of each cluster |
| `graph_strategies.csv`, `transferred_water_clashes.csv` | the cross-species comparison |
| `mouse_human_map.csv` | mouse residue to human codon, with the identity flag |
| `axis_scores.parquet`, `separability.json` | both axes for every substitution |
| `universe.parquet`, `universe_annotated.parquet` | the substitution universe and its covariates |
| `oe_synonymous.csv`, `oe_missense.csv` | observed/expected by class and cohort |
| `rate_ratios.csv` | the covariate ladder |
| `oe_by_subunit.csv` | loading-site enrichment within each ND subunit |
| `interaction.csv` | the wire class by tissue-type interaction |
| `recurrent_variants.csv` | substitutions of the two axis classes seen in four or more patients |
| `disease_overlap.csv` | overlap between reported-pathogenic and tumour-observed substitutions |
| `heteroplasmy.csv` | enrichment by heteroplasmy stratum |
| `disease_enrichment.csv`, `axis_a_by_group.csv` | disease variants against the wire |
| `nuclear_signatures.csv`, `nuclear_signature_shares.csv` | the oxidative readout |
| `sensitivity_*.csv` | the four sensitivity analyses |
| `recurrence.csv`, `clonality.csv`, `recurrent_variants.csv` | recurrence and allele fractions |
| `haplogroup_enrichment.csv`, `timescales.csv`, `axis_a_by_timescale.csv` | the evolutionary comparison |
| `structural_concordance.csv`, `oe_missense_concordant.csv`, `filter_comparison.csv` | cross-species concordance |
| `interaction.csv` | the interaction test |

Figures read only these tables, so a panel cannot disagree with the text.

## Notes on method

Backbone amide and carbonyl atoms are excluded from the wire graph. They are
not Grotthuss-competent, and including them makes the network continuous across
the membrane at 2.85 Å in every module — an artefact of α-helical hydrogen
bonding rather than a proton pathway.

The wire geometry comes from the mouse structure because no human structure of
comparable resolution models hydration. `scripts/01_build_wire.py` reports the
comparison that justifies this: the human structure alone yields 292 wire edges
against 2,006, and mouse waters superposed into the human frame clash with a
human heavy atom in 2,042 of the 2,945 deposited waters (828 of the 1,329 that
lie within 12 A of an ND atom).

Residue classes used for the selection analyses are derived from the graph
alone. Axis B enters only the separability diagnostic of Figure 1e.

Hydrogen-bond cutoff, species filter, polymorphism threshold and the definition
of axis B are the four discretionary choices in the pipeline;
`scripts/08_sensitivity.py` rebuilds the analysis under each alternative. At a
4.0 Å cutoff the cut-vertex class becomes depleted in tumours rather than
unchanged, which is why the manuscript claims only that it is never enriched.

## Licence

MIT.
