## Supplementary code for "Cancers relax selection on charge at the mitochondrial complex I quinone cavity": run_all.sh

```
#!/usr/bin/env bash
# Run the whole analysis in order. Each script reads what the previous ones
# wrote into results/, so the order matters.
set -euo pipefail

cd "$(dirname "$0")"
mkdir -p results figures logs

for script in scripts/0[1-9]_*.py; do
    name=$(basename "$script" .py)
    echo "--- ${name}"
    python -u "$script" 2>&1 | tee "logs/${name}.log"
done

echo "--- done; tables in results/, figures in figures/"
```
