## Supplementary code for "Cancers relax selection on charge at the mitochondrial complex I quinone cavity": 01_build_wire.py

```
"""Build the proton wire and classify the mtDNA-encoded residues.

Also records the comparison that justifies taking the wire geometry from the
mouse structure rather than a human one.
"""
import pickle
import sys
from pathlib import Path

import numpy as np
import pandas as pd
from scipy.spatial import cKDTree

sys.path.insert(0, str(Path(__file__).resolve().parents[1]))
import config as cfg
from src import wire


def kabsch(P, Q):
    """Rotation and translation taking P onto Q."""
    cp, cq = P.mean(0), Q.mean(0)
    U, _, Vt = np.linalg.svd((P - cp).T @ (Q - cq))
    d = np.sign(np.linalg.det(Vt.T @ U.T))
    R = Vt.T @ np.diag([1.0, 1.0, d]) @ U.T
    return R, cq - R @ cp


def main():
    model = wire.load_model(cfg.MOUSE_CIF, cfg.MOUSE_CHAINS)
    centre, normal, in_plane, lipids = wire.membrane_frame(model)
    side = wire.matrix_side(model, centre, normal)

    nodes = wire.collect_nodes(model, cfg.MOUSE_CHAINS)
    graph, pairs, distances, X = wire.build_graph(nodes, cfg.HBOND_CUTOFF)
    proj_normal = (X - centre) @ normal
    proj_plane = (X - centre) @ in_plane
    classes, comps, major, articulation = wire.classify(nodes, graph, proj_normal)
    geometry = wire.residue_geometry(model, cfg.MOUSE_CHAINS)

    counts = pd.Series(list(classes.values())).value_counts()
    n_waters = sum(1 for n in nodes if n[1] == "HOH")
    low, high = np.percentile((lipids - centre) @ normal, [5, 95])

    print(f"modelled ND residues      {len(geometry)}")
    print(f"wire nodes                {len(nodes)} ({n_waters} waters)")
    print(f"wire edges at {cfg.HBOND_CUTOFF} A      {len(pairs)}")
    print(f"hydrated clusters         {len(comps)} ({len(major)} major)")
    for name in ("loading_site", "cut_vertex", "cluster_member"):
        print(f"  {name:15s} {int(counts.get(name, 0))}")
    print(f"bilayer on the normal     {low:.1f} to {high:.1f} A "
          f"(matrix on the {'+' if side > 0 else '-'} side)")

    # Is there a continuous path across the membrane in this static structure?
    n_face = proj_normal > high - 3.0 if side > 0 else proj_normal < low + 3.0
    p_face = proj_normal < low + 3.0 if side > 0 else proj_normal > high - 3.0
    import networkx as nx
    reachable = set()
    for component in nx.connected_components(graph):
        if any(n_face[i] for i in component) and any(p_face[i] for i in component):
            reachable |= component
    print(f"continuous N-to-P wire    {'yes' if reachable else 'no'}")

    # Why the mouse structure: the human one models almost no water, and mouse
    # waters cannot simply be transferred into the human frame.
    human = wire.load_model(cfg.HUMAN_CIF, cfg.HUMAN_CHAINS)
    human_nodes = [n for n in wire.collect_nodes(human, cfg.HUMAN_CHAINS)
                   if n[1] != "HOH"]
    _, human_pairs, _, _ = wire.build_graph(human_nodes, cfg.HBOND_CUTOFF)

    def ca(m, chains):
        out = {}
        for ch in m:
            if ch.name not in chains:
                continue
            for r in ch:
                a = r.find_atom("CA", "*")
                if a:
                    out[(chains[ch.name], r.seqid.num)] = [a.pos.x, a.pos.y, a.pos.z]
        return out

    mouse_ca, human_ca = ca(model, cfg.MOUSE_CHAINS), ca(human, cfg.HUMAN_CHAINS)
    shared = sorted(set(mouse_ca) & set(human_ca))
    P = np.asarray([mouse_ca[k] for k in shared])
    Q = np.asarray([human_ca[k] for k in shared])
    R, t = kabsch(P, Q)
    rmsd = float(np.sqrt((((P @ R.T + t) - Q) ** 2).sum(1).mean()))

    waters = np.asarray([n[3] for n in nodes if n[1] == "HOH"]) @ R.T + t
    human_heavy = np.asarray([[a.pos.x, a.pos.y, a.pos.z] for ch in human
                              for r in ch if r.het_flag != "H" for a in r])
    tree = cKDTree(human_heavy)
    clash = tree.query(waters)[0]

    # The same test over every water in the deposited model, not only those
    # within WATER_SHELL of an ND atom.
    deposited = np.asarray([[a.pos.x, a.pos.y, a.pos.z] for ch in model
                            for r in ch if r.name == "HOH" for a in r])
    clash_all = tree.query(deposited @ R.T + t)[0]

    print(f"\nhuman structure, side chains only: {len(human_pairs)} wire edges")
    print(f"superposition on {len(shared)} CA atoms: RMSD {rmsd:.2f} A")
    print(f"transferred waters clashing below {cfg.HBOND_MIN} A: "
          f"{int((clash < cfg.HBOND_MIN).sum())} of {len(clash)} within "
          f"{cfg.WATER_SHELL:.0f} A of an ND atom, "
          f"{int((clash_all < cfg.HBOND_MIN).sum())} of {len(clash_all)} "
          f"over the whole deposited model")

    pd.DataFrame([{"subunit": k[0], "pos": k[1], "residue": geometry[k]["residue"],
                   "burial": geometry[k]["burial"],
                   "wire_class": classes.get(k, "other")}
                  for k in sorted(geometry)]).to_csv(
        cfg.RESULTS / "residue_classes.csv", index=False)

    pd.DataFrame([{"cluster": k, "n_nodes": len(c),
                   "n_waters": sum(1 for i in c if nodes[i][1] == "HOH"),
                   "span_normal": float(np.ptp(proj_normal[list(c)])),
                   "major": k in major} for k, c in enumerate(comps)]).to_csv(
        cfg.RESULTS / "hydrated_clusters.csv", index=False)

    pd.DataFrame({"min_distance": clash}).to_csv(
        cfg.RESULTS / "transferred_water_clashes.csv", index=False)

    # Third strategy: human side chains plus the mouse waters that survive the
    # superposition without a steric clash.
    kept = waters[clash >= cfg.HBOND_MIN]
    transferred = human_nodes + [(("HOH", i), "HOH", "O", xyz)
                                 for i, xyz in enumerate(kept)]
    _, transferred_pairs, _, _ = wire.build_graph(transferred, cfg.HBOND_CUTOFF)

    pd.DataFrame([
        {"strategy": "mouse structure, modelled waters",
         "n_edges": len(pairs), "n_waters": n_waters},
        {"strategy": "human structure, side chains only",
         "n_edges": len(human_pairs), "n_waters": 0},
        {"strategy": "human structure, mouse waters transferred",
         "n_edges": len(transferred_pairs), "n_waters": len(kept)},
    ]).to_csv(cfg.RESULTS / "graph_strategies.csv", index=False)
    print(f"human structure with {len(kept)} transferable mouse waters: "
          f"{len(transferred_pairs)} wire edges")

    with open(cfg.RESULTS / "wire.pkl", "wb") as fh:
        pickle.dump({"nodes": [(n[0], n[1], n[2], n[3].tolist()) for n in nodes],
                     "graph": graph, "classes": classes, "comps": comps,
                     "major": major, "geometry": geometry, "X": X,
                     "proj_normal": proj_normal, "proj_plane": proj_plane,
                     "centre": centre, "normal": normal, "in_plane": in_plane,
                     "lipid_xyz": lipids, "slab": (low, high),
                     "matrix_side": side}, fh)
    print(f"\nwrote {cfg.RESULTS / 'wire.pkl'}")


if __name__ == "__main__":
    main()
```
