## Supplementary code for "Cancers relax selection on charge at the mitochondrial complex I quinone cavity": 02_map_to_human.py

```
"""Translate the ND genes from the rCRS and map the mouse classification onto
human codons.

The wire geometry is mouse; the variant data are human. Positions at which the
two residues differ are flagged, and the primary analyses are restricted to
those that are identical, where the structural inference transfers directly.
"""
import pickle
import sys
from pathlib import Path

import gemmi
import pandas as pd
from Bio import Align
from Bio.Align import substitution_matrices

sys.path.insert(0, str(Path(__file__).resolve().parents[1]))
import config as cfg
from src import variants, wire


def mouse_sequences(model, chains):
    """Modelled sequence and residue numbers for each ND chain."""
    out = {}
    for ch in model:
        if ch.name not in chains:
            continue
        letters, numbers = [], []
        for r in ch:
            if r.het_flag == "H" and r.name != "FME":
                continue
            name = "MET" if r.name == "FME" else r.name
            info = gemmi.find_tabulated_residue(name)
            if info is None or not info.is_amino_acid():
                continue
            letters.append(info.one_letter_code.upper())
            numbers.append(r.seqid.num)
        out[chains[ch.name]] = ("".join(letters), numbers)
    return out


def main():
    with open(cfg.RESULTS / "wire.pkl", "rb") as fh:
        built = pickle.load(fh)
    classes = built["classes"]

    seq = variants.read_rcrs(cfg.RCRS_FASTA)
    proteins, codons = variants.translate_nd_genes(seq)
    print("translation in the vertebrate mitochondrial code")
    for gene in cfg.ND_GENES:
        print(f"  {gene:5s} {len(proteins[gene]):4d} aa")

    model = wire.load_model(cfg.MOUSE_CIF, cfg.MOUSE_CHAINS)
    mouse = mouse_sequences(model, cfg.MOUSE_CHAINS)

    aligner = Align.PairwiseAligner(
        mode="global", open_gap_score=-11, extend_gap_score=-1,
        substitution_matrix=substitution_matrices.load("BLOSUM62"))

    rows = []
    print("\nmouse to human alignment")
    for gene in cfg.ND_GENES:
        mouse_seq, numbers = mouse[gene]
        human_seq = proteins[gene]
        alignment = aligner.align(mouse_seq, human_seq)[0]
        identical = 0
        mapping = {}
        for (ms, me), (hs, _) in zip(*alignment.aligned):
            for k in range(me - ms):
                mi, hi = ms + k, hs + k
                mapping[numbers[mi]] = (hi + 1, mouse_seq[mi], human_seq[hi])
                identical += mouse_seq[mi] == human_seq[hi]
        print(f"  {gene:5s} mouse {len(mouse_seq):4d}  human {len(human_seq):4d}  "
              f"identity {100 * identical / len(mouse_seq):5.1f}%")
        for number in numbers:
            hit = mapping.get(number)
            rows.append({
                "subunit": gene,
                "mouse_pos": number,
                "mouse_aa": hit[1] if hit else None,
                "human_pos": hit[0] if hit else None,
                "human_aa": hit[2] if hit else None,
                "identical": bool(hit and hit[1] == hit[2]),
                "transferable": hit is not None,
                "wire_class": classes.get((gene, number), "other"),
                "codon_start": codons[gene][hit[0] - 1][0] if hit else None,
                "codon_end": codons[gene][hit[0] - 1][2] if hit else None,
            })

    table = pd.DataFrame(rows)
    wire_rows = table[table.wire_class != "other"]
    print(f"\nwire residues {len(wire_rows)}, identical in both species "
          f"{int(wire_rows.identical.sum())} "
          f"({100 * wire_rows.identical.mean():.1f}%)")
    for name in ("loading_site", "cut_vertex", "cluster_member"):
        s = wire_rows[wire_rows.wire_class == name]
        print(f"  {name:15s} {int(s.identical.sum())}/{len(s)}")

    table.to_csv(cfg.RESULTS / "mouse_human_map.csv", index=False)
    with open(cfg.RESULTS / "translation.pkl", "wb") as fh:
        pickle.dump({"proteins": proteins, "codons": codons}, fh)
    print(f"\nwrote {cfg.RESULTS / 'mouse_human_map.csv'}")


if __name__ == "__main__":
    main()
```
