## Supplementary code for "Cancers relax selection on charge at the mitochondrial complex I quinone cavity": 03_score_axes.py

```
"""Score every possible ND missense substitution on both axes.

Axis A is the electrostatic shift at a proton loading site; axis B is the
disruption of wire connectivity. The two are computed from different
quantities, so a charge-neutral substitution can score on B alone and a distant
charge change on A alone.
"""
import json
import pickle
import sys
from pathlib import Path

import numpy as np

sys.path.insert(0, str(Path(__file__).resolve().parents[1]))
import config as cfg
from src import axes


def main():
    with open(cfg.RESULTS / "wire.pkl", "rb") as fh:
        built = pickle.load(fh)

    nodes = [(k, rn, an, np.asarray(xyz)) for k, rn, an, xyz in built["nodes"]]
    graph, classes, geometry = built["graph"], built["classes"], built["geometry"]

    metrics = axes.wire_metrics(nodes, graph, classes)
    loading_xyz = axes.loading_site_geometry(classes, geometry)

    major_nodes = {i for k in built["major"] for i in built["comps"][k]}
    cluster_water_xyz = np.asarray([nodes[i][3] for i in sorted(major_nodes)
                                    if nodes[i][1] == "HOH"])

    scores = axes.score_substitutions(geometry, metrics, loading_xyz,
                                      cluster_water_xyz)
    print(f"scored {len(scores):,} possible missense substitutions at "
          f"{scores.groupby(['subunit', 'pos']).ngroups:,} residues")
    print(f"proton loading sites used for axis A: {len(loading_xyz)}")

    full = axes.separability(scores, "axis_b_connectivity")
    topology = axes.separability(scores, "axis_b_topology_only")
    print("\nseparability of the two axes")
    print(f"  Pearson r, all substitutions        {full['pearson_r_all']:.3f}")
    print(f"  Pearson r, charge-changing only     "
          f"{full['pearson_r_charge_changing']:.3f}")
    print(f"  connectivity-only substitutions     {full['n_connectivity_only']:,} "
          f"at {full['residues_connectivity_only']} residues")
    print(f"  charge-only substitutions           {full['n_charge_only']:,} "
          f"at {full['residues_charge_only']} residues")
    print("\nwith the steric term of axis B removed")
    print(f"  Pearson r, all substitutions        {topology['pearson_r_all']:.3f}")
    print(f"  connectivity-only substitutions     "
          f"{topology['n_connectivity_only']:,}")

    scores.to_parquet(cfg.RESULTS / "axis_scores.parquet", index=False)
    with open(cfg.RESULTS / "separability.json", "w") as fh:
        json.dump({"with_steric_term": full, "topology_only": topology}, fh, indent=2)
    print(f"\nwrote {cfg.RESULTS / 'axis_scores.parquet'}")


if __name__ == "__main__":
    main()
```
