## Supplementary code for "Cancers relax selection on charge at the mitochondrial complex I quinone cavity": 04_universe.py

```
"""Enumerate the substitution universe and attach the observed variant counts.

The universe is built from first principles by translating every mutant codon,
rather than taken from a missense-only annotation database, so that the
synonymous set used to calibrate the null model is unbiased with respect to
codon position.
"""
import collections
import pickle
import sys
from pathlib import Path

import pandas as pd

sys.path.insert(0, str(Path(__file__).resolve().parents[1]))
import config as cfg
from src import variants

COHORT_FILES = {
    "germline": "data_mutations_helix.txt.gz",
    "TCGA": "data_mutations_tcga.txt.gz",
    "PCAWG": "data_mutations_pcawg.txt.gz",
    "MSK-IMPACT": "data_mutations_impact.txt.gz",
}


def load_cohorts():
    return {name: pd.read_csv(cfg.MTDNA / "original_data" / fn, sep="\t",
                              low_memory=False)
            for name, fn in COHORT_FILES.items()}


def main():
    seq = variants.read_rcrs(cfg.RCRS_FASTA)
    with open(cfg.RESULTS / "translation.pkl", "rb") as fh:
        translation = pickle.load(fh)

    mapping = pd.read_csv(cfg.RESULTS / "mouse_human_map.csv")
    mapping = mapping[mapping.transferable]
    human_class = {(r.subunit, int(r.human_pos)): r.wire_class
                   for r in mapping.itertuples()}
    human_identical = {(r.subunit, int(r.human_pos)): bool(r.identical)
                       for r in mapping.itertuples()}

    universe = variants.build_universe(seq, translation["proteins"],
                                       translation["codons"], human_class)
    universe["identical"] = [human_identical.get((s, p), False)
                             for s, p in zip(universe.subunit, universe.aa_pos)]
    universe["gene"] = "MT-" + universe.subunit
    universe["key"] = list(zip(universe.pos, universe.ref, universe.alt))

    print(f"substitution universe {len(universe):,} over "
          f"{universe.groupby(['subunit', 'aa_pos']).ngroups:,} codons")
    print(universe.kind.value_counts().to_string())

    cohorts = load_cohorts()
    nd_genes = sorted(universe.gene.unique())
    spectra = {}
    print("\nobserved ND variants and spectrum size")
    for name, calls in cohorts.items():
        spectra[name] = variants.trinucleotide_spectrum(calls, seq, nd_genes)
        for kind in ("missense", "synonymous"):
            counts = variants.observed_counts(calls, nd_genes, kind)
            column = f"n_{name}_{kind}"
            universe[column] = universe.key.map(counts).fillna(0).astype(int)
            universe.loc[universe.kind != kind, column] = 0
        print(f"  {name:11s} missense {int(universe[f'n_{name}_missense'].sum()):5d}  "
              f"synonymous {int(universe[f'n_{name}_synonymous'].sum()):5d}")

    # Somatic calls at common germline sites are population polymorphisms.
    helix = cohorts["germline"][["Start_Position", "Reference_Allele",
                                 "Tumor_Seq_Allele2", "TOTAL_COUNT"]].copy()
    helix["TOTAL_COUNT"] = pd.to_numeric(helix.TOTAL_COUNT, errors="coerce").fillna(0)
    carriers = collections.defaultdict(float)
    for pos, ref, alt, total in helix.itertuples(index=False):
        carriers[(pos, ref, alt)] = max(carriers[(pos, ref, alt)], total)
    universe["polymorphism"] = (universe.key.map(carriers).fillna(0)
                                > cfg.POLYMORPHISM_CARRIERS)
    print(f"\nsites excluded from the somatic analysis as polymorphisms: "
          f"{int(universe.polymorphism.sum())}")

    # The tuple key is rebuilt on read; parquet cannot store it.
    universe.drop(columns="key").to_parquet(
        cfg.RESULTS / "universe.parquet", index=False)
    with open(cfg.RESULTS / "spectra.pkl", "wb") as fh:
        pickle.dump(spectra, fh)
    print(f"wrote {cfg.RESULTS / 'universe.parquet'}")


if __name__ == "__main__":
    main()
```
