## Supplementary code for "Cancers relax selection on charge at the mitochondrial complex I quinone cavity": 05_selection.py

```
"""Selection on the proton-wire classes.

Observed/expected ratios against each cohort's own mutational spectrum, the
synonymous calibration of that null, and quasi-Poisson regressions adding
conservation and structural covariates in a fixed ladder.
"""
import pickle
import sys
from pathlib import Path

import numpy as np
import pandas as pd
import statsmodels.api as sm
import statsmodels.formula.api as smf

sys.path.insert(0, str(Path(__file__).resolve().parents[1]))
import config as cfg
from src import variants

LADDER = [
    ("unadjusted", "y ~ C(wire_class)"),
    ("conservation", "y ~ C(wire_class) + phastcons"),
    ("conservation + structure",
     "y ~ C(wire_class) + phastcons + burial + delta_volume + delta_hydropathy"),
    ("conservation + structure + distance",
     "y ~ C(wire_class) + phastcons + burial + delta_volume + delta_hydropathy"
     " + dist_to_loading_site"),
]


def oe_table(universe, spectra, kind):
    rows = []
    for cohort in cfg.COHORTS:
        subset = universe[(universe.kind == kind) & universe.identical]
        if cohort != "germline":
            subset = subset[~subset.polymorphism]
        column = f"n_{cohort}_{kind}"
        for wire_class in cfg.WIRE_CLASSES:
            mask = (subset.wire_class == wire_class).to_numpy()
            obs, exp, oe, p = variants.observed_expected(
                subset, spectra[cohort], column, mask)
            low, high = (variants.poisson_interval(obs, exp)
                         if exp and np.isfinite(exp) else (np.nan, np.nan))
            rows.append(dict(cohort=cohort, wire_class=wire_class, observed=obs,
                             expected=round(exp, 1) if np.isfinite(exp) else np.nan,
                             obs_over_exp=round(oe, 3) if np.isfinite(oe) else np.nan,
                             ci_low=round(low, 3), ci_high=round(high, 3),
                             p_value=p))
    return pd.DataFrame(rows)


def annotate_covariates(universe):
    """Attach conservation and the structural covariates."""
    mitimpact = pd.read_csv(cfg.MTDNA / "ext" / "MitImpact_db_2.9.txt", sep="\t",
                            low_memory=False,
                            usecols=["Start", "Ref", "Alt", "PhastCons_100V"])
    phastcons = dict(zip(zip(mitimpact.Start, mitimpact.Ref, mitimpact.Alt),
                         pd.to_numeric(mitimpact.PhastCons_100V, errors="coerce")))
    universe["phastcons"] = universe.key.map(phastcons)

    scores = pd.read_parquet(cfg.RESULTS / "axis_scores.parquet")
    mapping = pd.read_csv(cfg.RESULTS / "mouse_human_map.csv")
    mapping = mapping[mapping.transferable]
    to_human = {(r.subunit, int(r.mouse_pos)): int(r.human_pos)
                for r in mapping.itertuples()}
    scores["human_pos"] = [to_human.get((s, p)) for s, p in
                           zip(scores.subunit, scores.pos)]
    scores = scores[scores.human_pos.notna()]
    lookup = {(r.subunit, int(r.human_pos), r.mut_aa):
              (r.burial, r.delta_volume, r.delta_hydropathy, r.dist_to_loading_site)
              for r in scores.itertuples()}
    hit = [lookup.get((s, p, a)) for s, p, a in
           zip(universe.subunit, universe.aa_pos, universe.mut_aa)]
    for i, name in enumerate(["burial", "delta_volume", "delta_hydropathy",
                              "dist_to_loading_site"]):
        universe[name] = [h[i] if h else np.nan for h in hit]
    return universe


def regressions(universe, spectra):
    data = universe[(universe.kind == "missense") & universe.identical
                    & universe.phastcons.notna() & universe.burial.notna()].copy()
    data["wire_class"] = pd.Categorical(
        data.wire_class, categories=list(cfg.WIRE_CLASSES)[::-1])
    rows = []
    for cohort in cfg.COHORTS:
        subset = data if cohort == "germline" else data[~data.polymorphism]
        subset = subset.copy()
        subset["y"] = subset[f"n_{cohort}_missense"]
        weights = subset.channel.map(spectra[cohort]).fillna(spectra[cohort].min())
        subset["offset"] = np.log(weights / weights.mean())
        for label, formula in LADDER:
            fit = smf.glm(formula, data=subset, family=sm.families.Poisson(),
                          offset=subset["offset"]).fit(scale="X2")
            for wire_class in ("loading_site", "cut_vertex", "cluster_member"):
                term = f"C(wire_class)[T.{wire_class}]"
                low, high = np.exp(fit.conf_int().loc[term])
                rows.append(dict(cohort=cohort, model=label, wire_class=wire_class,
                                 rate_ratio=round(float(np.exp(fit.params[term])), 3),
                                 ci_low=round(float(low), 3),
                                 ci_high=round(float(high), 3),
                                 p_value=float(fit.pvalues[term]),
                                 n_substitutions=len(subset)))
    return pd.DataFrame(rows)


def oe_by_subunit(universe, spectra, cohort="PCAWG"):
    """Loading-site observed/expected within each ND subunit.

    The expectation is taken over the subunit's own substitutions, so the
    ratio asks whether that subunit concentrates its missense variants on
    loading sites, not whether the subunit is mutated more than the rest.
    """
    rows = []
    subset = universe[(universe.kind == "missense") & universe.identical
                      & ~universe.polymorphism]
    column = f"n_{cohort}_missense"
    for subunit in sorted(subset.subunit.unique()):
        gene = subset[subset.subunit == subunit]
        mask = (gene.wire_class == "loading_site").to_numpy()
        if not mask.any():
            continue
        obs, exp, oe, p = variants.observed_expected(
            gene, spectra[cohort], column, mask)
        low, high = variants.poisson_interval(obs, exp)
        rows.append(dict(cohort=cohort, subunit=subunit, observed=obs,
                         expected=round(exp, 1),
                         obs_over_exp=round(oe, 3) if exp else np.nan,
                         ci_low=round(low, 3), ci_high=round(high, 3),
                         p_value=p))
    return pd.DataFrame(rows)


def main():
    universe = pd.read_parquet(cfg.RESULTS / "universe.parquet")
    universe["key"] = list(zip(universe.pos, universe.ref, universe.alt))
    with open(cfg.RESULTS / "spectra.pkl", "rb") as fh:
        spectra = pickle.load(fh)

    synonymous = oe_table(universe, spectra, "synonymous")
    print("synonymous calibration (expect observed/expected near 1)")
    print(synonymous.pivot(index="wire_class", columns="cohort",
                           values=["obs_over_exp", "p_value"])
          .reindex(list(cfg.WIRE_CLASSES)).round(3).to_string())

    missense = oe_table(universe, spectra, "missense")
    print("\nmissense selection")
    print(missense.pivot(index="wire_class", columns="cohort",
                         values=["obs_over_exp", "p_value"])
          .reindex(list(cfg.WIRE_CLASSES)).round(4).to_string())

    universe = annotate_covariates(universe)
    models = regressions(universe, spectra)
    print("\nloading-site rate ratio through the covariate ladder")
    print(models[models.wire_class == "loading_site"]
          .pivot(index="model", columns="cohort", values="rate_ratio")
          .reindex([m[0] for m in LADDER]).to_string())

    subunits = oe_by_subunit(universe, spectra)
    print("\nloading-site enrichment in PCAWG, by subunit")
    print(subunits.to_string(index=False))

    synonymous.to_csv(cfg.RESULTS / "oe_synonymous.csv", index=False)
    subunits.to_csv(cfg.RESULTS / "oe_by_subunit.csv", index=False)
    missense.to_csv(cfg.RESULTS / "oe_missense.csv", index=False)
    models.to_csv(cfg.RESULTS / "rate_ratios.csv", index=False)
    universe.drop(columns="key").to_parquet(
        cfg.RESULTS / "universe_annotated.parquet", index=False)
    print(f"\nwrote {cfg.RESULTS / 'oe_missense.csv'} and "
          f"{cfg.RESULTS / 'rate_ratios.csv'}")


if __name__ == "__main__":
    main()
```
