## Supplementary code for "Cancers relax selection on charge at the mitochondrial complex I quinone cavity": 06_heteroplasmy_disease.py

```
"""Heteroplasmy strata and the placement of mitochondrial disease variants."""
import pickle
import sys
from pathlib import Path

import numpy as np
import pandas as pd
from scipy import stats

sys.path.insert(0, str(Path(__file__).resolve().parents[1]))
import config as cfg

TUMOUR_FILES = {"TCGA": "data_mutations_tcga.txt.gz",
                "PCAWG": "data_mutations_pcawg.txt.gz",
                "MSK-IMPACT": "data_mutations_impact.txt.gz"}
TRUNCATING = ["Nonsense_Mutation", "Frame_Shift_Del", "Frame_Shift_Ins"]


def tumour_purity():
    table = pd.read_csv(cfg.MTDNA / "ext" /
                        "TCGA_mastercalls.abs_tables_JSedit.fixed.txt", sep="\t")
    return dict(zip(table["array"].str[:12],
                    pd.to_numeric(table.purity, errors="coerce")))


def heteroplasmy(universe, spectra):
    purity = tumour_purity()
    rows = []
    for cohort, filename in TUMOUR_FILES.items():
        calls = pd.read_csv(cfg.MTDNA / "original_data" / filename, sep="\t",
                            low_memory=False)
        nd_genes = sorted(universe.gene.unique())
        missense = calls[calls.Hugo_Symbol.isin(nd_genes)
                         & (calls.Variant_Classification == "Missense_Mutation")
                         & (calls.Variant_Type == "SNP")].copy()
        missense["key"] = list(zip(missense.Start_Position,
                                   missense.Reference_Allele,
                                   missense.Tumor_Seq_Allele2))
        lookup = dict(zip(universe.key, universe.wire_class))
        usable = dict(zip(universe.key, universe.identical))
        missense["wire_class"] = missense.key.map(lookup)
        missense["usable"] = missense.key.map(usable)
        missense["vaf"] = pd.to_numeric(missense.TumorVAF, errors="coerce")
        if cohort == "TCGA":
            patient = missense.Tumor_Sample_Barcode.str[:12]
            missense["vaf_corrected"] = np.minimum(
                missense.vaf / patient.map(purity), 1.0)
        missense = missense[missense.usable.fillna(False) & missense.vaf.notna()]

        subset = universe[(universe.kind == "missense") & universe.identical
                          & ~universe.polymorphism]
        weights = subset.channel.map(spectra[cohort]).fillna(spectra[cohort].min())
        columns = ["vaf"] + (["vaf_corrected"] if cohort == "TCGA" else [])
        for column in columns:
            d = missense[missense[column].notna()]
            for label, mask in [("< 0.5", d[column] < cfg.VAF_SPLIT),
                                (">= 0.5", d[column] >= cfg.VAF_SPLIT)]:
                stratum = d[mask]
                for wire_class in cfg.WIRE_CLASSES:
                    sel = (subset.wire_class == wire_class).to_numpy()
                    expected = len(stratum) * weights[sel].sum() / weights.sum()
                    observed = int((stratum.wire_class == wire_class).sum())
                    rows.append(dict(cohort=cohort, vaf=column, stratum=label,
                                     wire_class=wire_class, n_stratum=len(stratum),
                                     observed=observed,
                                     expected=round(expected, 1),
                                     obs_over_exp=round(observed / expected, 3)
                                     if expected else np.nan))
        truncating = pd.to_numeric(
            calls[calls.Hugo_Symbol.isin(nd_genes)
                  & calls.Variant_Classification.isin(TRUNCATING)].TumorVAF,
            errors="coerce").dropna()
        loading = missense[missense.wire_class == "loading_site"].vaf
        print(f"{cohort:11s} median VAF: loading site {loading.median():.3f}, "
              f"truncating {truncating.median():.3f}, "
              f"Mann-Whitney P = {stats.mannwhitneyu(loading, truncating).pvalue:.3f}")
    return pd.DataFrame(rows)


def disease(universe, spectra):
    columns = ["Start", "Ref", "Alt", "Mitomap_Status", "Mitomap_Phenotype",
               "ClinVar_Oct2017_CLINSIG"]
    mitimpact = pd.read_csv(cfg.MTDNA / "ext" / "MitImpact_db_2.9.txt", sep="\t",
                            low_memory=False, usecols=columns)
    keys = list(zip(mitimpact.Start, mitimpact.Ref, mitimpact.Alt))
    status = dict(zip(keys, mitimpact.Mitomap_Status))
    clinvar = dict(zip(keys, mitimpact.ClinVar_Oct2017_CLINSIG))

    subset = universe[(universe.kind == "missense") & universe.identical].copy()
    subset["mitomap"] = subset.key.map(status)
    subset["clinvar"] = subset.key.map(clinvar)
    subset["confirmed"] = subset.mitomap.eq("Cfrm")
    subset["reported"] = (
        subset.mitomap.fillna(".").str.contains("Cfrm|Reported|P.M.", regex=True)
        | subset.clinvar.fillna(".").str.contains("Pathogenic", na=False))
    subset["in_tumour"] = (subset.n_TCGA_missense + subset.n_PCAWG_missense
                           + subset["n_MSK-IMPACT_missense"]) > 0

    weights = subset.channel.map(spectra["germline"]).fillna(
        spectra["germline"].min())
    rows = []
    for label, column in [("MITOMAP confirmed", "confirmed"),
                          ("reported pathogenic", "reported")]:
        total = subset[column].sum()
        for wire_class in cfg.WIRE_CLASSES:
            mask = (subset.wire_class == wire_class).to_numpy()
            expected = total * weights[mask].sum() / weights.sum()
            observed = int(subset.loc[mask, column].sum())
            p = (stats.poisson.sf(observed - 1, expected) * 2 if observed > expected
                 else stats.poisson.cdf(observed, expected) * 2)
            rows.append(dict(variant_set=label, wire_class=wire_class,
                             observed=observed, expected=round(expected, 1),
                             obs_over_exp=round(observed / expected, 2)
                             if expected else np.nan, p_value=min(p, 1.0)))

    table = pd.crosstab(subset.reported, subset.in_tumour)
    odds, p = stats.fisher_exact(table.to_numpy())
    print(f"\ndisease and tumour variant overlap: odds ratio {odds:.2f}, P = {p:.3g}")
    overlap = pd.DataFrame([
        dict(group="reported pathogenic",
             observed_in_tumour=int(table.loc[True, True]),
             not_observed=int(table.loc[True, False])),
        dict(group="all other substitutions",
             observed_in_tumour=int(table.loc[False, True]),
             not_observed=int(table.loc[False, False]))])
    overlap["pct"] = (100 * overlap.observed_in_tumour
                      / (overlap.observed_in_tumour + overlap.not_observed)).round(2)
    overlap["odds_ratio"] = round(float(odds), 2)
    overlap["p_value"] = float(p)

    threshold = cfg.AXIS_A_THRESHOLD
    carriers = subset.key.map(dict(zip(universe.key, universe.polymorphism)))
    groups = [("all possible", subset),
              ("tumour-observed", subset[subset.in_tumour]),
              ("reported pathogenic", subset[subset.reported]),
              ("common germline", subset[carriers.fillna(False)])]
    scores = pd.read_parquet(cfg.RESULTS / "axis_scores.parquet")
    mapping = pd.read_csv(cfg.RESULTS / "mouse_human_map.csv")
    mapping = mapping[mapping.transferable]
    to_human = {(r.subunit, int(r.mouse_pos)): int(r.human_pos)
                for r in mapping.itertuples()}
    scores["human_pos"] = [to_human.get((s, p)) for s, p in
                           zip(scores.subunit, scores.pos)]
    axis_a = {(r.subunit, int(r.human_pos), r.mut_aa): r.axis_a_charge
              for r in scores[scores.human_pos.notna()].itertuples()}
    axis_rows = []
    for label, d in groups:
        values = np.array([axis_a.get((s, p, a), np.nan) for s, p, a in
                           zip(d.subunit, d.aa_pos, d.mut_aa)], float)
        values = values[~np.isnan(values)]
        axis_rows.append(dict(group=label, n=len(values),
                              pct_above_threshold=round(
                                  100 * float((values >= threshold).mean()), 2),
                              mean_axis_a=round(float(values.mean()), 5)))
    return pd.DataFrame(rows), pd.DataFrame(axis_rows), overlap


def main():
    universe = pd.read_parquet(cfg.RESULTS / "universe.parquet")
    universe["key"] = list(zip(universe.pos, universe.ref, universe.alt))
    with open(cfg.RESULTS / "spectra.pkl", "rb") as fh:
        spectra = pickle.load(fh)

    strata = heteroplasmy(universe, spectra)
    print("\nloading-site enrichment by heteroplasmy stratum")
    print(strata[strata.wire_class == "loading_site"]
          .pivot_table(index=["cohort", "vaf"], columns="stratum",
                       values="obs_over_exp").to_string())

    enrichment, axis_groups, overlap = disease(universe, spectra)
    print("\ndisease-variant enrichment")
    print(enrichment.to_string(index=False))
    print("\naxis A by variant group")
    print(axis_groups.to_string(index=False))
    print("\ndisease and tumour variant overlap")
    print(overlap.to_string(index=False))

    strata.to_csv(cfg.RESULTS / "heteroplasmy.csv", index=False)
    enrichment.to_csv(cfg.RESULTS / "disease_enrichment.csv", index=False)
    axis_groups.to_csv(cfg.RESULTS / "axis_a_by_group.csv", index=False)
    overlap.to_csv(cfg.RESULTS / "disease_overlap.csv", index=False)


if __name__ == "__main__":
    main()
```
