## Supplementary code for "Cancers relax selection on charge at the mitochondrial complex I quinone cavity": 07_nuclear.py

```
"""Nuclear mutational signatures in tumours carrying proton-wire variants.

If a loading-site substitution raised steady-state superoxide production, the
nuclear genome of the same tumour should carry more of the oxidative-damage
signatures. This script builds SBS96 profiles for carriers and non-carriers of
each wire class, fits the COSMIC v3.4 reference set, and tests the oxidative
readouts stratified by cancer type.
"""
import sys
from pathlib import Path

import numpy as np
import pandas as pd
from scipy import stats
from scipy.optimize import nnls

sys.path.insert(0, str(Path(__file__).resolve().parents[1]))
import config as cfg

PURINES = {"A": "T", "G": "C", "C": "G", "T": "A"}
PYRIMIDINES = ("C", "T")
OXIDATIVE = ("SBS18", "SBS36")
MAF_COLUMNS = ["Tumor_Sample_Barcode", "Chromosome", "Start_Position",
               "Reference_Allele", "Tumor_Seq_Allele2", "Variant_Type",
               "CONTEXT"]


def sbs96_channels():
    out = []
    for ref in PYRIMIDINES:
        for alt in "ACGT":
            if alt == ref:
                continue
            for left in "ACGT":
                for right in "ACGT":
                    out.append(f"{left}[{ref}>{alt}]{right}")
    return sorted(out)


CHANNELS = sbs96_channels()


def revcomp(seq):
    return "".join(PURINES[b] for b in reversed(seq))


def to_channel(ref, alt, context):
    """Pyrimidine-normalised SBS96 channel from a 3-base context."""
    if len(context) < 3:
        return None
    mid = len(context) // 2
    trinuc = context[mid - 1:mid + 2].upper()
    if len(trinuc) != 3 or any(b not in PURINES for b in trinuc):
        return None
    if ref not in PYRIMIDINES:
        ref, alt, trinuc = PURINES[ref], PURINES[alt], revcomp(trinuc)
    if ref not in PYRIMIDINES or alt not in "ACGT" or alt == ref:
        return None
    return f"{trinuc[0]}[{ref}>{alt}]{trinuc[2]}"


def nuclear_profiles(carriers):
    """One SBS96 count vector per patient, pooled over the TCGA studies."""
    profiles, cancer_type = {}, {}
    for path in sorted(cfg.NUCLEAR_MAFS.glob("*.maf.gz")):
        study = path.name.split("_")[0].upper()
        maf = pd.read_csv(path, sep="\t", comment="#", low_memory=False,
                          usecols=lambda c: c in MAF_COLUMNS)
        maf = maf[(maf.Variant_Type == "SNP") & maf.CONTEXT.notna()]
        if maf.empty:
            continue
        maf["patient"] = maf.Tumor_Sample_Barcode.str[:12]
        maf["channel"] = [to_channel(r, a, c) for r, a, c in
                          zip(maf.Reference_Allele, maf.Tumor_Seq_Allele2,
                              maf.CONTEXT)]
        maf = maf[maf.channel.notna()]
        table = (maf.pivot_table(index="patient", columns="channel",
                                 values="Start_Position", aggfunc="size")
                 .reindex(columns=CHANNELS).fillna(0))
        for patient, row in table.iterrows():
            profiles[patient] = row.to_numpy(float)
            cancer_type[patient] = study
        print(f"  {study:12s} {len(table):5d} patients, "
              f"{int(table.to_numpy().sum()):8d} SNVs")
    frame = pd.DataFrame(profiles, index=CHANNELS).T
    frame["cancer_type"] = pd.Series(cancer_type)
    for wire_class in ("loading_site", "cut_vertex", "cluster_member"):
        frame[wire_class] = frame.index.isin(carriers.get(wire_class, set()))
    return frame


def fit_cosmic(profile, reference, tolerance=0.01):
    """NNLS fit with backward elimination of signatures below `tolerance`."""
    total = profile.sum()
    if total == 0:
        return pd.Series(0.0, index=reference.columns)
    target = profile / total
    keep = list(reference.columns)
    while keep:
        weights, _ = nnls(reference[keep].to_numpy(), target)
        share = weights / weights.sum() if weights.sum() else weights
        weak = [name for name, s in zip(keep, share) if s < tolerance]
        if not weak or len(keep) - len(weak) < 1:
            break
        keep = [name for name in keep if name not in weak]
    out = pd.Series(0.0, index=reference.columns)
    out[keep] = share
    return out


def mantel_haenszel(frame, column, wire_class, quantile=0.75):
    """CMH test of a high signature share against carrier status by stratum."""
    threshold = frame[column].quantile(quantile)
    high = frame[column] >= threshold
    num = den = 0.0
    tables = []
    for study, stratum in frame.groupby("cancer_type"):
        h, c = high.loc[stratum.index], stratum[wire_class]
        a = int((h & c).sum()); b = int((~h & c).sum())
        cc = int((h & ~c).sum()); d = int((~h & ~c).sum())
        n = a + b + cc + d
        if n == 0 or (a + b) == 0 or (cc + d) == 0:
            continue
        num += a * d / n
        den += b * cc / n
        tables.append([[a, b], [cc, d]])
    if not tables or den == 0:
        return np.nan, np.nan, 0
    pooled = np.array(tables).sum(axis=0)
    return num / den, stats.fisher_exact(pooled)[1], len(tables)


def main():
    universe = pd.read_parquet(cfg.RESULTS / "universe.parquet")
    universe["key"] = list(zip(universe.pos, universe.ref, universe.alt))
    lookup = dict(zip(universe.key, universe.wire_class))
    usable = dict(zip(universe.key, universe.identical))

    calls = pd.read_csv(cfg.MTDNA / "original_data" / "data_mutations_tcga.txt.gz",
                        sep="\t", low_memory=False)
    nd = calls[calls.Hugo_Symbol.isin(sorted(universe.gene.unique()))
               & (calls.Variant_Classification == "Missense_Mutation")].copy()
    nd["key"] = list(zip(nd.Start_Position, nd.Reference_Allele,
                         nd.Tumor_Seq_Allele2))
    nd["wire_class"] = nd.key.map(lookup)
    nd = nd[nd.key.map(usable).fillna(False)]
    nd["patient"] = nd.Tumor_Sample_Barcode.str[:12]
    carriers = {name: set(group.patient)
                for name, group in nd.groupby("wire_class")}
    for name, patients in sorted(carriers.items()):
        print(f"carriers of a {name} variant: {len(patients)}")

    print("\nnuclear SBS96 profiles")
    frame = nuclear_profiles(carriers)
    print(f"total {len(frame):,} patients with both mitochondrial and nuclear "
          f"calls across {frame.cancer_type.nunique()} studies")

    reference = pd.read_csv(cfg.COSMIC, sep="\t", index_col=0)
    reference = reference.reindex(CHANNELS)
    reference = reference / reference.sum()

    shares = pd.DataFrame(
        [fit_cosmic(frame.loc[p, CHANNELS].to_numpy(float), reference)
         for p in frame.index], index=frame.index)
    frame = pd.concat([frame.drop(columns=CHANNELS), shares], axis=1)

    rows = []
    for wire_class in ("loading_site", "cut_vertex", "cluster_member"):
        if frame[wire_class].sum() < 5:
            continue
        for signature in OXIDATIVE:
            if signature not in frame:
                continue
            carrier = frame.loc[frame[wire_class], signature]
            other = frame.loc[~frame[wire_class], signature]
            odds, p_cmh, strata = mantel_haenszel(frame, signature, wire_class)
            rows.append(dict(
                wire_class=wire_class, signature=signature,
                n_carriers=int(frame[wire_class].sum()),
                median_carrier=round(float(carrier.median()), 4),
                median_other=round(float(other.median()), 4),
                mannwhitney_p=float(stats.mannwhitneyu(carrier, other).pvalue),
                cmh_odds_ratio=round(odds, 3) if np.isfinite(odds) else np.nan,
                cmh_p=p_cmh, n_strata=strata))
    result = pd.DataFrame(rows)
    print("\noxidative signature share by carrier status")
    print(result.to_string(index=False))

    burden = frame.groupby("loading_site").apply(
        lambda d: d[list(OXIDATIVE)].sum(axis=1).median())
    print(f"\ncombined {' + '.join(OXIDATIVE)} share, median: "
          f"non-carriers {burden.get(False, np.nan):.4f}, "
          f"carriers {burden.get(True, np.nan):.4f}")

    result.to_csv(cfg.RESULTS / "nuclear_signatures.csv", index=False)
    frame.to_csv(cfg.RESULTS / "nuclear_signature_shares.csv")
    print(f"\nwrote {cfg.RESULTS / 'nuclear_signatures.csv'}")


if __name__ == "__main__":
    main()
```
