## Supplementary code for "Cancers relax selection on charge at the mitochondrial complex I quinone cavity": 08_sensitivity.py

```
"""Sensitivity of the main result to the four discretionary choices.

1. the hydrogen-bond distance cutoff used to build the wire graph
2. the restriction to codons where mouse and human residues are identical
3. the HelixMTdb carrier count above which a somatic call is called a
   polymorphism
4. the definition of axis B, with and without its steric term

The wire is rebuilt from the structure at each cutoff rather than re-thresholded
from the primary graph, so the articulation points are those of the graph that
cutoff actually produces.
"""
import pickle
import sys
from pathlib import Path

import numpy as np
import pandas as pd

sys.path.insert(0, str(Path(__file__).resolve().parents[1]))
import config as cfg
from src import axes, variants, wire

POLYMORPHISM_SWEEP = (0, 5, 10, 50, 100)
IDENTITY_SETTINGS = (("identical codons only", True), ("all mapped codons", False))


def missense_oe(universe, spectra, require_identical=True,
                polymorphism_carriers=cfg.POLYMORPHISM_CARRIERS,
                carrier_counts=None):
    """Observed/expected per wire class, for one set of filter settings."""
    subset = universe[universe.kind == "missense"]
    if require_identical:
        subset = subset[subset.identical]
    out = {}
    for cohort in cfg.COHORTS:
        d = subset
        if cohort != "germline" and carrier_counts is not None:
            d = d[d.key.map(carrier_counts).fillna(0) <= polymorphism_carriers]
        for wire_class in cfg.WIRE_CLASSES:
            mask = (d.wire_class == wire_class).to_numpy()
            _, _, oe, _ = variants.observed_expected(
                d, spectra[cohort], f"n_{cohort}_missense", mask)
            out[(cohort, wire_class)] = oe
    return out


def reclassify(cutoff):
    """Rebuild the wire graph at `cutoff` and return the residue classes."""
    model = wire.load_model(cfg.MOUSE_CIF, cfg.MOUSE_CHAINS)
    centre, normal, _, proj = wire.membrane_frame(model)
    nodes = wire.collect_nodes(model, cfg.MOUSE_CHAINS)
    coords = np.asarray([n[3] for n in nodes])
    proj_normal = (coords - centre) @ normal
    graph, _, _, _ = wire.build_graph(nodes, cutoff)
    classes, comps, major, _ = wire.classify(nodes, graph, proj_normal)
    return classes, graph


def apply_classes(universe, mapping, classes):
    """Transfer a mouse classification onto the human universe in place."""
    human_class = {}
    for row in mapping.itertuples():
        human_class[(row.subunit, int(row.human_pos))] = classes.get(
            (row.subunit, int(row.mouse_pos)), "other")
    universe = universe.copy()
    universe["wire_class"] = [human_class.get((s, p), "other") for s, p in
                              zip(universe.subunit, universe.aa_pos)]
    return universe


def main():
    universe = pd.read_parquet(cfg.RESULTS / "universe.parquet")
    universe["key"] = list(zip(universe.pos, universe.ref, universe.alt))
    with open(cfg.RESULTS / "spectra.pkl", "rb") as fh:
        spectra = pickle.load(fh)

    mapping = pd.read_csv(cfg.RESULTS / "mouse_human_map.csv")
    mapping = mapping[mapping.transferable & mapping.human_pos.notna()]

    helix = pd.read_csv(cfg.MTDNA / "original_data" / "data_mutations_helix.txt.gz",
                        sep="\t", low_memory=False,
                        usecols=["Start_Position", "Reference_Allele",
                                 "Tumor_Seq_Allele2", "TOTAL_COUNT"])
    helix["TOTAL_COUNT"] = pd.to_numeric(helix.TOTAL_COUNT,
                                         errors="coerce").fillna(0)
    carrier_counts = (helix.groupby(["Start_Position", "Reference_Allele",
                                     "Tumor_Seq_Allele2"]).TOTAL_COUNT.max()
                      .to_dict())

    # --- 1. hydrogen-bond cutoff -----------------------------------------
    rows = []
    for cutoff in cfg.HBOND_SWEEP:
        classes, graph = reclassify(cutoff)
        counts = pd.Series(list(classes.values())).value_counts()
        reclassified = apply_classes(universe, mapping, classes)
        oe = missense_oe(reclassified, spectra, carrier_counts=carrier_counts)
        for (cohort, wire_class), value in oe.items():
            rows.append(dict(cutoff=cutoff, n_edges=graph.number_of_edges(),
                             n_residues=int(counts.get(wire_class, 0)),
                             cohort=cohort, wire_class=wire_class,
                             obs_over_exp=round(value, 3)
                             if np.isfinite(value) else np.nan))
        print(f"cutoff {cutoff} A: {graph.number_of_edges():,} edges, "
              + ", ".join(f"{k} {int(counts.get(k, 0))}" for k in
                          ("loading_site", "cut_vertex", "cluster_member")))
    cutoff_table = pd.DataFrame(rows)
    print("\nloading-site observed/expected by cutoff")
    print(cutoff_table[cutoff_table.wire_class == "loading_site"]
          .pivot(index="cutoff", columns="cohort", values="obs_over_exp")
          .to_string())
    print("\ncut-vertex observed/expected by cutoff")
    print(cutoff_table[cutoff_table.wire_class == "cut_vertex"]
          .pivot(index="cutoff", columns="cohort", values="obs_over_exp")
          .to_string())

    # --- 2. the mouse-human identity filter ------------------------------
    rows = []
    for label, require in IDENTITY_SETTINGS:
        oe = missense_oe(universe, spectra, require_identical=require,
                         carrier_counts=carrier_counts)
        n = int(((universe.kind == "missense")
                 & (universe.identical | ~require)).sum())
        for (cohort, wire_class), value in oe.items():
            rows.append(dict(setting=label, n_substitutions=n, cohort=cohort,
                             wire_class=wire_class,
                             obs_over_exp=round(value, 3)
                             if np.isfinite(value) else np.nan))
    identity_table = pd.DataFrame(rows)
    print("\nloading-site observed/expected with and without the identity filter")
    print(identity_table[identity_table.wire_class == "loading_site"]
          .pivot(index="setting", columns="cohort", values="obs_over_exp")
          .to_string())

    # --- 3. the polymorphism threshold -----------------------------------
    rows = []
    for threshold in POLYMORPHISM_SWEEP:
        oe = missense_oe(universe, spectra, polymorphism_carriers=threshold,
                         carrier_counts=carrier_counts)
        for (cohort, wire_class), value in oe.items():
            if cohort == "germline":
                continue
            rows.append(dict(carrier_threshold=threshold, cohort=cohort,
                             wire_class=wire_class,
                             obs_over_exp=round(value, 3)
                             if np.isfinite(value) else np.nan))
    polymorphism_table = pd.DataFrame(rows)
    print("\nloading-site observed/expected by polymorphism threshold")
    print(polymorphism_table[polymorphism_table.wire_class == "loading_site"]
          .pivot(index="carrier_threshold", columns="cohort",
                 values="obs_over_exp").to_string())

    # --- 4. the definition of axis B -------------------------------------
    scores = pd.read_parquet(cfg.RESULTS / "axis_scores.parquet")
    rows = []
    for label, column in [("with steric term", "axis_b_connectivity"),
                          ("topology only", "axis_b_topology_only")]:
        stats_ = axes.separability(scores, column)
        above = scores[column] >= cfg.AXIS_B_THRESHOLD
        rows.append(dict(definition=label,
                         n_above_threshold=int(above.sum()),
                         residues_above=int(scores[above]
                                            .groupby(["subunit", "pos"]).ngroups),
                         pearson_r=round(stats_["pearson_r_all"], 3),
                         n_connectivity_only=stats_["n_connectivity_only"],
                         n_charge_only=stats_["n_charge_only"]))
    axis_table = pd.DataFrame(rows)
    print("\naxis B definition")
    print(axis_table.to_string(index=False))

    cutoff_table.to_csv(cfg.RESULTS / "sensitivity_cutoff.csv", index=False)
    identity_table.to_csv(cfg.RESULTS / "sensitivity_identity.csv", index=False)
    polymorphism_table.to_csv(cfg.RESULTS / "sensitivity_polymorphism.csv",
                              index=False)
    axis_table.to_csv(cfg.RESULTS / "sensitivity_axis_b.csv", index=False)
    print(f"\nwrote four sensitivity tables to {cfg.RESULTS}")


if __name__ == "__main__":
    main()
```
