## Supplementary code for "Cancers relax selection on charge at the mitochondrial complex I quinone cavity": 09_figures.py

```
"""Build the figures that scripts 01-08 support.

Each figure is written as a vector PDF and a 600 dpi PNG. Nothing here
recomputes a statistic; the panels read the tables produced by scripts 01-08,
so a figure can never disagree with the numbers in the text. Figures 2 and 4
draw on the interaction and quinone analyses as well, and are built by
26_main_figures.py once those have run.
"""
import pickle
import sys
from pathlib import Path

import numpy as np
import pandas as pd
import matplotlib.pyplot as plt

sys.path.insert(0, str(Path(__file__).resolve().parents[1]))
import config as cfg
from src import plotting
from src.plotting import CLASS_COLOURS, CLASS_LABELS, COHORT_MARKERS

CLASSES = ("loading_site", "cut_vertex", "cluster_member", "other")
SCORED = ("loading_site", "cut_vertex", "cluster_member")


def read(name, **kwargs):
    return pd.read_csv(cfg.RESULTS / name, **kwargs)


# --- Figure 1: the proton wire -------------------------------------------

def figure1(wire_built, clusters, residues, scores):
    fig = plt.figure(figsize=(11.0, 8.4))
    grid = fig.add_gridspec(2, 3, hspace=0.42, wspace=0.36)

    nodes = wire_built["nodes"]
    coords = np.asarray([n[3] for n in nodes])
    names = [n[1] for n in nodes]
    major = {i for k in wire_built["major"] for i in wire_built["comps"][k]}
    normal = np.asarray(wire_built["normal"])
    centre = np.asarray(wire_built["centre"])
    along = (coords - centre) @ normal
    lateral = (coords - centre) @ np.asarray(wire_built["in_plane"])

    ax = fig.add_subplot(grid[0, :2])
    rest = [i for i in range(len(nodes)) if i not in major]
    ax.scatter(lateral[rest], along[rest], s=4, c="#d5d8dc",
               label="Outside a major cluster", linewidths=0)
    water = [i for i in sorted(major) if names[i] == "HOH"]
    side = [i for i in sorted(major) if names[i] != "HOH"]
    ax.scatter(lateral[water], along[water], s=7, c="#5dade2",
               label="Cluster water", linewidths=0)
    ax.scatter(lateral[side], along[side], s=22, c=CLASS_COLOURS["loading_site"],
               label="Cluster side chain", linewidths=0)
    ax.axhline(0, color="black", lw=0.6, ls=":")
    ax.set_xlabel("Position along the membrane plane (Å)")
    ax.set_ylabel("Position along the membrane normal (Å)\nmatrix side positive")
    ax.legend(loc="upper left", ncol=3, handletextpad=0.3, columnspacing=1.0)
    plotting.panel_label(ax, "a", dx=-0.09)

    ax = fig.add_subplot(grid[0, 2])
    sizes = np.sort(clusters.n_nodes.to_numpy())[::-1]
    ax.bar(np.arange(1, len(sizes) + 1), sizes, color="#5dade2", width=0.9)
    ax.set_yscale("log")
    ax.set_xlabel("Hydrated cluster, rank by size")
    ax.set_ylabel("Nodes in cluster")
    ax.axhline(cfg.MAJOR_MIN_NODES, color="black", lw=0.8, ls="--")
    ax.text(len(sizes) * 0.45, cfg.MAJOR_MIN_NODES * 1.25,
            f"major cluster\nthreshold ({cfg.MAJOR_MIN_NODES})", fontsize=9)
    plotting.panel_label(ax, "b")

    ax = fig.add_subplot(grid[1, 0])
    counts = (residues.wire_class.value_counts()
              .reindex(SCORED).fillna(0).astype(int))
    ax.bar(range(len(counts)), counts.to_numpy(),
           color=[CLASS_COLOURS[c] for c in counts.index], width=0.65)
    for x, value in enumerate(counts.to_numpy()):
        ax.text(x, value + counts.max() * 0.02, str(value), ha="center",
                fontsize=10)
    ax.set_xticks(range(len(counts)))
    ax.set_xticklabels(["Loading\nsite", "Cut\nvertex", "Cluster\nmember"])
    ax.set_ylabel("ND residues")
    ax.set_ylim(0, counts.max() * 1.18)
    plotting.panel_label(ax, "c")

    ax = fig.add_subplot(grid[1, 1])
    by_subunit = (residues[residues.wire_class.isin(SCORED)]
                  .pivot_table(index="subunit", columns="wire_class",
                               values="pos", aggfunc="size")
                  .reindex(index=list(cfg.SUBUNITS), columns=list(SCORED))
                  .fillna(0))
    bottom = np.zeros(len(by_subunit))
    for wire_class in SCORED:
        values = by_subunit[wire_class].to_numpy()
        ax.bar(range(len(by_subunit)), values, bottom=bottom, width=0.7,
               color=CLASS_COLOURS[wire_class], label=CLASS_LABELS[wire_class])
        bottom += values
    ax.set_xticks(range(len(by_subunit)))
    ax.set_xticklabels(by_subunit.index, rotation=45, ha="right")
    ax.set_ylabel("Wire residues")
    ax.legend(loc="upper left", fontsize=9)
    plotting.panel_label(ax, "d")

    ax = fig.add_subplot(grid[1, 2])
    sample = scores.sample(min(len(scores), 40000), random_state=cfg.RANDOM_SEED)
    ax.scatter(sample.axis_a_charge, sample.axis_b_connectivity, s=5,
               c="#aab7b8", linewidths=0, alpha=0.5)
    both = sample[(sample.axis_a_charge >= cfg.AXIS_A_THRESHOLD)
                  & (sample.axis_b_connectivity >= cfg.AXIS_B_THRESHOLD)]
    ax.scatter(both.axis_a_charge, both.axis_b_connectivity, s=10,
               c=CLASS_COLOURS["loading_site"], linewidths=0)
    ax.axvline(cfg.AXIS_A_THRESHOLD, color="black", lw=0.8, ls="--")
    ax.axhline(cfg.AXIS_B_THRESHOLD, color="black", lw=0.8, ls="--")
    r = np.corrcoef(scores.axis_a_charge, scores.axis_b_connectivity)[0, 1]
    ax.set_xlabel("Axis A, electrostatic shift (eV)")
    ax.set_ylabel("Axis B, connectivity loss (a.u.)")
    ax.set_title(f"Pearson $r$ = {r:.2f}", fontsize=11)
    plotting.panel_label(ax, "e")

    return plotting.save(fig, cfg.FIGURES, "Figure1")


# --- Figure 2: selection --------------------------------------------------

def _oe_panel(ax, table, title, log=True):
    offsets = np.linspace(-0.26, 0.26, len(cfg.COHORTS))
    for offset, cohort in zip(offsets, cfg.COHORTS):
        d = table[table.cohort == cohort].set_index("wire_class").reindex(CLASSES)
        x = np.arange(len(CLASSES)) + offset
        ax.errorbar(x, d.obs_over_exp,
                    yerr=[d.obs_over_exp - d.ci_low, d.ci_high - d.obs_over_exp],
                    fmt=COHORT_MARKERS[cohort], ms=6, lw=1.2, capsize=2.5,
                    color="black", mfc=CLASS_COLOURS["loading_site"]
                    if cohort == "germline" else "white", label=cohort)
    ax.axhline(1.0, color="black", lw=0.8, ls=":")
    ax.set_xticks(np.arange(len(CLASSES)))
    ax.set_xticklabels([CLASS_LABELS[c].replace(" ", "\n") for c in CLASSES])
    ax.set_ylabel("Observed / expected")
    if log:
        ax.set_yscale("log")
        ax.set_yticks([0.25, 0.5, 1.0, 2.0])
        ax.set_yticklabels(["0.25", "0.5", "1", "2"])
    ax.set_title(title, fontsize=11)


# --- Figure 3: heteroplasmy ----------------------------------------------

def figure3(heteroplasmy, missense):
    fig, axes_ = plt.subplots(1, 3, figsize=(13.0, 4.4))

    ax = axes_[0]
    d = heteroplasmy[(heteroplasmy.wire_class == "loading_site")
                     & (heteroplasmy.vaf == "vaf")]
    cohorts = [c for c in cfg.COHORTS if c != "germline"]
    width = 0.36
    for i, stratum in enumerate(("< 0.5", ">= 0.5")):
        s = d[d.stratum == stratum].set_index("cohort").reindex(cohorts)
        ax.bar(np.arange(len(cohorts)) + (i - 0.5) * width, s.obs_over_exp,
               width=width, color=["#aab7b8", CLASS_COLOURS["loading_site"]][i],
               label=f"Heteroplasmy {stratum}")
    ax.axhline(1.0, color="black", lw=0.8, ls=":")
    ax.set_xticks(np.arange(len(cohorts)))
    ax.set_xticklabels(cohorts, rotation=20, ha="right")
    ax.set_ylabel("Observed / expected, loading sites")
    ax.legend(fontsize=9)
    plotting.panel_label(ax, "a")

    ax = axes_[1]
    d = heteroplasmy[(heteroplasmy.cohort == "TCGA")
                     & (heteroplasmy.wire_class == "loading_site")]
    labels = {"vaf": "Uncorrected", "vaf_corrected": "Purity-corrected"}
    for i, column in enumerate(("vaf", "vaf_corrected")):
        s = d[d.vaf == column].set_index("stratum").reindex(["< 0.5", ">= 0.5"])
        ax.bar(np.arange(2) + (i - 0.5) * width, s.obs_over_exp, width=width,
               color=["#aab7b8", CLASS_COLOURS["loading_site"]][i],
               label=labels[column])
    ax.axhline(1.0, color="black", lw=0.8, ls=":")
    ax.set_xticks(np.arange(2))
    ax.set_xticklabels(["Heteroplasmy < 0.5", "Heteroplasmy ≥ 0.5"])
    ax.set_ylabel("Observed / expected, TCGA")
    ax.legend(fontsize=9)
    plotting.panel_label(ax, "b")

    ax = axes_[2]
    for offset, wire_class in zip((-0.2, 0.0, 0.2), SCORED):
        d = (missense[missense.wire_class == wire_class]
             .set_index("cohort").reindex(list(cfg.COHORTS)))
        ax.errorbar(np.arange(len(cfg.COHORTS)) + offset, d.obs_over_exp,
                    yerr=[d.obs_over_exp - d.ci_low, d.ci_high - d.obs_over_exp],
                    fmt="o", ms=6, lw=1.2, capsize=2.5, color="black",
                    mfc=CLASS_COLOURS[wire_class],
                    label=CLASS_LABELS[wire_class])
    ax.axhline(1.0, color="black", lw=0.8, ls=":")
    ax.set_xticks(np.arange(len(cfg.COHORTS)))
    ax.set_xticklabels(list(cfg.COHORTS), rotation=20, ha="right")
    ax.set_yscale("log")
    ax.set_ylabel("Observed / expected")
    ax.legend(fontsize=9)
    plotting.panel_label(ax, "c")

    fig.tight_layout()
    return plotting.save(fig, cfg.FIGURES, "Figure3")


# --- Supplementary figures ------------------------------------------------

def supplementary1(wire_built):
    """The membrane frame recovered from the modelled lipids."""
    fig, axes_ = plt.subplots(1, 2, figsize=(9.5, 4.2))
    lipid = np.asarray(wire_built["lipid_xyz"])
    centre = np.asarray(wire_built["centre"])
    normal = np.asarray(wire_built["normal"])
    lateral = np.asarray(wire_built["in_plane"])
    along = (lipid - centre) @ normal
    across = (lipid - centre) @ lateral

    axes_[0].scatter(across, along, s=3, c="#5dade2", linewidths=0)
    axes_[0].axhline(0, color="black", lw=0.8, ls="--")
    axes_[0].set_xlabel("Membrane plane (Å)")
    axes_[0].set_ylabel("Membrane normal (Å)")
    axes_[0].set_title("Modelled lipid atoms", fontsize=11)
    plotting.panel_label(axes_[0], "a")

    axes_[1].hist(along, bins=60, color="#5dade2")
    axes_[1].set_xlabel("Position along the membrane normal (Å)")
    axes_[1].set_ylabel("Lipid atoms")
    axes_[1].set_title("Bilayer thickness", fontsize=11)
    plotting.panel_label(axes_[1], "b")

    fig.tight_layout()
    return plotting.save(fig, cfg.FIGURES, "SuppFig1")


def supplementary2(mapping, clashes, strategies):
    """Why the wire is built in the mouse structure."""
    fig, axes_ = plt.subplots(1, 3, figsize=(13.0, 4.2))

    ax = axes_[0]
    identity = (mapping[mapping.transferable].groupby("subunit").identical
                .mean().reindex(list(cfg.SUBUNITS)) * 100)
    ax.bar(range(len(identity)), identity.to_numpy(), color="#aab7b8", width=0.7)
    wire_identity = (mapping[mapping.transferable
                             & (mapping.wire_class != "other")]
                     .groupby("subunit").identical.mean()
                     .reindex(list(cfg.SUBUNITS)) * 100)
    ax.plot(range(len(identity)), wire_identity.to_numpy(), "o-",
            color=CLASS_COLOURS["loading_site"], ms=6, label="Wire residues")
    ax.set_xticks(range(len(identity)))
    ax.set_xticklabels(identity.index, rotation=45, ha="right")
    ax.set_ylabel("Mouse–human identity (%)")
    ax.set_ylim(0, 105)
    ax.legend(fontsize=9)
    plotting.panel_label(ax, "a")

    ax = axes_[1]
    ax.bar(range(len(strategies)), strategies.n_edges, color="#5dade2", width=0.6)
    ax.set_yscale("log")
    ax.set_xticks(range(len(strategies)))
    ax.set_xticklabels([s.replace(" ", "\n") for s in strategies.strategy],
                       fontsize=9)
    ax.set_ylabel("Hydrogen-bond edges")
    ax.set_title("Graph size by water source", fontsize=11)
    plotting.panel_label(ax, "b")

    ax = axes_[2]
    ax.hist(clashes.min_distance, bins=40, color="#aab7b8")
    ax.axvline(cfg.HBOND_MIN, color="black", lw=1.0, ls="--")
    ax.set_xlabel("Closest protein atom after transfer (Å)")
    ax.set_ylabel("Transferred waters")
    ax.set_title(f"{int((clashes.min_distance < cfg.HBOND_MIN).sum()):,} of "
                 f"{len(clashes):,} clash", fontsize=11)
    plotting.panel_label(ax, "c")

    fig.tight_layout()
    return plotting.save(fig, cfg.FIGURES, "SuppFig2")


def supplementary3(synonymous, universe):
    """The null model and its calibration."""
    fig, axes_ = plt.subplots(1, 2, figsize=(10.5, 4.2))

    ax = axes_[0]
    counts = universe[universe.kind == "synonymous"].codon_pos.value_counts()
    counts = counts.reindex([1, 2, 3]).fillna(0)
    ax.bar(range(3), counts.to_numpy(), color="#5dade2", width=0.6)
    ax.set_xticks(range(3))
    ax.set_xticklabels(["1st", "2nd", "3rd"])
    ax.set_xlabel("Codon position")
    ax.set_ylabel("Synonymous substitutions")
    ax.set_title("Universe built from first principles", fontsize=11)
    plotting.panel_label(ax, "a")

    _oe_panel(axes_[1], synonymous, "Synonymous observed / expected", log=False)
    axes_[1].set_ylim(0.5, 1.6)
    axes_[1].legend(fontsize=9, ncol=2)
    plotting.panel_label(axes_[1], "b")

    fig.tight_layout()
    return plotting.save(fig, cfg.FIGURES, "SuppFig3")


def supplementary4(mapping):
    """Coverage of the mouse to human mapping, subunit by subunit."""
    fig, axes_ = plt.subplots(3, 1, figsize=(10.0, 9.0))

    ax = axes_[0]
    coverage = mapping.groupby("subunit").transferable.agg(["sum", "size"])
    coverage = coverage.reindex(list(cfg.SUBUNITS))
    ax.bar(range(len(coverage)), coverage["size"], color="#d5d8dc", width=0.7,
           label="Modelled in 8OM1")
    ax.bar(range(len(coverage)), coverage["sum"], color="#5dade2", width=0.7,
           label="Mapped to a human codon")
    ax.set_xticks(range(len(coverage)))
    ax.set_xticklabels(coverage.index, rotation=45, ha="right")
    ax.set_ylabel("Residues")
    ax.legend(fontsize=9)
    plotting.panel_label(ax, "a", dx=-0.08)

    ax = axes_[1]
    wire_rows = mapping[mapping.wire_class != "other"]
    table = (wire_rows.pivot_table(index="subunit", columns="wire_class",
                                   values="identical", aggfunc="mean")
             .reindex(index=list(cfg.SUBUNITS), columns=list(SCORED)) * 100)
    for i, wire_class in enumerate(SCORED):
        ax.bar(np.arange(len(table)) + (i - 1) * 0.27, table[wire_class],
               width=0.27, color=CLASS_COLOURS[wire_class],
               label=CLASS_LABELS[wire_class])
    ax.set_xticks(np.arange(len(table)))
    ax.set_xticklabels(table.index, rotation=45, ha="right")
    ax.set_ylabel("Identical in both species (%)")
    ax.legend(fontsize=9)
    plotting.panel_label(ax, "b", dx=-0.08)

    ax = axes_[2]
    for wire_class in SCORED:
        d = wire_rows[(wire_rows.wire_class == wire_class)
                      & wire_rows.human_pos.notna()]
        ax.scatter(d.mouse_pos, d.human_pos, s=14, linewidths=0,
                   color=CLASS_COLOURS[wire_class],
                   label=CLASS_LABELS[wire_class])
    limit = float(np.nanmax(wire_rows.mouse_pos)) * 1.05
    ax.plot([0, limit], [0, limit], color="black", lw=0.8, ls=":")
    ax.set_xlabel("Mouse residue number")
    ax.set_ylabel("Human residue number")
    ax.legend(fontsize=9)
    plotting.panel_label(ax, "c", dx=-0.08)

    fig.tight_layout()
    return plotting.save(fig, cfg.FIGURES, "SuppFig4")


def supplementary5(cutoff, identity, polymorphism, axis_b):
    """The four sensitivity analyses."""
    fig, axes_ = plt.subplots(2, 2, figsize=(11.0, 8.0))

    ax = axes_[0, 0]
    for wire_class in SCORED:
        d = (cutoff[(cutoff.wire_class == wire_class)
                    & (cutoff.cohort == "TCGA")]
             .sort_values("cutoff"))
        ax.plot(d.cutoff, d.obs_over_exp, "o-", ms=6,
                color=CLASS_COLOURS[wire_class], label=CLASS_LABELS[wire_class])
    ax.axhline(1.0, color="black", lw=0.8, ls=":")
    ax.set_xlabel("Hydrogen-bond cutoff (Å)")
    ax.set_ylabel("Observed / expected, TCGA")
    ax.legend(fontsize=9)
    plotting.panel_label(ax, "a")

    ax = axes_[0, 1]
    settings = list(identity.setting.unique())
    for i, setting in enumerate(settings):
        d = (identity[(identity.setting == setting)
                      & (identity.wire_class == "loading_site")]
             .set_index("cohort").reindex(list(cfg.COHORTS)))
        ax.bar(np.arange(len(cfg.COHORTS)) + (i - 0.5) * 0.38, d.obs_over_exp,
               width=0.38, color=["#aab7b8", CLASS_COLOURS["loading_site"]][i],
               label=setting)
    ax.axhline(1.0, color="black", lw=0.8, ls=":")
    ax.set_xticks(np.arange(len(cfg.COHORTS)))
    ax.set_xticklabels(list(cfg.COHORTS), rotation=20, ha="right")
    ax.set_ylabel("Observed / expected, loading sites")
    ax.legend(fontsize=9)
    plotting.panel_label(ax, "b")

    ax = axes_[1, 0]
    for cohort in [c for c in cfg.COHORTS if c != "germline"]:
        d = (polymorphism[(polymorphism.cohort == cohort)
                          & (polymorphism.wire_class == "loading_site")]
             .sort_values("carrier_threshold"))
        ax.plot(d.carrier_threshold, d.obs_over_exp,
                COHORT_MARKERS[cohort] + "-", ms=6, color="black",
                mfc=CLASS_COLOURS["loading_site"], label=cohort)
    ax.axhline(1.0, color="black", lw=0.8, ls=":")
    ax.set_xscale("symlog", linthresh=5)
    ax.set_xlabel("HelixMTdb carrier count treated as polymorphic")
    ax.set_ylabel("Observed / expected, loading sites")
    ax.legend(fontsize=9)
    plotting.panel_label(ax, "c")

    ax = axes_[1, 1]
    x = np.arange(len(axis_b))
    ax.bar(x - 0.19, axis_b.n_connectivity_only, width=0.38, color="#aab7b8",
           label="Connectivity only")
    ax.bar(x + 0.19, axis_b.n_charge_only, width=0.38,
           color=CLASS_COLOURS["loading_site"], label="Charge only")
    for xi, r in zip(x, axis_b.pearson_r):
        ax.text(xi, max(axis_b.n_connectivity_only.max(),
                        axis_b.n_charge_only.max()) * 1.04,
                f"$r$ = {r:.2f}", ha="center", fontsize=9)
    ax.set_xticks(x)
    ax.set_xticklabels([d.replace(" ", "\n") for d in axis_b.definition])
    ax.set_ylabel("Substitutions scoring on one axis")
    ax.legend(fontsize=9)
    plotting.panel_label(ax, "d")

    fig.tight_layout()
    return plotting.save(fig, cfg.FIGURES, "SuppFig5")


def supplementary6(shares, nuclear):
    """Nuclear signature shares behind the oxidative comparison."""
    fig, axes_ = plt.subplots(1, 2, figsize=(11.0, 4.4))
    signatures = [s for s in nuclear.signature.unique() if s in shares]

    ax = axes_[0]
    data = [shares.loc[shares.loading_site, s].dropna() for s in signatures]
    data += [shares.loc[~shares.loading_site, s].dropna() for s in signatures]
    box = ax.boxplot(data, patch_artist=True, widths=0.6, showfliers=False)
    for patch, colour in zip(box["boxes"],
                             [CLASS_COLOURS["loading_site"]] * len(signatures)
                             + ["#aab7b8"] * len(signatures)):
        patch.set_facecolor(colour)
    for element in ("medians", "whiskers", "caps"):
        for item in box[element]:
            item.set_color("black")
    ax.set_xticklabels([f"{s}\ncarrier" for s in signatures]
                       + [f"{s}\nnon-carrier" for s in signatures], fontsize=9)
    ax.set_ylabel("Signature share of the nuclear profile")
    plotting.panel_label(ax, "a")

    ax = axes_[1]
    by_study = (shares.groupby("cancer_type")
                .agg(n=("loading_site", "size"),
                     carriers=("loading_site", "sum")))
    by_study = by_study[by_study.carriers > 0].sort_values("carriers")
    ax.barh(range(len(by_study)), by_study.carriers,
            color=CLASS_COLOURS["loading_site"], height=0.7)
    ax.set_yticks(range(len(by_study)))
    ax.set_yticklabels([f"{s} ({n})" for s, n in zip(by_study.index, by_study.n)],
                       fontsize=8)
    ax.set_xlabel("Loading-site carriers with nuclear calls")
    ax.spines["left"].set_visible(True)
    plotting.panel_label(ax, "b", dx=-0.34)

    fig.tight_layout()
    return plotting.save(fig, cfg.FIGURES, "SuppFig6")


def main():
    plotting.use_style()
    with open(cfg.RESULTS / "wire.pkl", "rb") as fh:
        built = pickle.load(fh)

    residues = read("residue_classes.csv")
    clusters = read("hydrated_clusters.csv")
    clashes = read("transferred_water_clashes.csv")
    strategies = read("graph_strategies.csv")
    mapping = read("mouse_human_map.csv")
    scores = pd.read_parquet(cfg.RESULTS / "axis_scores.parquet")
    universe = pd.read_parquet(cfg.RESULTS / "universe.parquet")

    written = [
        figure1(built, clusters, residues, scores),
        figure3(read("heteroplasmy.csv"), read("oe_missense.csv")),
        supplementary1(built),
        supplementary2(mapping, clashes, strategies),
        supplementary3(read("oe_synonymous.csv"), universe),
        supplementary4(mapping),
        supplementary5(read("sensitivity_cutoff.csv"),
                       read("sensitivity_identity.csv"),
                       read("sensitivity_polymorphism.csv"),
                       read("sensitivity_axis_b.csv")),
        supplementary6(read("nuclear_signature_shares.csv", index_col=0),
                       read("nuclear_signatures.csv")),
    ]
    for path in written:
        print(f"wrote {path} and {path.with_suffix('.png')}")


if __name__ == "__main__":
    main()
```
