## Supplementary code for "Cancers relax selection on charge at the mitochondrial complex I quinone cavity": 10_recurrence.py

```
"""Recurrence and clonality: tolerance or positive selection?

Cross-sectional allele fractions cannot separate the two on their own, but two
properties of the observed calls bear on the distinction.

Recurrence. Under tolerance, somatic substitutions at loading sites accumulate
wherever mutation puts them, so the calls should be spread over sites in
proportion to mutational opportunity. Under positive selection, particular
substitutions should recur across patients more often than that. The null is
the same spectrum-weighted multinomial used for the selection analysis, so the
test asks about the distribution of calls over sites, holding their total
number fixed.

Clonality. A variant under positive selection within a tumour should reach
fixation more often than a merely tolerated one. The comparison is against
other missense substitutions in the same genes and against truncating
variants, which are the class most plausibly under selection in complex I.
"""
import pickle
import sys
from pathlib import Path

import numpy as np
import pandas as pd
from scipy import stats

sys.path.insert(0, str(Path(__file__).resolve().parents[1]))
import config as cfg

TUMOUR_FILES = {"TCGA": "data_mutations_tcga.txt.gz",
                "PCAWG": "data_mutations_pcawg.txt.gz",
                "MSK-IMPACT": "data_mutations_impact.txt.gz"}
TRUNCATING = ("Nonsense_Mutation", "Frame_Shift_Del", "Frame_Shift_Ins")
CLONAL_VAF = 0.9
MIN_PATIENTS = 4
AXIS_CLASSES = ("loading_site", "cut_vertex")
AA3TO1 = {"Ala": "A", "Arg": "R", "Asn": "N", "Asp": "D", "Cys": "C",
          "Gln": "Q", "Glu": "E", "Gly": "G", "His": "H", "Ile": "I",
          "Leu": "L", "Lys": "K", "Met": "M", "Phe": "F", "Pro": "P",
          "Ser": "S", "Thr": "T", "Trp": "W", "Tyr": "Y", "Val": "V"}


def load_calls(universe):
    lookup = dict(zip(universe.key, universe.wire_class))
    usable = dict(zip(universe.key, universe.identical))
    polymorphic = dict(zip(universe.key, universe.polymorphism))
    genes = sorted(universe.gene.unique())
    out = {}
    for cohort, filename in TUMOUR_FILES.items():
        calls = pd.read_csv(cfg.MTDNA / "original_data" / filename, sep="\t",
                            low_memory=False)
        calls = calls[calls.Hugo_Symbol.isin(genes)].copy()
        calls["key"] = list(zip(calls.Start_Position, calls.Reference_Allele,
                                calls.Tumor_Seq_Allele2))
        calls["patient"] = calls.Tumor_Sample_Barcode.str[:12]
        calls["vaf"] = pd.to_numeric(calls.TumorVAF, errors="coerce")
        calls["wire_class"] = calls.key.map(lookup)
        calls["usable"] = calls.key.map(usable).fillna(False)
        calls["polymorphism"] = calls.key.map(polymorphic).fillna(False)
        out[cohort] = calls
    return out


def recurrence(universe, spectra, calls, rng):
    """Are the calls more concentrated on single sites than the null allows?"""
    rows = []
    for cohort, frame in calls.items():
        eligible = universe[(universe.kind == "missense") & universe.identical
                            & ~universe.polymorphism].reset_index(drop=True)
        weights = eligible.channel.map(spectra[cohort])
        weights = np.array((weights.fillna(weights.min())
                            / weights.sum()).to_numpy(), dtype=float)
        # numpy requires the probabilities to sum to at most one; scale just
        # below one so floating-point accumulation cannot exceed it.
        weights = weights / (weights.sum() * (1.0 + 1e-9))
        klass = eligible.wire_class.to_numpy()
        masks = {c: klass == c for c in cfg.WIRE_CLASSES}

        observed = frame[(frame.Variant_Classification == "Missense_Mutation")
                         & frame.usable & ~frame.polymorphism]
        observed = observed.drop_duplicates(["key", "patient"])
        n_calls = len(observed)
        if not n_calls:
            continue

        per_site = observed.groupby("key").size()
        site_class = observed.drop_duplicates("key").set_index("key").wire_class

        # Drawn in chunks: the full matrix of draws by sites does not fit.
        null_counts = {c: [] for c in cfg.WIRE_CLASSES}
        chunk = 250
        for start in range(0, cfg.N_PERMUTATIONS, chunk):
            size = min(chunk, cfg.N_PERMUTATIONS - start)
            draws = rng.multinomial(n_calls, weights, size=size)
            recurrent_draw = draws >= 2
            for wire_class, mask in masks.items():
                null_counts[wire_class].append(
                    recurrent_draw[:, mask].sum(axis=1))
        null_counts = {c: np.concatenate(v) for c, v in null_counts.items()}

        for wire_class in cfg.WIRE_CLASSES:
            mask = masks[wire_class]
            keys = site_class[site_class == wire_class].index
            counts = per_site.reindex(keys).fillna(0)
            if counts.empty:
                continue
            recurrent = int((counts >= 2).sum())
            sites = int((counts > 0).sum())
            total = int(counts.sum())
            null = null_counts[wire_class]
            p = float((null >= recurrent).mean())
            rows.append(dict(
                cohort=cohort, wire_class=wire_class, calls=total, sites=sites,
                calls_per_site=round(total / sites, 3) if sites else np.nan,
                recurrent_sites=recurrent,
                expected_recurrent=round(float(null.mean()), 2),
                p_value=min(1.0, max(p, 1 / cfg.N_PERMUTATIONS)),
                max_patients_at_one_site=int(counts.max())))
    return pd.DataFrame(rows)


def recurrent_variants(universe, calls):
    """Substitutions of the two axis classes seen in several patients.

    Counts are pooled over the three tumour cohorts and a patient is counted
    once, so a sample appearing in two of them does not count twice.
    """
    annotation = {}
    wire = universe[universe.wire_class.isin(AXIS_CLASSES)
                    & (universe.kind == "missense")]
    for row in wire.itertuples():
        annotation[row.key] = row
    seen = {}
    for cohort, frame in calls.items():
        missense = frame[(frame.Variant_Classification == "Missense_Mutation")
                         & frame.usable & ~frame.polymorphism]
        for row in missense.itertuples():
            if row.key not in annotation:
                continue
            record = seen.setdefault(row.key, dict(patients=set(),
                                                   cohorts=set(), vafs=[]))
            record["patients"].add(row.patient)
            record["cohorts"].add(cohort)
            if pd.notna(row.vaf):
                record["vafs"].append(float(row.vaf))

    rows = []
    for key, record in seen.items():
        if len(record["patients"]) < MIN_PATIENTS:
            continue
        meta = annotation[key]
        position, ref, alt = key
        wt = AA3TO1.get(meta.wt_aa, meta.wt_aa)
        mutant = AA3TO1.get(meta.mut_aa, meta.mut_aa)
        vafs = pd.Series(record["vafs"], dtype=float)
        rows.append(dict(
            wire_class=meta.wire_class,
            variant=f"m.{position}{ref}>{alt}",
            protein=f"{meta.subunit} {wt}{int(meta.aa_pos)}{mutant}",
            patients=len(record["patients"]),
            cohorts=len(record["cohorts"]),
            median_vaf=round(float(vafs.median()), 3) if len(vafs) else np.nan))
    return (pd.DataFrame(rows)
            .sort_values(["patients", "variant"], ascending=[False, True])
            .reset_index(drop=True))


def clonality(calls):
    """Does the loading-site class reach fixation more often than its peers?"""
    rows, pooled = [], []
    for cohort, frame in calls.items():
        missense = frame[(frame.Variant_Classification == "Missense_Mutation")
                         & frame.usable & ~frame.polymorphism
                         & frame.vaf.notna()]
        truncating = frame[frame.Variant_Classification.isin(TRUNCATING)
                           & frame.vaf.notna()]
        groups = {"loading_site": missense[missense.wire_class == "loading_site"],
                  "cut_vertex": missense[missense.wire_class == "cut_vertex"],
                  "other missense": missense[missense.wire_class == "other"],
                  "truncating": truncating}
        reference = groups["other missense"].vaf
        for label, group in groups.items():
            if len(group) < 5:
                continue
            vaf = group.vaf
            p = (np.nan if label == "other missense"
                 else float(stats.mannwhitneyu(vaf, reference).pvalue))
            clonal = int((vaf >= CLONAL_VAF).sum())
            table = [[clonal, len(vaf) - clonal],
                     [int((reference >= CLONAL_VAF).sum()),
                      len(reference) - int((reference >= CLONAL_VAF).sum())]]
            p_clonal = (np.nan if label == "other missense"
                        else float(stats.fisher_exact(table)[1]))
            rows.append(dict(cohort=cohort, group=label, n=len(vaf),
                             median_vaf=round(float(vaf.median()), 3),
                             q90_vaf=round(float(vaf.quantile(0.9)), 3),
                             pct_clonal=round(100 * clonal / len(vaf), 1),
                             p_vs_other_missense=p,
                             p_clonal_vs_other_missense=p_clonal))
            pooled.append(group.assign(group=label, cohort=cohort))
    return pd.DataFrame(rows), pd.concat(pooled, ignore_index=True)


def main():
    rng = np.random.default_rng(cfg.RANDOM_SEED)
    universe = pd.read_parquet(cfg.RESULTS / "universe.parquet")
    universe["key"] = list(zip(universe.pos, universe.ref, universe.alt))
    with open(cfg.RESULTS / "spectra.pkl", "rb") as fh:
        spectra = pickle.load(fh)

    calls = load_calls(universe)
    table = recurrence(universe, spectra, calls, rng)
    print("recurrence of individual substitutions "
          f"({cfg.N_PERMUTATIONS:,} multinomial draws)")
    print(table.to_string(index=False))

    recurrent = recurrent_variants(universe, calls)
    print(f"\nsubstitutions seen in at least {MIN_PATIENTS} patients")
    print(recurrent.to_string(index=False))

    clones, detail = clonality(calls)
    print("\nallele fractions and clonality")
    print(clones.to_string(index=False))

    table.to_csv(cfg.RESULTS / "recurrence.csv", index=False)
    recurrent.to_csv(cfg.RESULTS / "recurrent_variants.csv", index=False)
    clones.to_csv(cfg.RESULTS / "clonality.csv", index=False)
    detail[["cohort", "group", "key", "patient", "vaf"]].to_csv(
        cfg.RESULTS / "clonality_calls.csv", index=False)
    print(f"\nwrote {cfg.RESULTS / 'recurrence.csv'} and "
          f"{cfg.RESULTS / 'clonality.csv'}")


if __name__ == "__main__":
    main()
```
