## Supplementary code for "Cancers relax selection on charge at the mitochondrial complex I quinone cavity": 11_haplogroup.py

```
"""Haplogroup-defining variants: constraint over evolutionary time.

The germline cohort measures depletion in the present population. Variants
that define branches of the human mitochondrial phylogeny are a different
object: substitutions that reached fixation along a lineage and have been
carried for tens of thousands of years. If charge at a proton loading site is
under purifying selection, those positions should be close to absent from the
set of branch-defining variants, which gives the depletion a timescale.

Branch-defining substitutions are read from PhyloTree build 17 as distributed
with HaploGrep. Reversions, indels, heteroplasmic calls and positions in the
control region are discarded, leaving unambiguous substitutions.
"""
import pickle
import re
import sys
import xml.etree.ElementTree as ET
from pathlib import Path

import numpy as np
import pandas as pd
from scipy import stats

sys.path.insert(0, str(Path(__file__).resolve().parents[1]))
import config as cfg
from src import variants

POLY = re.compile(r"^(\d+)([ACGT])$")


def haplogroup_variants(path, seq):
    """Branch-defining substitutions, with the branch count for each."""
    tree = ET.parse(path)
    counts, branches = {}, 0
    for node in tree.iter("haplogroup"):
        name = node.get("name")
        details = node.find("details")
        if details is None:
            continue
        branches += 1
        for poly in details.findall("poly"):
            token = (poly.text or "").strip()
            # '!' marks a reversion, '(' a dubious call, '.' an insertion,
            # 'd' a deletion; lower case marks a heteroplasmy.
            if not token or any(c in token for c in "!().d"):
                continue
            match = POLY.match(token)
            if not match:
                continue
            pos, alt = int(match.group(1)), match.group(2)
            if not 2 <= pos <= 16568:
                continue
            ref = seq[pos - 1]
            if ref == alt or ref not in "ACGT":
                continue
            key = (pos, ref, alt)
            entry = counts.setdefault(key, {"branches": 0, "names": []})
            entry["branches"] += 1
            if len(entry["names"]) < 4:
                entry["names"].append(name)
    print(f"PhyloTree 17: {branches:,} branches, "
          f"{len(counts):,} distinct branch-defining substitutions")
    return counts


def main():
    seq = variants.read_rcrs(cfg.RCRS_FASTA)
    defining = haplogroup_variants(cfg.DATA / "phylotree17_FU1a.xml", seq)

    universe = pd.read_parquet(cfg.RESULTS / "universe.parquet")
    universe["key"] = list(zip(universe.pos, universe.ref, universe.alt))
    with open(cfg.RESULTS / "spectra.pkl", "rb") as fh:
        spectra = pickle.load(fh)

    universe["haplogroup_branches"] = [defining.get(k, {}).get("branches", 0)
                                       for k in universe.key]
    universe["haplogroup"] = universe.haplogroup_branches > 0

    rows = []
    for kind in ("missense", "synonymous"):
        subset = universe[(universe.kind == kind) & universe.identical]
        weights = subset.channel.map(spectra["germline"])
        weights = weights.fillna(weights.min())
        total = int(subset.haplogroup.sum())
        print(f"\n{kind} branch-defining substitutions in the ND genes: {total}")
        for wire_class in cfg.WIRE_CLASSES:
            mask = (subset.wire_class == wire_class).to_numpy()
            expected = total * weights[mask].sum() / weights.sum()
            observed = int(subset.loc[mask, "haplogroup"].sum())
            p = (stats.poisson.sf(observed - 1, expected) * 2
                 if observed > expected else stats.poisson.cdf(observed, expected) * 2)
            rows.append(dict(kind=kind, wire_class=wire_class,
                             n_sites=int(mask.sum()), observed=observed,
                             expected=round(float(expected), 2),
                             obs_over_exp=round(observed / expected, 3)
                             if expected else np.nan,
                             p_value=min(1.0, float(p))))
    table = pd.DataFrame(rows)
    print("\nbranch-defining variants against the proton wire")
    print(table.to_string(index=False))

    # The three timescales side by side, for the missense set.
    missense = universe[(universe.kind == "missense") & universe.identical]
    oe = pd.read_csv(cfg.RESULTS / "oe_missense.csv")
    summary = []
    for wire_class in cfg.WIRE_CLASSES:
        row = {"wire_class": wire_class}
        hit = table[(table.kind == "missense") & (table.wire_class == wire_class)]
        row["phylogeny"] = float(hit.obs_over_exp.iloc[0])
        for cohort in cfg.COHORTS:
            value = oe[(oe.cohort == cohort) & (oe.wire_class == wire_class)]
            row[cohort] = float(value.obs_over_exp.iloc[0])
        summary.append(row)
    summary = pd.DataFrame(summary)
    print("\nobserved/expected across three timescales")
    print(summary.to_string(index=False))

    # Which loading-site substitutions did reach fixation on a branch?
    fixed = missense[missense.haplogroup
                     & (missense.wire_class == "loading_site")].copy()
    fixed["branches"] = fixed.key.map(
        lambda k: defining.get(k, {}).get("branches", 0))
    fixed["haplogroups"] = fixed.key.map(
        lambda k: ", ".join(defining.get(k, {}).get("names", [])))
    fixed = fixed.sort_values("branches", ascending=False)
    print(f"\nloading-site substitutions carried by a branch: {len(fixed)}")
    if len(fixed):
        show = fixed[["subunit", "wt_aa", "aa_pos", "mut_aa", "pos", "ref",
                      "alt", "branches", "haplogroups"]].head(12)
        print(show.to_string(index=False))

    # How large an electrostatic shift does a substitution that fixed on a
    # branch impose, compared with one that recurs in tumours?
    scores = pd.read_parquet(cfg.RESULTS / "axis_scores.parquet")
    mapping = pd.read_csv(cfg.RESULTS / "mouse_human_map.csv")
    mapping = mapping[mapping.transferable & mapping.human_pos.notna()]
    to_human = {(r.subunit, int(r.mouse_pos)): int(r.human_pos)
                for r in mapping.itertuples()}
    scores["human_pos"] = [to_human.get((s, p)) for s, p in
                           zip(scores.subunit, scores.pos)]
    axis_a = {(r.subunit, int(r.human_pos), r.mut_aa): r.axis_a_charge
              for r in scores[scores.human_pos.notna()].itertuples()}

    def values(frame):
        out = [axis_a.get((s, int(p), m)) for s, p, m in
               zip(frame.subunit, frame.aa_pos, frame.mut_aa)]
        return np.array([v for v in out if v is not None], float)

    recurrent_keys = set()
    recurrent_path = cfg.RESULTS / "recurrence.csv"
    calls_path = cfg.RESULTS / "clonality_calls.csv"
    if calls_path.exists():
        calls = pd.read_csv(calls_path)
        calls = calls[calls.group == "loading_site"]
        counted = calls.groupby("key").patient.nunique()
        recurrent_keys = {eval(k) for k in counted[counted >= 4].index}

    groups = [("branch-defining", fixed),
              ("tumour-recurrent", missense[missense.key.isin(recurrent_keys)]),
              ("all possible", missense[missense.wire_class == "loading_site"])]
    shift_rows = []
    for label, frame in groups:
        v = values(frame)
        shift_rows.append(dict(group=label, n=len(v),
                               mean_axis_a=round(float(v.mean()), 4)
                               if len(v) else np.nan,
                               median_axis_a=round(float(np.median(v)), 4)
                               if len(v) else np.nan,
                               pct_above_threshold=round(
                                   100 * float((v >= cfg.AXIS_A_THRESHOLD).mean()), 1)
                               if len(v) else np.nan))
    shifts = pd.DataFrame(shift_rows)
    branch, recurrent = values(groups[0][1]), values(groups[1][1])
    if len(branch) and len(recurrent):
        p = float(stats.mannwhitneyu(branch, recurrent).pvalue)
        shifts["p_branch_vs_recurrent"] = [p] + [np.nan] * (len(shifts) - 1)
        print(f"\nelectrostatic shift imposed at a loading site, by group "
              f"(Mann-Whitney P = {p:.2g})")
    print(shifts.to_string(index=False))
    shifts.to_csv(cfg.RESULTS / "axis_a_by_timescale.csv", index=False)

    table.to_csv(cfg.RESULTS / "haplogroup_enrichment.csv", index=False)
    summary.to_csv(cfg.RESULTS / "timescales.csv", index=False)
    fixed[["subunit", "wt_aa", "aa_pos", "mut_aa", "pos", "ref", "alt",
           "branches", "haplogroups"]].to_csv(
        cfg.RESULTS / "haplogroup_loading_sites.csv", index=False)
    print(f"\nwrote {cfg.RESULTS / 'haplogroup_enrichment.csv'} and "
          f"{cfg.RESULTS / 'timescales.csv'}")


if __name__ == "__main__":
    main()
```
