## Supplementary code for "Cancers relax selection on charge at the mitochondrial complex I quinone cavity": 12_local_structure.py

```
"""Per-residue structural concordance between the two species.

Restricting the analysis to codons at which mouse and human carry the same
amino acid asks about sequence. The inference being transferred is structural,
so the stronger filter asks whether the residue sits in the same place. For
each mapped residue the surrounding Ca atoms are superposed locally, and two
quantities are recorded: how well that local environment agrees, and how far
the side-chain centroid moves once it does. The selection analysis is then
repeated on the residues that agree by both measures.
"""
import pickle
import sys
from pathlib import Path

import numpy as np
import pandas as pd
from scipy.spatial import cKDTree

sys.path.insert(0, str(Path(__file__).resolve().parents[1]))
import config as cfg
from src import variants, wire

PATCH_RADIUS = 10.0        # A, Ca atoms entering the local superposition
MIN_PATCH = 5              # residues, fewer than this is not a superposition
LOCAL_RMSD_MAX = 1.0       # A
SIDE_CHAIN_MAX = 2.0       # A


def alpha_carbons(model, chains):
    out = {}
    for chain in model:
        if chain.name not in chains:
            continue
        for residue in chain:
            atom = residue.find_atom("CA", "*")
            if atom:
                out[(chains[chain.name], residue.seqid.num)] = np.array(
                    [atom.pos.x, atom.pos.y, atom.pos.z])
    return out


def kabsch(P, Q):
    cp, cq = P.mean(0), Q.mean(0)
    U, _, Vt = np.linalg.svd((P - cp).T @ (Q - cq))
    d = np.sign(np.linalg.det(Vt.T @ U.T))
    R = Vt.T @ np.diag([1.0, 1.0, d]) @ U.T
    return R, cq - R @ cp


def concordance(mouse_model, human_model, mapping):
    mouse_ca = alpha_carbons(mouse_model, cfg.MOUSE_CHAINS)
    human_ca = alpha_carbons(human_model, cfg.HUMAN_CHAINS)
    mouse_geom = wire.residue_geometry(mouse_model, cfg.MOUSE_CHAINS)
    human_geom = wire.residue_geometry(human_model, cfg.HUMAN_CHAINS)

    pairs = {(r.subunit, int(r.mouse_pos)): (r.subunit, int(r.human_pos))
             for r in mapping.itertuples()}
    usable = [k for k in pairs if k in mouse_ca and pairs[k] in human_ca]
    keys = list(usable)
    coords = np.asarray([mouse_ca[k] for k in keys])
    tree = cKDTree(coords)

    rows = []
    for i, key in enumerate(keys):
        patch = [keys[j] for j in tree.query_ball_point(coords[i], PATCH_RADIUS)]
        if len(patch) < MIN_PATCH:
            continue
        P = np.asarray([mouse_ca[k] for k in patch])
        Q = np.asarray([human_ca[pairs[k]] for k in patch])
        R, t = kabsch(P, Q)
        moved = P @ R.T + t
        local_rmsd = float(np.sqrt(((moved - Q) ** 2).sum(1).mean()))

        deviation = np.nan
        mouse_side = mouse_geom.get(key)
        human_side = human_geom.get(pairs[key])
        if mouse_side is not None and human_side is not None:
            centroid = np.asarray(mouse_side["centroid"]) @ R.T + t
            deviation = float(np.linalg.norm(
                centroid - np.asarray(human_side["centroid"])))
        rows.append(dict(subunit=key[0], mouse_pos=key[1],
                         human_pos=pairs[key][1], patch=len(patch),
                         local_rmsd=round(local_rmsd, 3),
                         side_chain_deviation=round(deviation, 3)
                         if np.isfinite(deviation) else np.nan))
    table = pd.DataFrame(rows)
    table["concordant"] = ((table.local_rmsd <= LOCAL_RMSD_MAX)
                           & (table.side_chain_deviation.fillna(1e3)
                              <= SIDE_CHAIN_MAX))
    return table


def missense_oe(universe, spectra, keep):
    subset = universe[(universe.kind == "missense") & universe.identical]
    subset = subset[[(s, p) in keep for s, p in
                     zip(subset.subunit, subset.aa_pos)]]
    rows = []
    for cohort in cfg.COHORTS:
        d = subset if cohort == "germline" else subset[~subset.polymorphism]
        for wire_class in cfg.WIRE_CLASSES:
            mask = (d.wire_class == wire_class).to_numpy()
            obs, exp, oe, p = variants.observed_expected(
                d, spectra[cohort], f"n_{cohort}_missense", mask)
            rows.append(dict(cohort=cohort, wire_class=wire_class,
                             n_substitutions=int(mask.sum()), observed=obs,
                             expected=round(exp, 1) if np.isfinite(exp) else np.nan,
                             obs_over_exp=round(oe, 3) if np.isfinite(oe) else np.nan,
                             p_value=p))
    return pd.DataFrame(rows)


def main():
    mapping = pd.read_csv(cfg.RESULTS / "mouse_human_map.csv")
    mapping = mapping[mapping.transferable & mapping.human_pos.notna()]

    mouse_model = wire.load_model(cfg.MOUSE_CIF, cfg.MOUSE_CHAINS)
    human_model = wire.load_model(cfg.HUMAN_CIF, cfg.HUMAN_CHAINS)
    table = concordance(mouse_model, human_model, mapping)

    classes = {(r.subunit, int(r.mouse_pos)): r.wire_class
               for r in mapping.itertuples()}
    identical = {(r.subunit, int(r.mouse_pos)): bool(r.identical)
                 for r in mapping.itertuples()}
    table["wire_class"] = [classes.get((s, p), "other") for s, p in
                           zip(table.subunit, table.mouse_pos)]
    table["identical"] = [identical.get((s, p), False) for s, p in
                          zip(table.subunit, table.mouse_pos)]

    print(f"residues with a local superposition: {len(table):,}")
    print(f"median local Ca RMSD {table.local_rmsd.median():.2f} A, "
          f"median side-chain deviation "
          f"{table.side_chain_deviation.median():.2f} A")
    print(f"\nconcordant at <= {LOCAL_RMSD_MAX} A local RMSD and "
          f"<= {SIDE_CHAIN_MAX} A side-chain deviation: "
          f"{int(table.concordant.sum()):,} of {len(table):,} "
          f"({100 * table.concordant.mean():.1f}%)")
    summary = (table.groupby("wire_class")
               .agg(n=("concordant", "size"),
                    identical=("identical", "sum"),
                    concordant=("concordant", "sum"),
                    both=("concordant", lambda s: int((s & table.loc[s.index,
                                                       "identical"]).sum())),
                    median_local_rmsd=("local_rmsd", "median"),
                    median_deviation=("side_chain_deviation", "median"))
               .reindex(list(cfg.WIRE_CLASSES)))
    print("\nconcordance by wire class")
    print(summary.to_string())

    universe = pd.read_parquet(cfg.RESULTS / "universe.parquet")
    with open(cfg.RESULTS / "spectra.pkl", "rb") as fh:
        spectra = pickle.load(fh)

    keep = {(r.subunit, int(r.human_pos)) for r in
            table[table.concordant & table.identical].itertuples()}
    restricted = missense_oe(universe, spectra, keep)
    print(f"\nmissense selection on the {len(keep):,} structurally concordant "
          f"codons")
    print(restricted.pivot(index="wire_class", columns="cohort",
                           values=["obs_over_exp", "p_value"])
          .reindex(list(cfg.WIRE_CLASSES)).round(4).to_string())

    primary = pd.read_csv(cfg.RESULTS / "oe_missense.csv")
    comparison = (primary.rename(columns={"obs_over_exp": "identity_filter"})
                  [["cohort", "wire_class", "identity_filter"]]
                  .merge(restricted.rename(
                      columns={"obs_over_exp": "concordance_filter"})
                      [["cohort", "wire_class", "concordance_filter"]],
                      on=["cohort", "wire_class"]))
    print("\nobserved/expected under the two filters")
    print(comparison[comparison.wire_class != "other"].to_string(index=False))

    table.to_csv(cfg.RESULTS / "structural_concordance.csv", index=False)
    restricted.to_csv(cfg.RESULTS / "oe_missense_concordant.csv", index=False)
    comparison.to_csv(cfg.RESULTS / "filter_comparison.csv", index=False)
    print(f"\nwrote {cfg.RESULTS / 'structural_concordance.csv'}")


if __name__ == "__main__":
    main()
```
