## Supplementary code for "Cancers relax selection on charge at the mitochondrial complex I quinone cavity": 13_interaction.py

```
"""A single test of the claim the paper makes.

The observed/expected tables show depletion at loading sites in the germline
and enrichment in tumours, but comparing two ratios is not a test of the
difference between them. This script fits the germline and tumour counts in one
model and tests the wire class by tissue-type interaction directly, so that the
central claim — that constraint on loading-site charge is relaxed in tumours —
has one coefficient and one P value.

Counts for the same substitution appear once per cohort, so the standard errors
are clustered on the substitution. The polymorphism filter cannot be applied
to the two arms in the same way (common germline variants are the tolerated
ones, and excluding them would remove the germline signal being measured), so
the model is fitted both with and without it.
"""
import pickle
import sys
from pathlib import Path

import numpy as np
import pandas as pd
import statsmodels.api as sm
import statsmodels.formula.api as smf

sys.path.insert(0, str(Path(__file__).resolve().parents[1]))
import config as cfg

TUMOUR_COHORTS = ("TCGA", "PCAWG", "MSK-IMPACT")
# The tumour indicator is a linear combination of the cohort dummies, so it
# is not entered as a main effect: the cohort terms absorb it. The
# interactions are built as explicit indicators, one per class, leaving the
# reference class out so the design stays full rank.
INTERACTIONS = ("x_loading_site", "x_cut_vertex", "x_cluster_member")
BASE = ("y ~ C(wire_class, Treatment('other')) + C(cohort) + "
        + " + ".join(INTERACTIONS))
ADJUSTED = (BASE + " + phastcons + burial + delta_volume + delta_hydropathy")
# Cluster members lie close to loading sites, so the same relaxation can
# reach them through proximity rather than through membership.
WITH_DISTANCE = ADJUSTED + " + dist_to_loading_site"


def long_format(universe, spectra, drop_polymorphisms):
    """One row per substitution and cohort, with the expected count as offset."""
    subset = universe[(universe.kind == "missense") & universe.identical]
    if drop_polymorphisms:
        subset = subset[~subset.polymorphism]
    frames = []
    for cohort in cfg.COHORTS:
        weights = subset.channel.map(spectra[cohort])
        weights = weights.fillna(weights.min())
        counts = subset[f"n_{cohort}_missense"].to_numpy()
        expected = counts.sum() * (weights / weights.sum()).to_numpy()
        frame = subset[["subunit", "aa_pos", "pos", "ref", "alt", "wire_class",
                        "phastcons", "burial", "delta_volume",
                        "delta_hydropathy", "dist_to_loading_site"]].copy()
        frame["cohort"] = cohort
        frame["tumour"] = int(cohort != "germline")
        frame["y"] = counts
        frame["offset"] = np.log(np.maximum(expected, 1e-12))
        for wire_class in ("loading_site", "cut_vertex", "cluster_member"):
            frame[f"x_{wire_class}"] = ((frame.wire_class == wire_class)
                                        * frame.tumour).astype(float)
        frames.append(frame)
    return pd.concat(frames, ignore_index=True)


def fit(data, formula):
    model = smf.glm(formula, data=data, family=sm.families.Poisson(),
                    offset=data["offset"])
    groups = data.subunit + ":" + data.pos.astype(str) + data.alt
    return model.fit(scale="X2", cov_type="cluster",
                     cov_kwds={"groups": groups.to_numpy()})


def report(fit_result, label, n_rows):
    rows = []
    for wire_class in ("loading_site", "cut_vertex", "cluster_member"):
        main = f"C(wire_class, Treatment('other'))[T.{wire_class}]"
        inter = f"x_{wire_class}"
        if inter not in fit_result.params:
            continue
        low, high = np.exp(fit_result.conf_int().loc[inter])
        rows.append(dict(
            model=label, wire_class=wire_class, n_rows=n_rows,
            germline_rate_ratio=round(float(np.exp(fit_result.params[main])), 3),
            interaction_ratio=round(float(np.exp(fit_result.params[inter])), 3),
            ci_low=round(float(low), 3), ci_high=round(float(high), 3),
            p_value=float(fit_result.pvalues[inter])))
    return rows


def main():
    universe = pd.read_parquet(cfg.RESULTS / "universe_annotated.parquet")
    with open(cfg.RESULTS / "spectra.pkl", "rb") as fh:
        spectra = pickle.load(fh)

    rows = []
    for drop in (False, True):
        label_suffix = ("polymorphisms excluded from both arms" if drop
                        else "all identical codons")
        data = long_format(universe, spectra, drop)
        rows += report(fit(data, BASE), f"unadjusted, {label_suffix}", len(data))

        adjusted = data[data.phastcons.notna() & data.burial.notna()]
        rows += report(fit(adjusted, ADJUSTED),
                       f"adjusted, {label_suffix}", len(adjusted))
        with_distance = adjusted[adjusted.dist_to_loading_site.notna()]
        rows += report(fit(with_distance, WITH_DISTANCE),
                       f"adjusted + distance, {label_suffix}",
                       len(with_distance))

    table = pd.DataFrame(rows)
    print("wire class by tissue-type interaction")
    print("germline_rate_ratio: the class against other ND residues in the "
          "germline")
    print("interaction_ratio:   how much that ratio changes in tumours\n")
    print(table.to_string(index=False))

    primary = table[(table.wire_class == "loading_site")
                    & table.model.str.startswith("unadjusted, all")]
    if len(primary):
        row = primary.iloc[0]
        print(f"\nloading sites: the germline rate ratio of "
              f"{row.germline_rate_ratio} is multiplied by "
              f"{row.interaction_ratio} in tumours "
              f"(95% CI {row.ci_low} to {row.ci_high}, P = {row.p_value:.2g})")

    table.to_csv(cfg.RESULTS / "interaction.csv", index=False)
    print(f"\nwrote {cfg.RESULTS / 'interaction.csv'}")


if __name__ == "__main__":
    main()
```
