## Supplementary code for "Cancers relax selection on charge at the mitochondrial complex I quinone cavity": 14_figure_s7.py

```
"""Supplementary Figure 7: structural concordance, timescales and recurrence."""
import sys
from pathlib import Path

import numpy as np
import pandas as pd
import matplotlib.pyplot as plt

sys.path.insert(0, str(Path(__file__).resolve().parents[1]))
import config as cfg
from src import plotting
from src.plotting import CLASS_COLOURS, CLASS_LABELS

SCORED = ("loading_site", "cut_vertex", "cluster_member")


def main():
    plotting.use_style()
    concordance = pd.read_csv(cfg.RESULTS / "structural_concordance.csv")
    comparison = pd.read_csv(cfg.RESULTS / "filter_comparison.csv")
    timescales = pd.read_csv(cfg.RESULTS / "timescales.csv")
    shifts = pd.read_csv(cfg.RESULTS / "axis_a_by_timescale.csv")
    recurrence = pd.read_csv(cfg.RESULTS / "recurrence.csv")

    fig, axes_ = plt.subplots(2, 2, figsize=(11.0, 8.2))

    # a) how well the two structures agree, residue by residue
    ax = axes_[0, 0]
    bins = np.linspace(0, 3, 46)
    ax.hist(concordance.local_rmsd.clip(upper=3), bins=bins, color="#5dade2",
            alpha=0.85, label="Local Cα RMSD")
    ax.hist(concordance.side_chain_deviation.dropna().clip(upper=3), bins=bins,
            histtype="step", lw=1.8, color=CLASS_COLOURS["loading_site"],
            label="Side-chain centroid deviation")
    ax.axvline(1.0, color="black", lw=0.9, ls="--")
    ax.axvline(2.0, color="black", lw=0.9, ls=":")
    ax.set_xlabel("Deviation after local superposition (Å)")
    ax.set_ylabel("Residues")
    ax.set_title(f"Median {concordance.local_rmsd.median():.2f} Å and "
                 f"{concordance.side_chain_deviation.median():.2f} Å",
                 fontsize=11)
    ax.set_xlim(0, 3.05)
    ax.set_xticks([0, 0.5, 1.0, 1.5, 2.0, 2.5, 3.0])
    ax.set_xticklabels(["0", "0.5", "1.0", "1.5", "2.0", "2.5", "≥3"])
    ax.legend(fontsize=9)
    plotting.panel_label(ax, "a")

    # b) the primary result under the sequence and the structure filter
    ax = axes_[0, 1]
    cohorts = list(cfg.COHORTS)
    loading = comparison[comparison.wire_class == "loading_site"].set_index(
        "cohort").reindex(cohorts)
    x = np.arange(len(cohorts))
    ax.bar(x - 0.19, loading.identity_filter, width=0.38, color="#aab7b8",
           label="Identical in sequence")
    ax.bar(x + 0.19, loading.concordance_filter, width=0.38,
           color=CLASS_COLOURS["loading_site"],
           label="Also structurally concordant")
    ax.axhline(1.0, color="black", lw=0.8, ls=":")
    ax.set_xticks(x)
    ax.set_xticklabels(cohorts, rotation=20, ha="right")
    ax.set_ylabel("Observed / expected, loading sites")
    ax.legend(fontsize=9)
    plotting.panel_label(ax, "b")

    # c) three timescales
    ax = axes_[1, 0]
    columns = ["phylogeny", "germline", "TCGA", "PCAWG", "MSK-IMPACT"]
    labels = ["PhyloTree\nbranches", "Germline", "TCGA", "PCAWG", "MSK-\nIMPACT"]
    table = timescales.set_index("wire_class")
    for wire_class in SCORED:
        ax.plot(np.arange(len(columns)), table.loc[wire_class, columns], "o-",
                ms=6, color=CLASS_COLOURS[wire_class],
                label=CLASS_LABELS[wire_class])
    ax.axhline(1.0, color="black", lw=0.8, ls=":")
    ax.axvline(1.5, color="#d5d8dc", lw=8, zorder=0)
    ax.set_xticks(np.arange(len(columns)))
    ax.set_xticklabels(labels, fontsize=9)
    ax.set_ylabel("Observed / expected")
    ax.set_title("Evolutionary time to somatic time", fontsize=11)
    ax.legend(fontsize=9)
    plotting.panel_label(ax, "c")

    # d) the size of the electrostatic shift, by group
    ax = axes_[1, 1]
    order = ["branch-defining", "tumour-recurrent", "all possible"]
    rows = shifts.set_index("group").reindex(order)
    colours = [CLASS_COLOURS["cut_vertex"], CLASS_COLOURS["loading_site"],
               "#aab7b8"]
    ax.bar(np.arange(len(order)), rows.mean_axis_a, color=colours, width=0.6)
    for i, (value, n) in enumerate(zip(rows.mean_axis_a, rows.n)):
        ax.text(i, value + rows.mean_axis_a.max() * 0.03,
                f"{value:.3f}\n(n = {int(n)})", ha="center", fontsize=9)
    ax.set_xticks(np.arange(len(order)))
    ax.set_xticklabels([o.replace("-", "-\n") for o in order], fontsize=9)
    ax.set_ylabel("Mean shift at a loading site (eV)")
    ax.set_ylim(0, rows.mean_axis_a.max() * 1.25)
    p = shifts.p_branch_vs_recurrent.dropna()
    if len(p):
        ax.set_title(f"Branch-defining versus recurrent, "
                     f"P = {float(p.iloc[0]):.1g}", fontsize=11)
    plotting.panel_label(ax, "d")

    fig.tight_layout()
    path = plotting.save(fig, cfg.FIGURES, "SuppFig7")
    print("wrote", path, "and", path.with_suffix(".png"))

    loading_rec = recurrence[recurrence.wire_class == "loading_site"]
    print("\nrecurrence, loading sites")
    print(loading_rec[["cohort", "recurrent_sites", "expected_recurrent",
                       "p_value"]].to_string(index=False))


if __name__ == "__main__":
    main()
```
