## Supplementary code for "Cancers relax selection on charge at the mitochondrial complex I quinone cavity": 15_coverage.py

```
"""Is the loading-site enrichment an artefact of sequencing coverage?

A position that is sequenced in more samples has more opportunity to be called
mutated, and the wire classes are not spread evenly over the genes. The
coverage matrices distributed with the variant calls give, for every sample and
every base of the mitochondrial genome, whether that base was covered. This
script compares coverage between the wire classes and then refits the selection
model with per-position coverage as a covariate.
"""
import pickle
import sys
from pathlib import Path

import numpy as np
import pandas as pd
import statsmodels.api as sm
import statsmodels.formula.api as smf
from scipy import stats

sys.path.insert(0, str(Path(__file__).resolve().parents[1]))
import config as cfg

MATRICES = {"TCGA": "coverage_matrix_tcga.txt.gz",
            "MSK-IMPACT": "coverage_matrix_impact.txt.gz"}
CHUNK = 200


def position_coverage(filename):
    """Fraction of samples covering each base, read in chunks."""
    total = counted = None
    samples = 0
    reader = pd.read_csv(cfg.MTDNA / "original_data" / filename, sep="\t",
                         chunksize=CHUNK, low_memory=False)
    for chunk in reader:
        values = chunk.drop(columns=["Tumor_Sample_Barcode"]).to_numpy(
            dtype=np.float32)
        # Some samples carry no entry at some bases; those are not counted
        # rather than treated as uncovered.
        present = np.isfinite(values)
        covered = np.where(present, values, 0.0).sum(0)
        total = covered if total is None else total + covered
        counted = present.sum(0) if counted is None else counted + present.sum(0)
        samples += len(chunk)
        columns = chunk.columns[1:]
    fraction = pd.Series(total / np.maximum(counted, 1),
                         index=[int(c) for c in columns])
    return fraction, samples


def main():
    universe = pd.read_parquet(cfg.RESULTS / "universe_annotated.parquet")
    with open(cfg.RESULTS / "spectra.pkl", "rb") as fh:
        spectra = pickle.load(fh)

    rows, models = [], []
    for cohort, filename in MATRICES.items():
        fraction, samples = position_coverage(filename)
        print(f"{cohort}: {samples:,} samples, per-base coverage "
              f"{fraction.min():.3f} to {fraction.max():.3f} "
              f"(median {fraction.median():.3f})")

        subset = universe[(universe.kind == "missense") & universe.identical
                          & ~universe.polymorphism].copy()
        subset["coverage"] = subset.pos.map(fraction)
        missing = int(subset.coverage.isna().sum())
        if missing:
            print(f"  positions without a coverage entry: {missing}")
        subset = subset[subset.coverage.notna()]

        reference = subset.loc[subset.wire_class == "other", "coverage"]
        for wire_class in cfg.WIRE_CLASSES:
            values = subset.loc[subset.wire_class == wire_class, "coverage"]
            p = (np.nan if wire_class == "other"
                 else float(stats.mannwhitneyu(values, reference).pvalue))
            rows.append(dict(cohort=cohort, wire_class=wire_class,
                             n_positions=len(values),
                             mean_coverage=round(float(values.mean()), 4),
                             median_coverage=round(float(values.median()), 4),
                             p_vs_other=p))

        # The selection model, with and without coverage as a covariate.
        weights = subset.channel.map(spectra[cohort])
        weights = weights.fillna(weights.min())
        subset["offset"] = np.log(weights / weights.mean())
        subset["y"] = subset[f"n_{cohort}_missense"]
        subset["wire_class"] = pd.Categorical(
            subset.wire_class, categories=["other", "cluster_member",
                                           "cut_vertex", "loading_site"])
        subset["log_coverage"] = np.log(np.maximum(subset.coverage, 1e-3))
        if subset.coverage.median() < 0.05:
            print(f"  {cohort} is a targeted panel and mitochondrial coverage "
                  f"is incidental; the adjusted model is not fitted")
            continue
        for label, formula in [
                ("unadjusted", "y ~ C(wire_class)"),
                ("coverage-adjusted", "y ~ C(wire_class) + log_coverage")]:
            fit = smf.glm(formula, data=subset, family=sm.families.Poisson(),
                          offset=subset["offset"]).fit(scale="X2")
            for wire_class in ("loading_site", "cut_vertex", "cluster_member"):
                term = f"C(wire_class)[T.{wire_class}]"
                low, high = np.exp(fit.conf_int().loc[term])
                models.append(dict(cohort=cohort, model=label,
                                   wire_class=wire_class,
                                   rate_ratio=round(float(np.exp(
                                       fit.params[term])), 3),
                                   ci_low=round(float(low), 3),
                                   ci_high=round(float(high), 3),
                                   p_value=float(fit.pvalues[term])))

    coverage = pd.DataFrame(rows)
    print("\ncoverage by wire class")
    print(coverage.to_string(index=False))

    fits = pd.DataFrame(models)
    print("\nrate ratios before and after adjusting for coverage")
    print(fits.pivot(index=["cohort", "wire_class"], columns="model",
                     values="rate_ratio").to_string())

    coverage.to_csv(cfg.RESULTS / "coverage_by_class.csv", index=False)
    fits.to_csv(cfg.RESULTS / "coverage_adjusted.csv", index=False)
    print(f"\nwrote {cfg.RESULTS / 'coverage_by_class.csv'}")


if __name__ == "__main__":
    main()
```
