## Supplementary code for "Cancers relax selection on charge at the mitochondrial complex I quinone cavity": 16_strata.py

```
"""Is the enrichment carried by a few cancer types, or by a few allele
fractions?

Two questions are answered here. First, whether loading-site enrichment is
spread across tumour types or produced by the handful of lineages that carry
most mitochondrial mutations: each cancer type is scored against the same
spectrum-weighted null, heterogeneity is tested by Cochran's Q, and the pooled
estimate is recomputed with each type removed in turn. Second, whether the
depletion of cluster members seen only in PCAWG is confined to a range of
allele fractions.
"""
import pickle
import sys
from pathlib import Path

import numpy as np
import pandas as pd
from scipy import stats

sys.path.insert(0, str(Path(__file__).resolve().parents[1]))
import config as cfg

COHORT_FILES = {"TCGA": ("data_mutations_tcga.txt.gz", "data_samples_tcga.txt.gz"),
                "PCAWG": ("data_mutations_pcawg.txt.gz", "data_samples_pcawg.txt.gz"),
                "MSK-IMPACT": ("data_mutations_impact.txt.gz",
                               "data_samples_impact.txt.gz")}
MIN_CALLS = 20
VAF_BANDS = [("< 0.1", 0.0, 0.1), ("0.1 to 0.5", 0.1, 0.5), (">= 0.5", 0.5, 1.01)]


def eligible(universe, cohort):
    subset = universe[(universe.kind == "missense") & universe.identical
                      & ~universe.polymorphism]
    return subset


def class_share(subset, spectrum, wire_class):
    weights = subset.channel.map(spectrum)
    weights = weights.fillna(weights.min())
    mask = (subset.wire_class == wire_class).to_numpy()
    return float(weights[mask].sum() / weights.sum())


def load(cohort, universe):
    mutations, samples = COHORT_FILES[cohort]
    calls = pd.read_csv(cfg.MTDNA / "original_data" / mutations, sep="\t",
                        low_memory=False)
    meta = pd.read_csv(cfg.MTDNA / "original_data" / samples, sep="\t",
                       low_memory=False)
    for candidate in ("CANCER_TYPE", "maintype", "dcc_project_code"):
        if candidate in meta.columns:
            column = candidate
            break
    else:
        raise KeyError(f"no cancer-type column in {samples}")
    types = dict(zip(meta.Tumor_Sample_Barcode, meta[column]))

    lookup = dict(zip(universe.key, universe.wire_class))
    usable = dict(zip(universe.key, universe.identical))
    polymorphic = dict(zip(universe.key, universe.polymorphism))
    calls = calls[calls.Hugo_Symbol.isin(sorted(universe.gene.unique()))
                  & (calls.Variant_Classification == "Missense_Mutation")].copy()
    calls["key"] = list(zip(calls.Start_Position, calls.Reference_Allele,
                            calls.Tumor_Seq_Allele2))
    calls["wire_class"] = calls.key.map(lookup)
    calls["vaf"] = pd.to_numeric(calls.TumorVAF, errors="coerce")
    calls["cancer_type"] = calls.Tumor_Sample_Barcode.map(types)
    calls = calls[calls.key.map(usable).fillna(False)
                  & ~calls.key.map(polymorphic).fillna(False)]
    return calls


def by_cancer_type(calls, share, cohort, wire_class):
    rows = []
    for cancer_type, stratum in calls.groupby("cancer_type"):
        if len(stratum) < MIN_CALLS:
            continue
        observed = int((stratum.wire_class == wire_class).sum())
        expected = len(stratum) * share
        low, high = (stats.chi2.ppf(0.025, 2 * observed) / 2 if observed else 0.0,
                     stats.chi2.ppf(0.975, 2 * (observed + 1)) / 2)
        rows.append(dict(cohort=cohort, cancer_type=cancer_type,
                         n_calls=len(stratum), observed=observed,
                         expected=round(expected, 1),
                         obs_over_exp=round(observed / expected, 3),
                         ci_low=round(low / expected, 3),
                         ci_high=round(high / expected, 3)))
    return pd.DataFrame(rows)


def heterogeneity(table):
    """Cochran's Q on the log ratios, with Poisson variances."""
    keep = table[table.observed > 0]
    if len(keep) < 2:
        return np.nan, np.nan, 0
    log_ratio = np.log(keep.obs_over_exp.to_numpy())
    weight = keep.observed.to_numpy().astype(float)   # 1 / var of log count
    pooled = float((weight * log_ratio).sum() / weight.sum())
    q = float((weight * (log_ratio - pooled) ** 2).sum())
    df = len(keep) - 1
    return q, float(stats.chi2.sf(q, df)), df


def leave_one_out(table):
    rows = []
    for cancer_type in table.cancer_type:
        rest = table[table.cancer_type != cancer_type]
        rows.append(dict(removed=cancer_type,
                         obs_over_exp=round(float(rest.observed.sum()
                                                  / rest.expected.sum()), 3)))
    return pd.DataFrame(rows).sort_values("obs_over_exp")


def main():
    universe = pd.read_parquet(cfg.RESULTS / "universe.parquet")
    universe["key"] = list(zip(universe.pos, universe.ref, universe.alt))
    with open(cfg.RESULTS / "spectra.pkl", "rb") as fh:
        spectra = pickle.load(fh)

    tables, summary = [], []
    for cohort in ("TCGA", "PCAWG", "MSK-IMPACT"):
        calls = load(cohort, universe)
        subset = eligible(universe, cohort)
        share = class_share(subset, spectra[cohort], "loading_site")
        table = by_cancer_type(calls, share, cohort, "loading_site")
        if table.empty:
            continue
        q, p, df = heterogeneity(table)
        loo = leave_one_out(table)
        pooled = float(table.observed.sum() / table.expected.sum())
        print(f"\n{cohort}: {len(table)} cancer types with at least "
              f"{MIN_CALLS} calls, pooled observed/expected {pooled:.3f}")
        print(f"  heterogeneity Cochran Q = {q:.1f} on {df} df, P = {p:.2f}")
        print(f"  types above 1: "
              f"{int((table.obs_over_exp > 1).sum())} of {len(table)}")
        print(f"  leave-one-type-out range {loo.obs_over_exp.min():.3f} to "
              f"{loo.obs_over_exp.max():.3f} "
              f"(lowest when removing {loo.removed.iloc[0]})")
        tables.append(table)
        summary.append(dict(cohort=cohort, n_types=len(table),
                            pooled=round(pooled, 3),
                            types_above_one=int((table.obs_over_exp > 1).sum()),
                            cochran_q=round(q, 2), q_df=df,
                            q_p_value=round(p, 3),
                            loo_min=float(loo.obs_over_exp.min()),
                            loo_max=float(loo.obs_over_exp.max())))

    types = pd.concat(tables, ignore_index=True)
    print("\nlargest and smallest strata")
    print(types.sort_values("n_calls", ascending=False)
          .head(12).to_string(index=False))

    # The PCAWG cluster-member depletion, by allele fraction.
    calls = load("PCAWG", universe)
    subset = eligible(universe, "PCAWG")
    rows = []
    for wire_class in cfg.WIRE_CLASSES:
        share = class_share(subset, spectra["PCAWG"], wire_class)
        for label, low, high in VAF_BANDS:
            band = calls[(calls.vaf >= low) & (calls.vaf < high)]
            if not len(band):
                continue
            observed = int((band.wire_class == wire_class).sum())
            expected = len(band) * share
            p = (stats.poisson.sf(observed - 1, expected) * 2
                 if observed > expected else stats.poisson.cdf(observed,
                                                               expected) * 2)
            rows.append(dict(wire_class=wire_class, vaf_band=label,
                             n_calls=len(band), observed=observed,
                             expected=round(expected, 1),
                             obs_over_exp=round(observed / expected, 3),
                             p_value=min(1.0, float(p))))
    bands = pd.DataFrame(rows)
    print("\nPCAWG by allele fraction")
    print(bands.pivot(index="vaf_band", columns="wire_class",
                      values="obs_over_exp")
          .reindex([b[0] for b in VAF_BANDS]).to_string())

    types.to_csv(cfg.RESULTS / "cancer_types.csv", index=False)
    pd.DataFrame(summary).to_csv(cfg.RESULTS / "cancer_type_summary.csv",
                                 index=False)
    bands.to_csv(cfg.RESULTS / "pcawg_vaf_bands.csv", index=False)
    print(f"\nwrote {cfg.RESULTS / 'cancer_types.csv'}")


if __name__ == "__main__":
    main()
```
