## Supplementary code for "Cancers relax selection on charge at the mitochondrial complex I quinone cavity": 17_jackknife.py

```
"""Leave-one-gene-out: is the result carried by one or two subunits?

The enrichment is strongest in ND1 and ND5, which raises the question of
whether those two subunits produce it. Each ND gene is removed in turn and the
selection analysis repeated on the remaining six, and each gene is also scored
on its own.
"""
import pickle
import sys
from pathlib import Path

import numpy as np
import pandas as pd

sys.path.insert(0, str(Path(__file__).resolve().parents[1]))
import config as cfg
from src import variants


def oe(subset, spectrum, column, wire_class):
    mask = (subset.wire_class == wire_class).to_numpy()
    observed, expected, ratio, p = variants.observed_expected(
        subset, spectrum, column, mask)
    return observed, expected, ratio, p


def main():
    universe = pd.read_parquet(cfg.RESULTS / "universe.parquet")
    with open(cfg.RESULTS / "spectra.pkl", "rb") as fh:
        spectra = pickle.load(fh)

    base = universe[(universe.kind == "missense") & universe.identical]
    rows = []
    for cohort in cfg.COHORTS:
        subset = base if cohort == "germline" else base[~base.polymorphism]
        column = f"n_{cohort}_missense"
        for wire_class in ("loading_site", "cut_vertex"):
            full = oe(subset, spectra[cohort], column, wire_class)
            rows.append(dict(cohort=cohort, wire_class=wire_class,
                             analysis="all seven genes", gene="-",
                             observed=full[0], expected=round(full[1], 1),
                             obs_over_exp=round(full[2], 3), p_value=full[3]))
            for gene in cfg.SUBUNITS:
                without = subset[subset.subunit != gene]
                result = oe(without, spectra[cohort], column, wire_class)
                rows.append(dict(cohort=cohort, wire_class=wire_class,
                                 analysis="gene removed", gene=gene,
                                 observed=result[0],
                                 expected=round(result[1], 1),
                                 obs_over_exp=round(result[2], 3),
                                 p_value=result[3]))
                only = subset[subset.subunit == gene]
                result = oe(only, spectra[cohort], column, wire_class)
                rows.append(dict(cohort=cohort, wire_class=wire_class,
                                 analysis="gene alone", gene=gene,
                                 observed=result[0],
                                 expected=round(result[1], 1),
                                 obs_over_exp=round(result[2], 3)
                                 if np.isfinite(result[2]) else np.nan,
                                 p_value=result[3]))
    table = pd.DataFrame(rows)

    for wire_class in ("loading_site", "cut_vertex"):
        d = table[(table.wire_class == wire_class)
                  & (table.analysis != "gene alone")]
        print(f"\n{wire_class}: observed/expected with each gene removed")
        print(d.pivot(index="gene", columns="cohort", values="obs_over_exp")
              .reindex(["-"] + list(cfg.SUBUNITS)).to_string())

    print("\nloading sites, each gene scored on its own")
    alone = table[(table.wire_class == "loading_site")
                  & (table.analysis == "gene alone")]
    print(alone.pivot(index="gene", columns="cohort",
                      values="obs_over_exp").reindex(
                          list(cfg.SUBUNITS)).to_string())

    removed = table[(table.wire_class == "loading_site")
                    & (table.analysis == "gene removed")]
    print("\nrange over the seven jackknife replicates, loading sites")
    for cohort in cfg.COHORTS:
        d = removed[removed.cohort == cohort]
        worst = d.loc[d.obs_over_exp.idxmin()]
        print(f"  {cohort:11s} {d.obs_over_exp.min():.3f} to "
              f"{d.obs_over_exp.max():.3f} "
              f"(lowest without {worst.gene}, P = {worst.p_value:.3g})")

    table.to_csv(cfg.RESULTS / "jackknife.csv", index=False)
    print(f"\nwrote {cfg.RESULTS / 'jackknife.csv'}")


if __name__ == "__main__":
    main()
```
