## Supplementary code for "Cancers relax selection on charge at the mitochondrial complex I quinone cavity": 18_axis_sensitivity.py

```
"""How much do the two free parameters of axis A matter?

Axis A is a screened Coulomb shift with an effective dielectric constant and a
floor on the donor-site distance, both chosen rather than derived. The
dielectric scales every score by the same factor, so it cannot change a
correlation or a ranking, but it does move scores across an absolute threshold;
the distance floor changes the ordering of nearby residues. Both are swept
here, and the variant-level contrasts are repeated with a quantile threshold,
which no choice of dielectric can affect.
"""
import pickle
import sys
from pathlib import Path

import numpy as np
import pandas as pd

sys.path.insert(0, str(Path(__file__).resolve().parents[1]))
import config as cfg
from src import axes

DIELECTRICS = (4.0, 8.0, 20.0)
FLOORS = (2.0, 3.0, 5.0)
QUANTILE = 0.75


def rebuild(built, dielectric, floor):
    cfg.DIELECTRIC, cfg.R_FLOOR = dielectric, floor
    nodes = [(k, rn, an, np.asarray(xyz)) for k, rn, an, xyz in built["nodes"]]
    metrics = axes.wire_metrics(nodes, built["graph"], built["classes"])
    loading_xyz = axes.loading_site_geometry(built["classes"], built["geometry"])
    major = {i for k in built["major"] for i in built["comps"][k]}
    waters = np.asarray([nodes[i][3] for i in sorted(major)
                         if nodes[i][1] == "HOH"])
    return axes.score_substitutions(built["geometry"], metrics, loading_xyz,
                                    waters)


def main():
    with open(cfg.RESULTS / "wire.pkl", "rb") as fh:
        built = pickle.load(fh)
    universe = pd.read_parquet(cfg.RESULTS / "universe.parquet")
    universe["key"] = list(zip(universe.pos, universe.ref, universe.alt))
    mapping = pd.read_csv(cfg.RESULTS / "mouse_human_map.csv")
    mapping = mapping[mapping.transferable & mapping.human_pos.notna()]
    to_human = {(r.subunit, int(r.mouse_pos)): int(r.human_pos)
                for r in mapping.itertuples()}

    missense = universe[(universe.kind == "missense") & universe.identical]
    common = missense[missense.polymorphism]
    default_dielectric, default_floor = cfg.DIELECTRIC, cfg.R_FLOOR

    rows = []
    for dielectric in DIELECTRICS:
        for floor in FLOORS:
            scores = rebuild(built, dielectric, floor)
            scores["human_pos"] = [to_human.get((s, p)) for s, p in
                                   zip(scores.subunit, scores.pos)]
            scores = scores[scores.human_pos.notna()]
            lookup = {(r.subunit, int(r.human_pos), r.mut_aa): r.axis_a_charge
                      for r in scores.itertuples()}

            def values(frame):
                out = [lookup.get((s, int(p), m)) for s, p, m in
                       zip(frame.subunit, frame.aa_pos, frame.mut_aa)]
                return np.array([v for v in out if v is not None], float)

            r = float(np.corrcoef(scores.axis_a_charge,
                                  scores.axis_b_connectivity)[0, 1])
            all_values, common_values = values(missense), values(common)
            cut = float(np.quantile(all_values, QUANTILE))
            rows.append(dict(
                dielectric=dielectric, r_floor=floor,
                pearson_r=round(r, 3),
                mean_axis_a=round(float(all_values.mean()), 4),
                pct_all_above_fixed=round(
                    100 * float((all_values >= cfg.AXIS_A_THRESHOLD).mean()), 1),
                pct_common_above_fixed=round(
                    100 * float((common_values >= cfg.AXIS_A_THRESHOLD).mean()), 1),
                quantile_cut=round(cut, 4),
                pct_common_above_quantile=round(
                    100 * float((common_values >= cut).mean()), 1)))
            print(f"  dielectric {dielectric:4.1f}, floor {floor:.1f} A: "
                  f"r = {r:.3f}, common germline above the fixed threshold "
                  f"{rows[-1]['pct_common_above_fixed']:.1f}% against "
                  f"{rows[-1]['pct_all_above_fixed']:.1f}%")

    cfg.DIELECTRIC, cfg.R_FLOOR = default_dielectric, default_floor
    table = pd.DataFrame(rows)
    print("\ncorrelation between the axes is unchanged by the dielectric "
          "(it scales axis A) and moves only with the distance floor:")
    print(table.pivot(index="r_floor", columns="dielectric",
                      values="pearson_r").to_string())
    print(f"\nthe depletion of loading-site charge change among common germline "
          f"variants, at a {int(QUANTILE * 100)}th-percentile threshold:")
    print(table.pivot(index="r_floor", columns="dielectric",
                      values="pct_common_above_quantile").to_string())
    print(f"(the same statistic for all possible substitutions is "
          f"{100 * (1 - QUANTILE):.0f}% by construction)")

    table.to_csv(cfg.RESULTS / "axis_a_sensitivity.csv", index=False)
    print(f"\nwrote {cfg.RESULTS / 'axis_a_sensitivity.csv'}")


if __name__ == "__main__":
    main()
```
