## Supplementary code for "Cancers relax selection on charge at the mitochondrial complex I quinone cavity": 19_multiple_testing.py

```
"""One multiple-testing table for the whole paper.

Each family of tests is corrected on its own by Benjamini-Hochberg, and the
whole set is corrected together as well, so that a reader can see at once which
claims survive which correction.
"""
import sys
from pathlib import Path

import pandas as pd

sys.path.insert(0, str(Path(__file__).resolve().parents[1]))
import config as cfg
from src import variants

SCORED = ("loading_site", "cut_vertex", "cluster_member")


def collect():
    rows = []

    table = pd.read_csv(cfg.RESULTS / "oe_missense.csv")
    for row in table[table.wire_class.isin(SCORED)].itertuples():
        rows.append(("Missense selection", f"{row.cohort}, {row.wire_class}",
                     row.obs_over_exp, row.p_value))

    table = pd.read_csv(cfg.RESULTS / "oe_synonymous.csv")
    for row in table[table.wire_class.isin(SCORED)].itertuples():
        rows.append(("Synonymous calibration",
                     f"{row.cohort}, {row.wire_class}",
                     row.obs_over_exp, row.p_value))

    table = pd.read_csv(cfg.RESULTS / "interaction.csv")
    table = table[table.model.str.startswith("unadjusted, all")]
    for row in table.itertuples():
        rows.append(("Tissue-type interaction", row.wire_class,
                     row.interaction_ratio, row.p_value))

    table = pd.read_csv(cfg.RESULTS / "recurrence.csv")
    for row in table[table.wire_class.isin(SCORED)].itertuples():
        rows.append(("Recurrence", f"{row.cohort}, {row.wire_class}",
                     round(row.recurrent_sites / row.expected_recurrent, 3)
                     if row.expected_recurrent else None, row.p_value))

    table = pd.read_csv(cfg.RESULTS / "haplogroup_enrichment.csv")
    for row in table[table.wire_class.isin(SCORED)].itertuples():
        rows.append((f"Phylogeny, {row.kind}", row.wire_class,
                     row.obs_over_exp, row.p_value))

    path = cfg.RESULTS / "disease_enrichment.csv"
    if path.exists():
        for row in pd.read_csv(path).itertuples():
            if row.wire_class in SCORED:
                rows.append(("Disease variants", f"{row.variant_set}, "
                             f"{row.wire_class}", row.obs_over_exp,
                             row.p_value))

    path = cfg.RESULTS / "nuclear_signatures.csv"
    if path.exists():
        for row in pd.read_csv(path).itertuples():
            rows.append(("Nuclear oxidative readouts",
                         f"{row.wire_class}, {row.signature}",
                         row.cmh_odds_ratio, row.cmh_p))

    return pd.DataFrame(rows, columns=["family", "test", "estimate",
                                       "p_value"])


def main():
    table = collect()
    table = table[table.p_value.notna()].reset_index(drop=True)

    table["q_within_family"] = (
        table.groupby("family").p_value
        .transform(lambda s: variants.benjamini_hochberg(s.to_numpy())))
    table["q_overall"] = variants.benjamini_hochberg(table.p_value.to_numpy())
    table = table.sort_values(["family", "p_value"])

    print(f"{len(table)} tests in {table.family.nunique()} families\n")
    print(table.to_string(index=False, float_format=lambda v: f"{v:.4g}"))

    print("\nsurviving a 5% false-discovery rate")
    for level, column in [("within its family", "q_within_family"),
                          ("over the whole paper", "q_overall")]:
        passing = table[table[column] < 0.05]
        print(f"\n  {level}: {len(passing)} of {len(table)}")
        for row in passing.itertuples():
            print(f"    {row.family:26s} {row.test:36s} "
                  f"{row.estimate}  q = {getattr(row, column):.4g}")

    table.to_csv(cfg.RESULTS / "multiple_testing.csv", index=False)
    print(f"\nwrote {cfg.RESULTS / 'multiple_testing.csv'}")


if __name__ == "__main__":
    main()
```
