## Supplementary code for "Cancers relax selection on charge at the mitochondrial complex I quinone cavity": 20_figure_s8.py

```
"""Supplementary Figure 8: coverage, cancer type, subunit and parameter checks."""
import sys
from pathlib import Path

import numpy as np
import pandas as pd
import matplotlib.pyplot as plt

sys.path.insert(0, str(Path(__file__).resolve().parents[1]))
import config as cfg
from src import plotting
from src.plotting import CLASS_COLOURS, CLASS_LABELS

SCORED = ("loading_site", "cut_vertex", "cluster_member")


def main():
    plotting.use_style()
    coverage = pd.read_csv(cfg.RESULTS / "coverage_by_class.csv")
    adjusted = pd.read_csv(cfg.RESULTS / "coverage_adjusted.csv")
    types = pd.read_csv(cfg.RESULTS / "cancer_types.csv")
    jackknife = pd.read_csv(cfg.RESULTS / "jackknife.csv")
    sensitivity = pd.read_csv(cfg.RESULTS / "axis_a_sensitivity.csv")

    fig, axes_ = plt.subplots(2, 2, figsize=(11.5, 9.0))

    # a) the rate ratio before and after adjusting for coverage
    ax = axes_[0, 0]
    cohorts = ["TCGA", "MSK-IMPACT"]
    loading = adjusted[adjusted.wire_class == "loading_site"]
    for i, model in enumerate(["unadjusted", "coverage-adjusted"]):
        values = [loading[(loading.cohort == c) & (loading.model == model)]
                  .rate_ratio.iloc[0] for c in cohorts]
        bars = ax.bar(np.arange(len(cohorts)) + (i - 0.5) * 0.36, values,
                      width=0.36,
                      color=["#aab7b8", CLASS_COLOURS["loading_site"]][i],
                      label=model.capitalize())
        for bar, value in zip(bars, values):
            ax.text(bar.get_x() + bar.get_width() / 2, value + 0.03,
                    f"{value:.2f}", ha="center", fontsize=9)
    ax.axhline(1.0, color="black", lw=0.8, ls=":")
    ax.set_xticks(np.arange(len(cohorts)))
    ax.set_xticklabels(cohorts)
    ax.set_ylim(0, 1.85)
    ax.set_ylabel("Rate ratio, loading sites")
    ax.legend(fontsize=9, loc="upper left")
    deficits = []
    for cohort in cohorts:
        d = coverage[coverage.cohort == cohort].set_index("wire_class")
        deficits.append(100 * (1 - d.loc["loading_site", "mean_coverage"]
                               / d.loc["other", "mean_coverage"]))
    ax.set_title(f"Loading sites are covered {deficits[0]:.0f}% less in TCGA "
                 f"and {deficits[1]:.0f}% less in MSK-IMPACT", fontsize=10)
    plotting.panel_label(ax, "a")

    # b) cancer types
    ax = axes_[0, 1]
    types = types.sort_values(["cohort", "obs_over_exp"]).reset_index(drop=True)
    colours = {"TCGA": CLASS_COLOURS["loading_site"],
               "PCAWG": "#5dade2", "MSK-IMPACT": "#aab7b8"}
    y = np.arange(len(types))
    for cohort, colour in colours.items():
        mask = (types.cohort == cohort).to_numpy()
        ax.errorbar(types.obs_over_exp[mask], y[mask],
                    xerr=[types.obs_over_exp[mask] - types.ci_low[mask],
                          types.ci_high[mask] - types.obs_over_exp[mask]],
                    fmt="o", ms=4, lw=0.9, color=colour, label=cohort)
    ax.axvline(1.0, color="black", lw=0.8, ls=":")
    ax.set_yticks(y)
    ax.set_yticklabels([f"{t[:18]}" for t in types.cancer_type], fontsize=6)
    ax.set_xscale("log")
    ax.set_xticks([0.25, 0.5, 1, 2, 4])
    ax.set_xticklabels(["0.25", "0.5", "1", "2", "4"])
    ax.xaxis.set_minor_formatter(plt.NullFormatter())
    ax.set_xlim(0.15, 7)
    ax.set_xlabel("Observed / expected, loading sites")
    ax.legend(fontsize=8, loc="lower right")
    ax.spines["left"].set_visible(True)
    plotting.panel_label(ax, "b", dx=-0.30)

    # c) leave-one-gene-out
    ax = axes_[1, 0]
    removed = jackknife[(jackknife.wire_class == "loading_site")
                        & (jackknife.analysis == "gene removed")]
    full = jackknife[(jackknife.wire_class == "loading_site")
                     & (jackknife.analysis == "all seven genes")]
    for i, cohort in enumerate(cfg.COHORTS):
        d = removed[removed.cohort == cohort]
        ax.scatter([i] * len(d), d.obs_over_exp, s=26, zorder=3,
                   color="#aab7b8", linewidths=0)
        value = full[full.cohort == cohort].obs_over_exp.iloc[0]
        ax.scatter([i], [value], s=70, marker="_", lw=2.5, zorder=4,
                   color=CLASS_COLOURS["loading_site"])
        worst = d.loc[d.obs_over_exp.idxmin()]
        ax.annotate(f"−{worst.gene}", (i, worst.obs_over_exp),
                    textcoords="offset points", xytext=(9, -3), fontsize=8)
    ax.axhline(1.0, color="black", lw=0.8, ls=":")
    ax.set_xticks(np.arange(len(cfg.COHORTS)))
    ax.set_xticklabels(list(cfg.COHORTS), rotation=20, ha="right")
    ax.set_ylabel("Observed / expected, loading sites")
    ax.set_title("Each ND gene removed in turn", fontsize=11)
    plotting.panel_label(ax, "c")

    # d) axis A parameters
    ax = axes_[1, 1]
    for dielectric, marker in zip(sorted(sensitivity.dielectric.unique()),
                                  ("o", "s", "^")):
        d = sensitivity[sensitivity.dielectric == dielectric].sort_values(
            "r_floor")
        ax.plot(d.r_floor, d.pearson_r, marker + "-", ms=7, color="black",
                mfc=CLASS_COLOURS["loading_site"],
                label=f"dielectric {dielectric:g}", alpha=0.85)
    ax.axhline(cfg.AXIS_A_THRESHOLD * 0 + 0.0, color="black", lw=0.8, ls=":")
    ax.set_ylim(0, 0.35)
    ax.set_xlabel("Distance floor of axis A (Å)")
    ax.set_ylabel("Pearson $r$ between the two axes")
    ax.set_title("The three curves coincide: the dielectric scales\naxis A "
                 "and cannot change a correlation", fontsize=10)
    ax.legend(fontsize=9)
    plotting.panel_label(ax, "d")

    fig.tight_layout()
    path = plotting.save(fig, cfg.FIGURES, "SuppFig8")
    print("wrote", path, "and", path.with_suffix(".png"))


if __name__ == "__main__":
    main()
```
