## Supplementary code for "Cancers relax selection on charge at the mitochondrial complex I quinone cavity": 21_structural_context.py

```
"""Where the classified residues sit in the enzyme.

Three questions. Whether the graph recovers the chain of charged residues that
runs along the middle of the membrane arm, which is the arrangement the
structural literature calls the central hydrophilic axis. Whether the ND1
signal comes from residues lining the quinone cavity, where a charge change
would act on quinone chemistry rather than on proton transfer, or from
residues further away. And where the substitutions that recur in tumours
actually are.
"""
import pickle
import sys
from pathlib import Path

import numpy as np
import pandas as pd

sys.path.insert(0, str(Path(__file__).resolve().parents[1]))
import config as cfg
from src import variants, wire

ANTIPORTERS = ("ND2", "ND4", "ND5")
CHARGED = {"ASP", "GLU", "LYS", "ARG", "HIS"}
MIDPLANE_SLAB = 6.0        # A either side of the bilayer centre
QUINONE_LIGAND = "WYK"     # the ligand bound in the quinone cavity of 8OM1
QUINONE_NEAR = 15.0        # A from the ligand centroid


def quinone_centroid(model):
    for chain in model:
        for residue in chain:
            if residue.name == QUINONE_LIGAND:
                return np.mean([[a.pos.x, a.pos.y, a.pos.z] for a in residue],
                               axis=0)
    raise SystemExit(f"{QUINONE_LIGAND} not found in the model")


def main():
    with open(cfg.RESULTS / "wire.pkl", "rb") as fh:
        built = pickle.load(fh)
    classes = built["classes"]
    geometry = built["geometry"]
    centre = np.asarray(built["centre"])
    normal = np.asarray(built["normal"])
    in_plane = np.asarray(built["in_plane"])

    model = wire.load_model(cfg.MOUSE_CIF, cfg.MOUSE_CHAINS)
    quinone = quinone_centroid(model)
    print(f"quinone-cavity ligand {QUINONE_LIGAND} centroid located; "
          f"{np.dot(quinone - centre, normal):.1f} Å from the bilayer centre "
          f"along the normal")

    rows = []
    for key, geom in geometry.items():
        wire_class = classes.get(key, "other")
        centroid = np.asarray(geom["centroid"])
        rows.append(dict(subunit=key[0], mouse_pos=key[1],
                         residue=geom["residue"], wire_class=wire_class,
                         along_normal=float(np.dot(centroid - centre, normal)),
                         along_plane=float(np.dot(centroid - centre, in_plane)),
                         dist_to_quinone=float(np.linalg.norm(
                             centroid - quinone)),
                         burial=geom["burial"]))
    residues = pd.DataFrame(rows)
    residues["charged"] = residues.residue.isin(CHARGED)

    # --- the central axis --------------------------------------------------
    axis = residues[(residues.wire_class == "loading_site")
                    & residues.charged
                    & residues.subunit.isin(ANTIPORTERS)
                    & (residues.along_normal.abs() <= MIDPLANE_SLAB)]
    axis = axis.sort_values("along_plane")
    print(f"\ncharged loading sites within {MIDPLANE_SLAB:.0f} Å of the "
          f"bilayer centre in the antiporter subunits: {len(axis)}")
    spacing = np.diff(axis.along_plane.to_numpy())
    print(f"they span {axis.along_plane.max() - axis.along_plane.min():.0f} Å "
          f"along the membrane plane, with neighbour spacing "
          f"{spacing.min():.1f} to {spacing.max():.1f} Å "
          f"(median {np.median(spacing):.1f} Å)")
    print(axis[["subunit", "mouse_pos", "residue", "along_plane",
                "along_normal"]].to_string(index=False))

    # How much of the midplane charge in these subunits did the graph pick up?
    pool = residues[residues.charged & residues.subunit.isin(ANTIPORTERS)
                    & (residues.along_normal.abs() <= MIDPLANE_SLAB)]
    print(f"\nof {len(pool)} charged residues in that slab, "
          f"{int((pool.wire_class != 'other').sum())} are in a hydrated "
          f"cluster and {int((pool.wire_class == 'loading_site').sum())} are "
          f"classified as loading sites")

    mapping = pd.read_csv(cfg.RESULTS / "mouse_human_map.csv")
    mapping = mapping[mapping.transferable & mapping.human_pos.notna()]
    to_human = {(r.subunit, int(r.mouse_pos)): int(r.human_pos)
                for r in mapping.itertuples()}
    residues["human_pos"] = [to_human.get((s, p)) for s, p in
                             zip(residues.subunit, residues.mouse_pos)]

    # --- ND1: quinone cavity or proton path --------------------------------
    nd1 = residues[(residues.subunit == "ND1")
                   & (residues.wire_class == "loading_site")]
    near = nd1[nd1.dist_to_quinone <= QUINONE_NEAR]
    print(f"\nND1 loading sites: {len(nd1)}, of which {len(near)} lie within "
          f"{QUINONE_NEAR:.0f} Å of the quinone ligand")
    print(nd1.sort_values("dist_to_quinone")[
        ["mouse_pos", "residue", "dist_to_quinone", "along_normal"]]
        .head(10).to_string(index=False))

    universe = pd.read_parquet(cfg.RESULTS / "universe.parquet")
    with open(cfg.RESULTS / "spectra.pkl", "rb") as fh:
        spectra = pickle.load(fh)
    missense = universe[(universe.kind == "missense") & universe.identical]

    groups = {
        "ND1, within 15 Å of the quinone site":
            {(r.subunit, int(r.human_pos)) for r in near.itertuples()
             if r.human_pos == r.human_pos},
        "ND1, further from the quinone site":
            {(r.subunit, int(r.human_pos)) for r in
             nd1[nd1.dist_to_quinone > QUINONE_NEAR].itertuples()
             if r.human_pos == r.human_pos},
        "loading sites outside ND1":
            {(r.subunit, int(r.human_pos)) for r in
             residues[(residues.wire_class == "loading_site")
                      & (residues.subunit != "ND1")].itertuples()
             if r.human_pos == r.human_pos},
    }
    split = []
    for label, keys in groups.items():
        for cohort in cfg.COHORTS:
            subset = missense if cohort == "germline" else missense[
                ~missense.polymorphism]
            mask = np.array([(s, p) in keys for s, p in
                             zip(subset.subunit, subset.aa_pos)])
            observed, expected, ratio, p = variants.observed_expected(
                subset, spectra[cohort], f"n_{cohort}_missense", mask)
            split.append(dict(group=label, cohort=cohort,
                              n_substitutions=int(mask.sum()),
                              observed=observed,
                              expected=round(expected, 1),
                              obs_over_exp=round(ratio, 3)
                              if np.isfinite(ratio) else np.nan, p_value=p))
    # The single most recurrent substitution carries part of this; how much?
    top = (3380, "G", "A")
    for label, keys in [("ND1, within 15 Å of the quinone site",
                         groups["ND1, within 15 Å of the quinone site"])]:
        for cohort in ("TCGA", "PCAWG", "MSK-IMPACT"):
            subset = missense[~missense.polymorphism].copy()
            drop = ((subset.pos == top[0]) & (subset.ref == top[1])
                    & (subset.alt == top[2]))
            subset.loc[drop, f"n_{cohort}_missense"] = 0
            mask = np.array([(s_, p_) in keys for s_, p_ in
                             zip(subset.subunit, subset.aa_pos)])
            observed, expected, ratio, p = variants.observed_expected(
                subset, spectra[cohort], f"n_{cohort}_missense", mask)
            split.append(dict(group=label + ", excluding m.3380G>A",
                              cohort=cohort, n_substitutions=int(mask.sum()),
                              observed=observed, expected=round(expected, 1),
                              obs_over_exp=round(ratio, 3)
                              if np.isfinite(ratio) else np.nan, p_value=p))

    split = pd.DataFrame(split)
    print("\nenrichment split by position relative to the quinone cavity")
    print(split.pivot(index="group", columns="cohort",
                      values="obs_over_exp").to_string())
    print("\nwith P values")
    print(split.to_string(index=False))

    # --- where the recurrent substitutions are -----------------------------
    recurrent = pd.read_csv(cfg.RESULTS / "recurrent_variants.csv")
    protein = recurrent.protein.str.extract(r"(\w+) (\w)(\d+)(\w)")
    recurrent["subunit"] = protein[0]
    recurrent["human_pos"] = protein[2].astype(int)
    lookup = {(r.subunit, int(r.human_pos)):
              (r.mouse_pos, r.residue, r.dist_to_quinone, r.along_normal,
               r.wire_class)
              for r in residues[residues.human_pos.notna()].itertuples()}
    context = []
    for row in recurrent.itertuples():
        hit = lookup.get((row.subunit, row.human_pos))
        if hit is None:
            continue
        context.append(dict(variant=row.variant, protein=row.protein,
                            patients=row.patients, wire_class=hit[4],
                            module="antiporter" if row.subunit in ANTIPORTERS
                            else "E-channel arm",
                            dist_to_quinone=round(hit[2], 1),
                            along_normal=round(hit[3], 1),
                            at_midplane=abs(hit[3]) <= MIDPLANE_SLAB))
    context = pd.DataFrame(context)
    print(f"\nstructural context of the {len(context)} recurrent "
          f"substitutions")
    print(context.head(16).to_string(index=False))
    print(f"\nat the bilayer midplane: {int(context.at_midplane.sum())} of "
          f"{len(context)}; within {QUINONE_NEAR:.0f} Å of the quinone "
          f"ligand: {int((context.dist_to_quinone <= QUINONE_NEAR).sum())}")

    residues.to_csv(cfg.RESULTS / "residue_structural_context.csv", index=False)
    axis.to_csv(cfg.RESULTS / "central_axis.csv", index=False)
    split.to_csv(cfg.RESULTS / "quinone_split.csv", index=False)
    context.to_csv(cfg.RESULTS / "recurrent_context.csv", index=False)
    print(f"\nwrote {cfg.RESULTS / 'central_axis.csv'}")


if __name__ == "__main__":
    main()
```
