## Supplementary code for "Cancers relax selection on charge at the mitochondrial complex I quinone cavity": 22_figure_s9.py

```
"""Supplementary Figure 9: how far the quinone localisation can be pushed.

The threshold, the model and the subunit effects are varied in turn, and the
null that the connectivity claim rests on is given a bound rather than a P
value alone. The last panel steps away from the quinone end and shows the
chain of charged loading sites at the bilayer midplane.
"""
import sys
from pathlib import Path

import numpy as np
import pandas as pd
import matplotlib.pyplot as plt

sys.path.insert(0, str(Path(__file__).resolve().parents[1]))
import config as cfg
from src import plotting
from src.plotting import CLASS_COLOURS

COLOURS = {"germline": "#aab7b8", "TCGA": "#5dade2",
           "PCAWG": CLASS_COLOURS["loading_site"], "MSK-IMPACT": "#1b2631"}
MODELS = ["proximity alone", "plus conservation and structure",
          "plus subunit", "within ND1 alone"]
LABELS = ["Unadjusted", "Conservation\nand structure", "Plus subunit\neffects",
          "Within ND1\nalone"]
ANTIPORTERS = ("ND2", "ND4", "ND5")
PANEL_SIZE = 19


def panel_label(ax, letter, dx=-0.16, dy=1.02):
    ax.text(dx, dy, letter, transform=ax.transAxes, fontsize=PANEL_SIZE,
            fontweight="bold", color="black", va="bottom", ha="left")


def main():
    plotting.use_style()
    residues = pd.read_csv(cfg.RESULTS / "residue_structural_context.csv")
    axis = pd.read_csv(cfg.RESULTS / "central_axis.csv")
    models = pd.read_csv(cfg.RESULTS / "quinone_adjusted.csv")
    interaction = pd.read_csv(cfg.RESULTS / "quinone_interaction.csv")
    bounds = pd.read_csv(cfg.RESULTS / "cut_vertex_bounds.csv")

    fig, axes_ = plt.subplots(2, 2, figsize=(11.5, 8.6))

    # a) proximity as a continuous variable, through the model ladder
    ax = axes_[0, 0]
    offsets = np.linspace(-0.27, 0.27, len(cfg.COHORTS))
    for offset, cohort in zip(offsets, cfg.COHORTS):
        d = models[models.cohort == cohort].set_index("model").reindex(MODELS)
        x = np.arange(len(MODELS)) + offset
        ax.errorbar(x, d.rate_ratio_per_e_fold,
                    yerr=[d.rate_ratio_per_e_fold - d.ci_low,
                          d.ci_high - d.rate_ratio_per_e_fold],
                    fmt="o", ms=5, lw=1.1, capsize=2.5, color=COLOURS[cohort],
                    label=cohort)
    ax.axhline(1.0, color="black", lw=0.8, ls=":")
    ax.set_yscale("log")
    ax.set_xticks(np.arange(len(MODELS)))
    ax.set_xticklabels(LABELS, fontsize=8)
    ax.set_ylabel("Rate ratio per e-fold\ndecrease in distance")
    ax.yaxis.set_minor_formatter(plt.NullFormatter())
    ax.legend(fontsize=8, ncol=2)
    panel_label(ax, "a")

    # b) is the gradient tumour-specific
    ax = axes_[0, 1]
    y = np.arange(len(interaction))
    ax.errorbar(interaction.interaction, y,
                xerr=[interaction.interaction - interaction.ci_low,
                      interaction.ci_high - interaction.interaction],
                fmt="o", ms=7, lw=1.3, capsize=3,
                color=CLASS_COLOURS["loading_site"])
    for yi, row in zip(y, interaction.itertuples()):
        ax.annotate(f"P = {row.p_value:.3f}", (row.interaction, yi),
                    textcoords="offset points", xytext=(0, 12), fontsize=8,
                    ha="center")
    ax.axvline(1.0, color="black", lw=0.8, ls=":")
    ax.set_xscale("log")
    ax.set_yticks(y)
    ax.set_yticklabels([m.replace(", ", ",\n") for m in interaction.model],
                       fontsize=8)
    ax.set_ylim(-0.6, len(interaction) - 0.2)
    ax.set_xlabel("Tumour gradient relative to the germline")
    ax.xaxis.set_minor_formatter(plt.NullFormatter())
    ax.spines["left"].set_visible(True)
    panel_label(ax, "b", dx=-0.38)

    # c) what the cut-vertex null excludes
    ax = axes_[1, 0]
    x = np.arange(len(bounds))
    ax.errorbar(x, bounds.obs_over_exp,
                yerr=[bounds.obs_over_exp - bounds.ci_low,
                      bounds.ci_high - bounds.obs_over_exp],
                fmt="o", ms=7, lw=1.3, capsize=3, color="#5dade2",
                label="Cut vertices, observed")
    ax.scatter(x, bounds.detectable_at_80_percent, marker="_", s=320, lw=2.2,
               color=CLASS_COLOURS["loading_site"],
               label="Detectable at 80% power")
    ax.axhline(1.0, color="black", lw=0.8, ls=":")
    ax.set_xticks(x)
    ax.set_xticklabels(bounds.cohort, rotation=15, ha="right")
    ax.set_ylabel("Observed / expected")
    ax.set_ylim(0.4, 1.8)
    ax.legend(fontsize=8)
    ax.set_title("The null the connectivity claim rests on", fontsize=10)
    panel_label(ax, "c")

    # d) the chain of charged residues at the bilayer midplane
    ax = axes_[1, 1]
    slab = residues[(residues.along_normal.abs() <= 6.0)
                    & residues.subunit.isin(ANTIPORTERS)
                    & residues.residue.isin(["ASP", "GLU", "LYS", "ARG",
                                             "HIS"])]
    rest = slab[slab.wire_class != "loading_site"]
    ax.scatter(rest.along_plane, rest.along_normal, s=28, c="#d5d8dc",
               linewidths=0, label="Charged, not a loading site")
    ax.scatter(axis.along_plane, axis.along_normal, s=60,
               c=CLASS_COLOURS["loading_site"], linewidths=0,
               label="Charged loading site")
    ordered = axis.sort_values("along_plane").reset_index(drop=True)
    ax.plot(ordered.along_plane, ordered.along_normal, lw=1.0,
            color=CLASS_COLOURS["loading_site"], zorder=0)
    for i, row in ordered.iterrows():
        ax.annotate(f"{row.subunit} {row.residue.title()}{row.mouse_pos}",
                    (row.along_plane, row.along_normal),
                    textcoords="offset points",
                    xytext=(0, 11 if i % 2 == 0 else -17), fontsize=6.5,
                    ha="center")
    ax.set_ylim(-10, 12)
    ax.set_xlabel("Position along the membrane plane (Å)")
    ax.set_ylabel("Position along the membrane normal (Å)")
    ax.set_title("Charged residues at the bilayer midplane", fontsize=11)
    ax.legend(fontsize=8, loc="lower right")
    panel_label(ax, "d")

    fig.tight_layout()
    path = plotting.save(fig, cfg.FIGURES, "SuppFig9")
    print("wrote", path, "and", path.with_suffix(".png"))


if __name__ == "__main__":
    main()
```
