## Supplementary code for "Cancers relax selection on charge at the mitochondrial complex I quinone cavity": 23_somatic_support.py

```
"""Are the recurrent calls somatic, and are they at mutational hotspots?

A substitution recurring across patients at a high allele fraction can mean
selection, but it can also mean a germline variant called somatic where the
matched normal is thin, or a position that mutates often. Both calls carry
matched-normal depth and alt counts, and the release annotates homopolymer
context, so both alternatives can be checked directly.
"""
import sys
from pathlib import Path

import numpy as np
import pandas as pd

sys.path.insert(0, str(Path(__file__).resolve().parents[1]))
import config as cfg

COHORT_FILES = {"TCGA": "data_mutations_tcga.txt.gz",
                "PCAWG": "data_mutations_pcawg.txt.gz",
                "MSK-IMPACT": "data_mutations_impact.txt.gz"}
MIN_NORMAL_DEPTH = 20
CLONAL_VAF = 0.9


def numeric(frame, column):
    return (pd.to_numeric(frame[column], errors="coerce") if column in frame
            else pd.Series(np.nan, index=frame.index))


def main():
    universe = pd.read_parquet(cfg.RESULTS / "universe.parquet")
    universe["key"] = list(zip(universe.pos, universe.ref, universe.alt))
    wire_class = dict(zip(universe.key, universe.wire_class))
    usable = dict(zip(universe.key, universe.identical))
    polymorphic = dict(zip(universe.key, universe.polymorphism))

    recurrent = pd.read_csv(cfg.RESULTS / "recurrent_variants.csv")
    parsed = recurrent.variant.str.extract(r"m\.(\d+)([ACGT])>([ACGT])")
    recurrent_keys = set(zip(parsed[0].astype(int), parsed[1], parsed[2]))

    rows, per_variant = [], []
    for cohort, filename in COHORT_FILES.items():
        calls = pd.read_csv(cfg.MTDNA / "original_data" / filename, sep="\t",
                            low_memory=False)
        calls = calls[calls.Hugo_Symbol.isin(sorted(universe.gene.unique()))
                      & (calls.Variant_Classification == "Missense_Mutation")
                      ].copy()
        calls["key"] = list(zip(calls.Start_Position, calls.Reference_Allele,
                                calls.Tumor_Seq_Allele2))
        calls = calls[calls.key.map(usable).fillna(False)
                      & ~calls.key.map(polymorphic).fillna(False)]
        calls["wire_class"] = calls.key.map(wire_class)
        calls["normal_depth"] = numeric(calls, "n_depth")
        calls["normal_alt"] = numeric(calls, "n_alt_count")
        calls["vaf"] = numeric(calls, "TumorVAF")
        calls["recurrent"] = calls.key.isin(recurrent_keys)
        calls["homopolymer"] = (numeric(calls, "homopolymer_width") >= 4
                                if "homopolymer_width" in calls else False)

        deep = calls[calls.normal_depth >= MIN_NORMAL_DEPTH]
        for label, subset in [("all wire-class calls",
                               calls[calls.wire_class.isin(
                                   ("loading_site", "cut_vertex",
                                    "cluster_member"))]),
                              ("recurrent calls", calls[calls.recurrent])]:
            with_normal = subset[subset.normal_depth >= MIN_NORMAL_DEPTH]
            rows.append(dict(
                cohort=cohort, set=label, calls=len(subset),
                median_normal_depth=float(subset.normal_depth.median()),
                calls_with_deep_normal=len(with_normal),
                with_any_normal_alt=int((with_normal.normal_alt > 0).sum()),
                pct_clonal=round(100 * float((subset.vaf >= CLONAL_VAF).mean()),
                                 1) if len(subset) else np.nan,
                pct_in_homopolymer=round(
                    100 * float(subset.homopolymer.mean()), 1)
                if len(subset) else np.nan))

        for key, group in calls[calls.recurrent].groupby("key"):
            deep_group = group[group.normal_depth >= MIN_NORMAL_DEPTH]
            per_variant.append(dict(
                cohort=cohort,
                variant=f"m.{key[0]}{key[1]}>{key[2]}",
                wire_class=group.wire_class.iloc[0],
                patients=group.Tumor_Sample_Barcode.nunique(),
                median_vaf=round(float(group.vaf.median()), 3),
                pct_clonal=round(100 * float((group.vaf >= CLONAL_VAF).mean()),
                                 1),
                median_normal_depth=float(group.normal_depth.median()),
                deep_normals=len(deep_group),
                normals_with_alt=int((deep_group.normal_alt > 0).sum()),
                in_homopolymer=bool(group.homopolymer.any())))

    summary = pd.DataFrame(rows)
    print(f"matched-normal support (a normal is counted when it reaches "
          f"{MIN_NORMAL_DEPTH} reads)")
    print(summary.to_string(index=False))

    variants_table = pd.DataFrame(per_variant)
    top = variants_table[variants_table.variant == "m.3380G>A"]
    print("\nthe most recurrent substitution, m.3380G>A")
    print(top.to_string(index=False))

    print("\nrecurrent substitutions with any alt read in a deep normal:")
    flagged = variants_table[variants_table.normals_with_alt > 0]
    print("  none" if flagged.empty else flagged.to_string(index=False))

    print("\nrecurrent substitutions annotated in a homopolymer of four or "
          "more:")
    hp = variants_table[variants_table.in_homopolymer]
    print("  none" if hp.empty else
          hp[["cohort", "variant", "patients"]].to_string(index=False))

    summary.to_csv(cfg.RESULTS / "somatic_support.csv", index=False)
    variants_table.to_csv(cfg.RESULTS / "recurrent_support.csv", index=False)
    print(f"\nwrote {cfg.RESULTS / 'somatic_support.csv'}")


if __name__ == "__main__":
    main()
```
