## Supplementary code for "Cancers relax selection on charge at the mitochondrial complex I quinone cavity": 24_quinone_robustness.py

```
"""Does the quinone result depend on the threshold, the covariates, or chance?

Four checks on the localisation. The 15 A radius is swept. The radius is then
removed altogether and the question asked as a distance: are the tumour calls
closer to the quinone cavity than the mutational null puts them? The distance
is entered into a regression alongside conservation and the structural
covariates, since residues near the cavity may simply be more conserved. And
the cut-vertex null, which carries half the claim, is given the effect size it
excludes.
"""
import pickle
import sys
from pathlib import Path

import numpy as np
import pandas as pd
import statsmodels.api as sm
import statsmodels.formula.api as smf
from scipy import stats

sys.path.insert(0, str(Path(__file__).resolve().parents[1]))
import config as cfg
from src import variants

RADII = (10.0, 15.0, 20.0, 25.0)
TUMOURS = ("TCGA", "PCAWG", "MSK-IMPACT")


def attach_distance(universe):
    context = pd.read_csv(cfg.RESULTS / "residue_structural_context.csv")
    context = context[context.human_pos.notna()]
    distance = {(r.subunit, int(r.human_pos)): r.dist_to_quinone
                for r in context.itertuples()}
    universe = universe.copy()
    universe["dist_to_quinone"] = [distance.get((s, p)) for s, p in
                                   zip(universe.subunit, universe.aa_pos)]
    return universe


def sweep(universe, spectra):
    rows = []
    loading = universe[(universe.kind == "missense") & universe.identical
                       & (universe.wire_class == "loading_site")
                       & universe.dist_to_quinone.notna()]
    for radius in RADII:
        inside = loading[loading.dist_to_quinone <= radius]
        outside = loading[loading.dist_to_quinone > radius]
        for label, group in (("within radius", inside),
                             ("beyond radius", outside)):
            for cohort in cfg.COHORTS:
                subset = universe[(universe.kind == "missense")
                                  & universe.identical]
                if cohort != "germline":
                    subset = subset[~subset.polymorphism]
                keys = set(zip(group.pos, group.ref, group.alt))
                mask = np.array([k in keys for k in
                                 zip(subset.pos, subset.ref, subset.alt)])
                observed, expected, ratio, p = variants.observed_expected(
                    subset, spectra[cohort], f"n_{cohort}_missense", mask)
                rows.append(dict(radius=radius, group=label, cohort=cohort,
                                 n_substitutions=int(mask.sum()),
                                 observed=observed,
                                 expected=round(expected, 1),
                                 obs_over_exp=round(ratio, 3)
                                 if np.isfinite(ratio) else np.nan,
                                 p_value=p))
    return pd.DataFrame(rows)


def distance_test(universe, spectra, rng):
    """No radius: are the observed calls closer to the cavity than expected?"""
    rows = []
    loading = universe[(universe.kind == "missense") & universe.identical
                       & (universe.wire_class == "loading_site")
                       & universe.dist_to_quinone.notna()]
    for cohort in cfg.COHORTS:
        subset = loading if cohort == "germline" else loading[
            ~loading.polymorphism]
        counts = subset[f"n_{cohort}_missense"].to_numpy()
        total = int(counts.sum())
        if total < 10:
            continue
        distance = subset.dist_to_quinone.to_numpy()
        observed = float((counts * distance).sum() / total)

        weights = subset.channel.map(spectra[cohort])
        weights = np.asarray((weights.fillna(weights.min())
                              / weights.sum()).to_numpy(), dtype=float)
        weights = weights / (weights.sum() * (1.0 + 1e-9))
        null = np.empty(cfg.N_PERMUTATIONS)
        chunk = 250
        filled = 0
        while filled < cfg.N_PERMUTATIONS:
            size = min(chunk, cfg.N_PERMUTATIONS - filled)
            draws = rng.multinomial(total, weights, size=size)
            null[filled:filled + size] = draws @ distance / total
            filled += size
        p = float((null <= observed).mean())
        rows.append(dict(cohort=cohort, calls=total,
                         mean_distance=round(observed, 2),
                         expected_distance=round(float(null.mean()), 2),
                         difference=round(observed - float(null.mean()), 2),
                         p_value=max(p, 1 / cfg.N_PERMUTATIONS)))
    return pd.DataFrame(rows)


def adjusted(universe, spectra):
    """Does proximity survive conservation and the structural covariates?"""
    rows = []
    loading = universe[(universe.kind == "missense") & universe.identical
                       & (universe.wire_class == "loading_site")
                       & universe.dist_to_quinone.notna()
                       & universe.phastcons.notna()
                       & universe.burial.notna()].copy()
    loading["proximity"] = -np.log(np.maximum(loading.dist_to_quinone, 1.0))
    for cohort in cfg.COHORTS:
        subset = (loading if cohort == "germline"
                  else loading[~loading.polymorphism]).copy()
        subset["y"] = subset[f"n_{cohort}_missense"]
        weights = subset.channel.map(spectra[cohort])
        weights = weights.fillna(weights.min())
        subset["offset"] = np.log(weights / weights.mean())
        for label, formula in [
                ("proximity alone", "y ~ proximity"),
                ("plus conservation", "y ~ proximity + phastcons"),
                ("plus conservation and structure",
                 "y ~ proximity + phastcons + burial + delta_volume"
                 " + delta_hydropathy"),
                ("plus subunit",
                 "y ~ proximity + C(subunit) + phastcons + burial"
                 " + delta_volume + delta_hydropathy"),
                ("within ND1 alone",
                 "y ~ proximity + phastcons + burial + delta_volume"
                 " + delta_hydropathy")]:
            fitted = subset[subset.subunit == "ND1"] if (
                label == "within ND1 alone") else subset
            fit = smf.glm(formula, data=fitted, family=sm.families.Poisson(),
                          offset=fitted["offset"]).fit(scale="X2")
            low, high = np.exp(fit.conf_int().loc["proximity"])
            rows.append(dict(cohort=cohort, model=label,
                             n_substitutions=len(fitted),
                             rate_ratio_per_e_fold=round(
                                 float(np.exp(fit.params["proximity"])), 3),
                             ci_low=round(float(low), 3),
                             ci_high=round(float(high), 3),
                             p_value=float(fit.pvalues["proximity"])))
    return pd.DataFrame(rows)


def proximity_interaction(universe, spectra):
    """Is the distance gradient steeper in tumours than in the germline?"""
    loading = universe[(universe.kind == "missense") & universe.identical
                       & (universe.wire_class == "loading_site")
                       & universe.dist_to_quinone.notna()
                       & universe.phastcons.notna()
                       & universe.burial.notna()].copy()
    loading["proximity"] = -np.log(np.maximum(loading.dist_to_quinone, 1.0))
    frames = []
    for cohort in cfg.COHORTS:
        subset = loading.copy()
        weights = subset.channel.map(spectra[cohort])
        weights = weights.fillna(weights.min())
        counts = subset[f"n_{cohort}_missense"].to_numpy()
        expected = counts.sum() * (weights / weights.sum()).to_numpy()
        subset["y"] = counts
        subset["offset"] = np.log(np.maximum(expected, 1e-12))
        subset["cohort"] = cohort
        subset["tumour"] = int(cohort != "germline")
        frames.append(subset)
    data = pd.concat(frames, ignore_index=True)
    data["proximity_tumour"] = data.proximity * data.tumour
    groups = (data.subunit + ":" + data.pos.astype(str) + data.alt).to_numpy()
    rows = []
    for label, formula in [
            ("subunit effects free to differ between arms",
             "y ~ proximity + C(cohort) + C(subunit) + C(subunit):tumour"
             " + proximity_tumour"),
            ("the same, adjusted",
             "y ~ proximity + C(cohort) + C(subunit) + C(subunit):tumour"
             " + proximity_tumour + phastcons + burial + delta_volume"
             " + delta_hydropathy")]:
        fit = smf.glm(formula, data=data, family=sm.families.Poisson(),
                      offset=data["offset"]).fit(
                          scale="X2", cov_type="cluster",
                          cov_kwds={"groups": groups})
        low, high = np.exp(fit.conf_int().loc["proximity_tumour"])
        rows.append(dict(model=label, n_rows=len(data),
                         germline_gradient=round(float(np.exp(
                             fit.params["proximity"])), 3),
                         interaction=round(float(np.exp(
                             fit.params["proximity_tumour"])), 3),
                         ci_low=round(float(low), 3),
                         ci_high=round(float(high), 3),
                         p_value=float(fit.pvalues["proximity_tumour"])))
    return pd.DataFrame(rows)


def cut_vertex_bound(universe, spectra):
    """What enrichment can the cut-vertex data exclude?"""
    rows = []
    subset = universe[(universe.kind == "missense") & universe.identical]
    for cohort in TUMOURS:
        d = subset[~subset.polymorphism]
        mask = (d.wire_class == "cut_vertex").to_numpy()
        observed, expected, ratio, _ = variants.observed_expected(
            d, spectra[cohort], f"n_{cohort}_missense", mask)
        low, high = variants.poisson_interval(observed, expected)
        # The smallest true ratio that would have been detected four times in
        # five, given this expected count.
        detectable = 1.0
        while detectable < 6.0:
            threshold = stats.poisson.ppf(0.975, expected)
            if stats.poisson.sf(threshold, expected * detectable) >= 0.8:
                break
            detectable += 0.01
        rows.append(dict(cohort=cohort, observed=observed,
                         expected=round(expected, 1),
                         obs_over_exp=round(ratio, 3),
                         ci_low=round(low, 3), ci_high=round(high, 3),
                         excluded_above=round(high, 2),
                         detectable_at_80_percent=round(detectable, 2)))
    return pd.DataFrame(rows)


def main():
    rng = np.random.default_rng(cfg.RANDOM_SEED)
    universe = pd.read_parquet(cfg.RESULTS / "universe_annotated.parquet")
    universe = attach_distance(universe)
    with open(cfg.RESULTS / "spectra.pkl", "rb") as fh:
        spectra = pickle.load(fh)

    radii = sweep(universe, spectra)
    print("loading-site enrichment by radius from the quinone ligand")
    print(radii[radii.group == "within radius"]
          .pivot(index="radius", columns="cohort", values="obs_over_exp")
          .to_string())
    print("\nand beyond that radius")
    print(radii[radii.group == "beyond radius"]
          .pivot(index="radius", columns="cohort", values="obs_over_exp")
          .to_string())

    distances = distance_test(universe, spectra, rng)
    print("\nmean distance to the quinone cavity of the observed calls, "
          "against the mutational null")
    print(distances.to_string(index=False))

    models = adjusted(universe, spectra)
    print("\nproximity to the cavity, adjusted "
          "(rate ratio per e-fold decrease in distance)")
    print(models.to_string(index=False))

    interaction = proximity_interaction(universe, spectra)
    print("\nis the distance gradient steeper in tumours than in the "
          "germline?")
    print(interaction.to_string(index=False))

    bounds = cut_vertex_bound(universe, spectra)
    print("\nwhat the cut-vertex data exclude")
    print(bounds.to_string(index=False))

    radii.to_csv(cfg.RESULTS / "quinone_radius_sweep.csv", index=False)
    distances.to_csv(cfg.RESULTS / "quinone_distance_test.csv", index=False)
    models.to_csv(cfg.RESULTS / "quinone_adjusted.csv", index=False)
    interaction.to_csv(cfg.RESULTS / "quinone_interaction.csv", index=False)
    bounds.to_csv(cfg.RESULTS / "cut_vertex_bounds.csv", index=False)
    print(f"\nwrote four tables to {cfg.RESULTS}")


if __name__ == "__main__":
    main()
```
