## Supplementary code for "Cancers relax selection on charge at the mitochondrial complex I quinone cavity": 25_figure_s10.py

```
"""Supplementary Figure 10: mitochondrial disease variants.

Pathogenic ND variants do not occupy the axis tumours leave alone, and the
two sets overlap strongly. What does separate at the variant level is the
population: substitutions common in the germline avoid loading-site charge
change specifically.
"""
import sys
from pathlib import Path

import numpy as np
import pandas as pd
import matplotlib.pyplot as plt

sys.path.insert(0, str(Path(__file__).resolve().parents[1]))
import config as cfg
from src import plotting
from src.plotting import CLASS_COLOURS, CLASS_LABELS
from src import variants

CLASSES = ("loading_site", "cut_vertex", "cluster_member", "other")
SET_COLOURS = {"MITOMAP confirmed": "#5b5b5b",
               "reported pathogenic": CLASS_COLOURS["loading_site"]}
GROUP_COLOURS = {"all possible": "#aab7b8", "tumour-observed": "#5dade2",
                 "reported pathogenic": CLASS_COLOURS["loading_site"],
                 "common germline": "#1b2631"}
CLIP = 5.4
PANEL_SIZE = 19


def panel_label(ax, letter, dx=-0.16, dy=1.02):
    ax.text(dx, dy, letter, transform=ax.transAxes, fontsize=PANEL_SIZE,
            fontweight="bold", color="black", va="bottom", ha="left")


def main():
    plotting.use_style()
    enrichment = pd.read_csv(cfg.RESULTS / "disease_enrichment.csv")
    groups = pd.read_csv(cfg.RESULTS / "axis_a_by_group.csv")
    overlap = pd.read_csv(cfg.RESULTS / "disease_overlap.csv")

    fig, axes_ = plt.subplots(1, 3, figsize=(13.0, 4.6))

    # a) are pathogenic variants enriched in any wire class
    ax = axes_[0]
    sets = list(enrichment.variant_set.unique())
    offsets = np.linspace(-0.18, 0.18, len(sets))
    for offset, variant_set in zip(offsets, sets):
        d = (enrichment[enrichment.variant_set == variant_set]
             .set_index("wire_class").reindex(CLASSES))
        low, high = [], []
        for row in d.itertuples():
            lo, hi = variants.poisson_interval(row.observed, row.expected)
            low.append(row.obs_over_exp - lo)
            high.append(min(hi, CLIP) - row.obs_over_exp)
        n = int(enrichment[enrichment.variant_set == variant_set].observed.sum())
        ax.errorbar(np.arange(len(CLASSES)) + offset, d.obs_over_exp,
                    yerr=[low, high], fmt="o", ms=6, lw=1.2, capsize=0,
                    color=SET_COLOURS[variant_set],
                    label=f"{variant_set} (n = {n})")
    ax.axhline(1.0, color="black", lw=0.8, ls="--")
    ax.set_xticks(np.arange(len(CLASSES)))
    ax.set_xticklabels([CLASS_LABELS[c].replace(" ", "\n") for c in CLASSES],
                       fontsize=9)
    ax.set_ylim(0, CLIP)
    ax.set_ylabel("observed / expected")
    ax.set_title("disease variants: underpowered,\nno distinct niche",
                 fontsize=10)
    ax.legend(fontsize=8.5)
    panel_label(ax, "a")

    # b) which groups avoid a charge change at a loading site
    ax = axes_[1]
    order = ["all possible", "tumour-observed", "reported pathogenic",
             "common germline"]
    d = groups.set_index("group").reindex(order)
    x = np.arange(len(order))
    ax.bar(x, d.pct_above_threshold, width=0.68,
           color=[GROUP_COLOURS[g] for g in order])
    for xi, (pct, n) in enumerate(zip(d.pct_above_threshold, d.n)):
        ax.text(xi, pct + 0.6, f"{pct:.1f}%", ha="center", fontsize=9)
        ax.text(xi, 0.6, f"n = {int(n):,}", ha="center", fontsize=8,
                color="white")
    ax.set_xticks(x)
    ax.set_xticklabels([g.replace(" ", "\n") for g in order], fontsize=9)
    ax.set_ylabel(f"% of substitutions shifting a\nloading site by "
                  f"≥ {cfg.AXIS_A_THRESHOLD} eV")
    ax.set_title("common germline variants avoid\nloading-site charge change",
                 fontsize=10)
    panel_label(ax, "b")

    # c) how far the disease and tumour sets overlap
    ax = axes_[2]
    x = np.arange(len(overlap))
    ax.bar(x, overlap.pct, width=0.62,
           color=[CLASS_COLOURS["loading_site"], "#aab7b8"])
    odds = float(overlap.odds_ratio.iloc[0])
    p = float(overlap.p_value.iloc[0])
    ax.text(0.97, 0.93, f"odds ratio {odds:.1f}\nP = {p:.0e}",
            transform=ax.transAxes, ha="right", va="top", fontsize=9)
    ax.set_xticks(x)
    ax.set_xticklabels([g.replace(" ", "\n") for g in overlap.group],
                       fontsize=9)
    ax.set_ylabel("% also observed in a tumour")
    ax.set_title("disease and tumour variants\noverlap strongly", fontsize=10)
    panel_label(ax, "c")

    fig.tight_layout()
    path = plotting.save(fig, cfg.FIGURES, "SuppFig10")
    print("wrote", path, "and", path.with_suffix(".png"))


if __name__ == "__main__":
    main()
```
