## Supplementary code for "Cancers relax selection on charge at the mitochondrial complex I quinone cavity": 26_main_figures.py

```
"""Figures 2 and 4.

Figure 2 sets the selection result against the mutational null, adjusts it,
breaks it down by subunit, and closes with the interaction that tests the
germline-to-tumour difference directly. Figure 4 places the tumour signal in
the enzyme: it sits at the quinone end of ND1 rather than along the proton
path.

Both read tables written by scripts 05, 06, 13, 21 and 24, so this script runs
after those rather than with the rest of the figures.
"""
import sys
from pathlib import Path

import numpy as np
import pandas as pd
import matplotlib.pyplot as plt

sys.path.insert(0, str(Path(__file__).resolve().parents[1]))
import config as cfg
from src import plotting
from src.plotting import CLASS_COLOURS

# Figure 2 separates the wire classes by colour rather than by marker, so it
# carries its own palette; the marker still names the cohort.
CLASS_PALETTE = {"loading_site": "#EB6834", "cut_vertex": "#2A78D6",
                 "cluster_member": "#1BAF7A", "other": "#9A9994"}
CLASS_STACKED = {"loading_site": "proton\nloading site",
                 "cut_vertex": "wire cut\nvertex",
                 "cluster_member": "hydrated\ncluster",
                 "other": "other ND\nresidues"}
MARKERS = {"germline": "o", "TCGA": "s", "PCAWG": "^", "MSK-IMPACT": "D"}
SCORED = ("loading_site", "cut_vertex", "cluster_member")

# Figure 4 keeps the cohort palette used for the quinone analyses.
COHORT_COLOURS = {"germline": "#aab7b8", "TCGA": "#5dade2",
                  "PCAWG": CLASS_COLOURS["loading_site"],
                  "MSK-IMPACT": "#1b2631"}
QUINONE_NEAR = 15.0
MIN_EXPECTED = 3.0
GROUPS = ["ND1, within 15 Å of the quinone site",
          "ND1, further from the quinone site",
          "loading sites outside ND1"]
GROUP_LABELS = ["ND1, within 15 Å\nof the quinone site",
                "ND1, further from\nthe quinone site",
                "Loading sites\noutside ND1"]

PANEL_SIZE = 19


def panel_label(ax, letter, dx=-0.16, dy=1.02):
    ax.text(dx, dy, letter, transform=ax.transAxes, fontsize=PANEL_SIZE,
            fontweight="bold", color="black", va="bottom", ha="left")


# --- Figure 2 -------------------------------------------------------------

def observed_expected(ax, missense):
    classes = list(cfg.WIRE_CLASSES)
    offsets = np.linspace(-0.26, 0.26, len(cfg.COHORTS))
    for offset, cohort in zip(offsets, cfg.COHORTS):
        for i, wire_class in enumerate(classes):
            row = missense[(missense.cohort == cohort)
                           & (missense.wire_class == wire_class)]
            if row.empty:
                continue
            row = row.iloc[0]
            colour = CLASS_PALETTE[wire_class]
            ax.errorbar(i + offset, row.obs_over_exp,
                        yerr=[[row.obs_over_exp - row.ci_low],
                              [row.ci_high - row.obs_over_exp]],
                        fmt=MARKERS[cohort], ms=8, lw=1.3, capsize=0,
                        color=colour, mec=colour, mew=1.6,
                        mfc="white" if cohort == "germline" else colour)
    for cohort in cfg.COHORTS:
        ax.plot([], [], MARKERS[cohort], color="black", ls="none", ms=8,
                mfc="white" if cohort == "germline" else "black", label=cohort)
    ax.axhline(1.0, color="black", lw=0.9, ls="--")
    ax.set_xticks(np.arange(len(classes)))
    ax.set_xticklabels([CLASS_STACKED[c] for c in classes], fontsize=9.5)
    ax.set_xlim(-0.6, len(classes) - 0.4)
    ax.set_yscale("log")
    ax.set_yticks([0.3, 0.5, 1.0, 1.5, 2.0, 2.5])
    ax.set_yticklabels(["0.3", "0.5", "1.0", "1.5", "2.0", "2.5"])
    ax.yaxis.set_minor_formatter(plt.NullFormatter())
    ax.set_ylim(0.26, 3.05)
    ax.set_ylabel("observed / expected")
    ax.set_title("germline purges the proton wire; "
                 "tumours enrich only its loading sites", fontsize=10.5, pad=26)
    ax.legend(loc="upper center", ncol=4, fontsize=9,
              bbox_to_anchor=(0.5, 1.04), handletextpad=0.3, columnspacing=1.4)
    panel_label(ax, "a", dx=-0.12)


def adjusted_rate_ratios(ax, ladder):
    d = ladder[ladder.model == "conservation + structure"]
    labels = {"loading_site": "loading site (charge)",
              "cut_vertex": "cut vertex (connectivity)"}
    cohorts = list(cfg.COHORTS)
    base = np.arange(len(cohorts))[::-1]
    for offset, wire_class in zip((0.17, -0.17), ("loading_site", "cut_vertex")):
        for y, cohort in zip(base, cohorts):
            row = d[(d.cohort == cohort) & (d.wire_class == wire_class)]
            if row.empty:
                continue
            row = row.iloc[0]
            ax.errorbar(row.rate_ratio, y + offset,
                        xerr=[[row.rate_ratio - row.ci_low],
                              [row.ci_high - row.rate_ratio]],
                        fmt="o", ms=7, lw=1.6, capsize=0,
                        color=CLASS_PALETTE[wire_class], zorder=3)
        ax.plot([], [], "o-", color=CLASS_PALETTE[wire_class], ms=7,
                label=labels[wire_class])
    ax.axvline(1.0, color="black", lw=0.9, ls="--")
    ax.set_xscale("log")
    ax.set_xticks([0.3, 0.5, 1, 2])
    ax.set_xticklabels(["0.3", "0.5", "1", "2"])
    ax.xaxis.set_minor_formatter(plt.NullFormatter())
    ax.set_yticks(base)
    ax.set_yticklabels(cohorts, fontsize=9.5)
    ax.set_ylim(-0.6, len(cohorts) - 0.4)
    ax.set_xlabel("rate ratio vs other ND residues\n"
                  "(adjusted for conservation, burial, volume, hydropathy)",
                  fontsize=9.5)
    ax.legend(loc="upper center", ncol=2, fontsize=9,
              bbox_to_anchor=(0.5, 1.12), handletextpad=0.3)
    ax.spines["left"].set_visible(True)
    panel_label(ax, "b", dx=-0.30)


def by_subunit(ax, subunits):
    d = (subunits[subunits.expected >= MIN_EXPECTED]
         .sort_values("subunit", ascending=False).reset_index(drop=True))
    y = np.arange(len(d))
    ax.errorbar(d.obs_over_exp, y,
                xerr=[d.obs_over_exp - d.ci_low, d.ci_high - d.obs_over_exp],
                fmt="o", ms=7, lw=1.6, capsize=0,
                color=CLASS_PALETTE["loading_site"], zorder=3)
    ax.axvline(1.0, color="black", lw=0.9, ls="--")
    ax.set_yticks(y)
    ax.set_yticklabels(d.subunit, fontsize=9.5)
    ax.set_ylim(-0.6, len(d) - 0.4)
    ax.set_xlim(0, 4.2)
    ax.set_xlabel("loading-site O/E in PCAWG", fontsize=9.5)
    ax.spines["left"].set_visible(True)
    ax.text(1.02, 1.02, "obs/exp", transform=ax.transAxes, fontsize=9,
            ha="left", va="bottom")
    trans = ax.get_yaxis_transform()
    for yi, row in zip(y, d.itertuples()):
        ax.text(1.02, yi, f"{int(row.observed)}/{row.expected:.0f}",
                transform=trans, fontsize=9, ha="left", va="center")
    panel_label(ax, "c", dx=-0.22)


def interaction_panel(ax, interaction):
    d = interaction.set_index("wire_class").reindex(SCORED)
    y = np.arange(len(SCORED))[::-1]
    for yi, wire_class in zip(y, SCORED):
        row = d.loc[wire_class]
        ax.errorbar(row.interaction_ratio, yi,
                    xerr=[[row.interaction_ratio - row.ci_low],
                          [row.ci_high - row.interaction_ratio]],
                    fmt="o", ms=9, lw=1.6, capsize=4,
                    color=CLASS_PALETTE[wire_class], zorder=3)
        p = row.p_value
        p_text = (f"P = {p:.2f}" if p >= 0.01
                  else f"P = {p:.1e}".replace("e-0", " × 10⁻").replace("e-", " × 10⁻"))
        ax.annotate(f"{row.interaction_ratio:.2f}", (row.interaction_ratio, yi),
                    textcoords="offset points", xytext=(0, 12), fontsize=9.5,
                    ha="center", fontweight="bold")
        ax.annotate(p_text, (row.interaction_ratio, yi),
                    textcoords="offset points", xytext=(0, -20), fontsize=8.5,
                    ha="center")
        ax.annotate(f"{row.germline_rate_ratio:.2f}", (1.04, yi),
                    xycoords=("axes fraction", "data"), fontsize=9,
                    ha="left", va="center", color="#444444",
                    annotation_clip=False)
    ax.axvline(1.0, color="black", lw=0.9, ls=":")
    ax.set_xscale("log")
    ax.set_xticks([1, 2, 5])
    ax.set_xticklabels(["1", "2", "5"])
    ax.xaxis.set_minor_formatter(plt.NullFormatter())
    ax.set_xlim(0.65, 7.0)
    ax.set_yticks(y)
    ax.set_yticklabels([CLASS_STACKED[c].capitalize() for c in SCORED],
                       fontsize=9)
    ax.set_ylim(-0.75, len(SCORED) - 0.25)
    ax.text(1.04, 1.02, "germline\nrate ratio", transform=ax.transAxes,
            fontsize=8.5, ha="left", va="bottom", color="#444444")
    ax.set_xlabel("tumour / germline rate ratio\n"
                  "(wire class × tissue type)", fontsize=9.5)
    ax.set_title("the constraint tumours relax\nis on charge, not connectivity",
                 fontsize=10)
    ax.spines["left"].set_visible(True)
    panel_label(ax, "d", dx=-0.20)


def figure2(missense, ladder, subunits, interaction):
    fig = plt.figure(figsize=(14.4, 9.8))
    grid = fig.add_gridspec(2, 3, height_ratios=[1.0, 0.92], hspace=0.52,
                            wspace=0.62)
    observed_expected(fig.add_subplot(grid[0, :]), missense)
    adjusted_rate_ratios(fig.add_subplot(grid[1, 0]), ladder)
    by_subunit(fig.add_subplot(grid[1, 1]), subunits)
    interaction_panel(fig.add_subplot(grid[1, 2]), interaction)
    return plotting.save(fig, cfg.FIGURES, "Figure2")


# --- Figure 4 -------------------------------------------------------------

def membrane_frame(ax, residues):
    other = residues[residues.wire_class == "other"]
    ax.scatter(other.along_plane, other.along_normal, s=3, c="#e5e8e8",
               linewidths=0)
    cluster = residues[residues.wire_class.isin(["cluster_member",
                                                 "cut_vertex"])]
    ax.scatter(cluster.along_plane, cluster.along_normal, s=9, c="#aab7b8",
               linewidths=0, label="Cluster member or cut vertex")
    loading = residues[residues.wire_class == "loading_site"]
    far = loading[loading.dist_to_quinone > QUINONE_NEAR]
    near = loading[loading.dist_to_quinone <= QUINONE_NEAR]
    ax.scatter(far.along_plane, far.along_normal, s=22, c="#5dade2",
               linewidths=0, label="Proton loading site")
    ax.scatter(near.along_plane, near.along_normal, s=46, marker="D",
               c=CLASS_COLOURS["loading_site"], linewidths=0,
               label=f"Loading site within {QUINONE_NEAR:.0f} Å of quinone")
    ax.axhline(0, color="black", lw=0.6, ls=":")
    ax.set_xlabel("Position along the membrane plane (Å)")
    ax.set_ylabel("Position along the membrane normal (Å)")
    ax.legend(fontsize=8, loc="lower left")
    ax.set_title("The tumour signal sits at one end of the arm", fontsize=11)
    panel_label(ax, "a", dx=-0.10)


def quinone_split(ax, split):
    offsets = np.linspace(-0.3, 0.3, len(cfg.COHORTS))
    for offset, cohort in zip(offsets, cfg.COHORTS):
        values = [split[(split.group == g) & (split.cohort == cohort)]
                  .obs_over_exp.iloc[0] for g in GROUPS]
        ax.bar(np.arange(len(GROUPS)) + offset, values, width=0.2,
               color=COHORT_COLOURS[cohort], label=cohort)
    ax.axhline(1.0, color="black", lw=0.8, ls=":")
    ax.set_xticks(np.arange(len(GROUPS)))
    ax.set_xticklabels(GROUP_LABELS, fontsize=9)
    ax.set_ylabel("Observed / expected")
    ax.legend(fontsize=8, ncol=2)
    panel_label(ax, "b")


def one_substitution(ax, split):
    cohorts = [c for c in cfg.COHORTS if c != "germline"]
    full = [split[(split.group == GROUPS[0]) & (split.cohort == c)]
            .obs_over_exp.iloc[0] for c in cohorts]
    dropped = [split[(split.group == GROUPS[0] + ", excluding m.3380G>A")
                     & (split.cohort == c)].obs_over_exp.iloc[0]
               for c in cohorts]
    x = np.arange(len(cohorts))
    ax.bar(x - 0.19, full, width=0.38, color=CLASS_COLOURS["loading_site"],
           label="All substitutions")
    ax.bar(x + 0.19, dropped, width=0.38, color="#aab7b8",
           label="Excluding m.3380G>A")
    for xi, (a, b) in enumerate(zip(full, dropped)):
        ax.text(xi - 0.19, a + 0.08, f"{a:.1f}", ha="center", fontsize=9)
        ax.text(xi + 0.19, b + 0.08, f"{b:.1f}", ha="center", fontsize=9)
    ax.axhline(1.0, color="black", lw=0.8, ls=":")
    ax.set_xticks(x)
    ax.set_xticklabels(cohorts, rotation=15, ha="right")
    ax.set_ylabel("Observed / expected,\nquinone-proximal ND1 loading sites")
    ax.legend(fontsize=9)
    panel_label(ax, "c")


def radius_sweep(ax, radii):
    for cohort in cfg.COHORTS:
        inside = radii[(radii.cohort == cohort)
                       & (radii.group == "within radius")].sort_values("radius")
        outside = radii[(radii.cohort == cohort)
                        & (radii.group == "beyond radius")].sort_values("radius")
        ax.plot(inside.radius, inside.obs_over_exp, "o-", ms=6,
                color=COHORT_COLOURS[cohort], label=cohort)
        ax.plot(outside.radius, outside.obs_over_exp, "o--", ms=4, lw=1.0,
                color=COHORT_COLOURS[cohort], alpha=0.55)
    ax.axhline(1.0, color="black", lw=0.8, ls=":")
    ax.set_yscale("log")
    ax.set_yticks([0.25, 0.5, 1, 2, 4, 6])
    ax.set_yticklabels(["0.25", "0.5", "1", "2", "4", "6"])
    ax.yaxis.set_minor_formatter(plt.NullFormatter())
    ax.set_ylim(0.22, 7.5)
    ax.set_xlabel("Radius from the quinone ligand (Å)")
    ax.set_ylabel("Observed / expected, loading sites")
    ax.set_title("Solid, within the radius; dashed, beyond it", fontsize=10)
    ax.legend(fontsize=8, ncol=2)
    panel_label(ax, "d")


def figure4(residues, split, radii):
    fig, axes_ = plt.subplots(2, 2, figsize=(11.5, 8.8))
    membrane_frame(axes_[0, 0], residues)
    quinone_split(axes_[0, 1], split)
    one_substitution(axes_[1, 0], split)
    radius_sweep(axes_[1, 1], radii)
    fig.tight_layout()
    return plotting.save(fig, cfg.FIGURES, "Figure4")


def main():
    plotting.use_style()
    read = lambda name: pd.read_csv(cfg.RESULTS / name)

    interaction = read("interaction.csv")
    primary = interaction[interaction.model == "unadjusted, all identical codons"]

    written = [
        figure2(read("oe_missense.csv"), read("rate_ratios.csv"),
                read("oe_by_subunit.csv"), primary),
        figure4(read("residue_structural_context.csv"),
                read("quinone_split.csv"), read("quinone_radius_sweep.csv")),
    ]
    for path in written:
        print(f"wrote {path} and {path.with_suffix('.png')}")


if __name__ == "__main__":
    main()
```
