## Supplementary code for "Cancers relax selection on charge at the mitochondrial complex I quinone cavity": axes.py

```
"""The two axes: proton-loading electrostatics and proton-wire connectivity."""
from __future__ import annotations

import networkx as nx
import numpy as np
import pandas as pd
from scipy.spatial import cKDTree

import config as cfg

AMINO_ACIDS = ["ALA", "ARG", "ASN", "ASP", "CYS", "GLN", "GLU", "GLY", "HIS",
               "ILE", "LEU", "LYS", "MET", "PHE", "PRO", "SER", "THR", "TRP",
               "TYR", "VAL"]
ONE_LETTER = dict(zip(AMINO_ACIDS, "ARNDCQEGHILKMFPSTWYV"))
# Side-chain volumes (A^3) and Kyte-Doolittle hydropathy.
VOLUME = dict(zip(AMINO_ACIDS, [88.6, 173.4, 114.1, 111.1, 108.5, 143.8, 138.4,
                                60.1, 153.2, 166.7, 166.7, 168.6, 162.9, 189.9,
                                112.7, 89.0, 116.1, 227.8, 193.6, 140.0]))
HYDROPATHY = dict(zip(AMINO_ACIDS, [1.8, -4.5, -3.5, -3.5, 2.5, -3.5, -3.5, -0.4,
                                    -3.2, 4.5, 3.8, -3.9, 1.9, 2.8, -1.6, -0.8,
                                    -0.7, -0.9, -1.3, 4.2]))


def loading_site_geometry(classes, geometry):
    """Centroids of the proton loading sites."""
    return np.asarray([geometry[k]["centroid"] for k, c in classes.items()
                       if c == "loading_site" and k in geometry])


def wire_metrics(nodes, graph, classes):
    """Per-residue betweenness, articulation status and fragmentation.

    Fragmentation is the number of extra components a residue's removal creates
    in its own cluster, which is what makes a cut vertex costly.
    """
    keys = [n[0] for n in nodes]
    comps = [c for c in nx.connected_components(graph) if len(c) >= cfg.MIN_CLUSTER]
    cluster_of = {i: k for k, c in enumerate(comps) for i in c}

    betweenness = {}
    for c in comps:
        if len(c) < 6:
            continue
        for i, b in nx.betweenness_centrality(graph.subgraph(c),
                                              weight="weight",
                                              normalized=True).items():
            betweenness[i] = b

    articulation = set()
    for c in comps:
        articulation |= set(nx.articulation_points(graph.subgraph(c)))

    nodes_of = {}
    for i, key in enumerate(keys):
        if isinstance(key, tuple) and key[0] in cfg.SUBUNITS:
            nodes_of.setdefault(key, []).append(i)

    out = {}
    for key, idx in nodes_of.items():
        clusters = {cluster_of[i] for i in idx if i in cluster_of}
        fragments = 0
        for k in clusters:
            sub = graph.subgraph(comps[k]).copy()
            sub.remove_nodes_from([i for i in idx if i in comps[k]])
            if sub.number_of_nodes():
                fragments = max(fragments, nx.number_connected_components(sub) - 1)
        out[key] = {
            "in_cluster": bool(clusters),
            "articulation": any(i in articulation for i in idx),
            "betweenness": max((betweenness.get(i, 0.0) for i in idx), default=0.0),
            "fragments": fragments,
            "wire_class": classes.get(key, "other"),
        }
    return out


def score_substitutions(geometry, metrics, loading_xyz, cluster_water_xyz):
    """Score every possible missense substitution on both axes.

    Axis A, charge: the screened Coulomb shift the substitution's net charge
    change imposes on a proton loading site, taken as the largest absolute
    shift over all sites.

    Axis B, connectivity: loss of a Grotthuss-competent side chain that is part
    of a hydrated cluster, weighted by how much of the cluster it holds
    together, plus a steric term for substitutions that enlarge a side chain
    lining cluster waters.
    """
    water_tree = cKDTree(cluster_water_xyz) if len(cluster_water_xyz) else None
    rows = []
    for key, geom in geometry.items():
        wt = geom["residue"]
        if wt not in AMINO_ACIDS:
            continue
        centroid = geom["centroid"]
        m = metrics.get(key, {"in_cluster": False, "articulation": False,
                              "betweenness": 0.0, "fragments": 0,
                              "wire_class": "other"})
        if len(loading_xyz):
            distances = np.maximum(np.linalg.norm(loading_xyz - centroid, axis=1),
                                   cfg.R_FLOOR)
        else:
            distances = np.array([1e3])
        n_waters = (len(water_tree.query_ball_point(centroid, cfg.STERIC_WATER_SHELL))
                    if water_tree is not None else 0)
        wt_charge = cfg.CHARGE.get(wt, 0.0)

        for mut in AMINO_ACIDS:
            if mut == wt:
                continue
            dq = cfg.CHARGE.get(mut, 0.0) - wt_charge
            axis_a = (float(np.max(np.abs(14.4 * dq / (cfg.DIELECTRIC * distances))))
                      if dq else 0.0)

            axis_b = 0.0
            loses_wire = (m["in_cluster"] and wt in cfg.WIRE_ATOMS
                          and mut not in cfg.WIRE_ATOMS)
            if loses_wire:
                axis_b += (1.0 + m["fragments"] + 4.0 * m["betweenness"]
                           + (1.0 if m["articulation"] else 0.0))
            dv = VOLUME[mut] - VOLUME[wt]
            steric = 0.0
            if dv > cfg.STERIC_VOLUME_THRESHOLD and n_waters:
                steric = 0.02 * (dv - cfg.STERIC_VOLUME_THRESHOLD) / 10.0 * n_waters
            axis_b += steric

            rows.append((key[0], key[1], ONE_LETTER[wt], ONE_LETTER[mut], dq,
                         axis_a, axis_b, axis_b - steric, m["wire_class"],
                         m["in_cluster"], m["articulation"], m["fragments"],
                         float(distances.min()), n_waters, dv,
                         HYDROPATHY[mut] - HYDROPATHY[wt], geom["burial"]))

    return pd.DataFrame(rows, columns=[
        "subunit", "pos", "wt_aa", "mut_aa", "delta_charge",
        "axis_a_charge", "axis_b_connectivity", "axis_b_topology_only",
        "wire_class", "in_cluster", "is_cut_vertex", "fragments",
        "dist_to_loading_site", "cluster_waters_5A", "delta_volume",
        "delta_hydropathy", "burial"])


def separability(scores, column="axis_b_connectivity"):
    """Correlation between the axes and the size of the off-diagonal classes."""
    a = scores.axis_a_charge.to_numpy()
    b = scores[column].to_numpy()
    charged = scores.delta_charge != 0
    connectivity_only = (a < 0.005) & (b >= cfg.AXIS_B_THRESHOLD)
    charge_only = (a >= cfg.AXIS_A_THRESHOLD) & (b <= 0)
    return {
        "pearson_r_all": float(np.corrcoef(a, b)[0, 1]),
        "pearson_r_charge_changing": float(np.corrcoef(
            scores.loc[charged, "axis_a_charge"], scores.loc[charged, column])[0, 1]),
        "n_connectivity_only": int(connectivity_only.sum()),
        "residues_connectivity_only": int(
            scores[connectivity_only].groupby(["subunit", "pos"]).ngroups),
        "n_charge_only": int(charge_only.sum()),
        "residues_charge_only": int(
            scores[charge_only].groupby(["subunit", "pos"]).ngroups),
    }
```
