## Supplementary code for "Cancers relax selection on charge at the mitochondrial complex I quinone cavity": plotting.py

```
"""Shared figure style.

Black text on white throughout, type large enough to survive reduction to a
single journal column, and vector output alongside a raster copy.
"""
import matplotlib
import matplotlib.pyplot as plt

matplotlib.use("Agg")

CLASS_COLOURS = {
    "loading_site": "#1b4f72",
    "cut_vertex": "#2e86c1",
    "cluster_member": "#aab7b8",
    "other": "#d5d8dc",
}
CLASS_LABELS = {
    "loading_site": "Proton loading site",
    "cut_vertex": "Cut vertex",
    "cluster_member": "Cluster member",
    "other": "Other ND residue",
}
COHORT_MARKERS = {"germline": "o", "TCGA": "s", "PCAWG": "^",
                  "MSK-IMPACT": "D"}

STYLE = {
    "figure.dpi": 150,
    "savefig.dpi": 600,
    "figure.facecolor": "white",
    "axes.facecolor": "white",
    "savefig.facecolor": "white",
    "savefig.bbox": "tight",
    "font.family": "sans-serif",
    "font.sans-serif": ["Helvetica", "Arial", "DejaVu Sans"],
    "font.size": 11,
    "axes.titlesize": 12,
    "axes.labelsize": 11,
    "xtick.labelsize": 10,
    "ytick.labelsize": 10,
    "legend.fontsize": 10,
    "text.color": "black",
    "axes.labelcolor": "black",
    "axes.edgecolor": "black",
    "xtick.color": "black",
    "ytick.color": "black",
    "axes.spines.top": False,
    "axes.spines.right": False,
    "axes.linewidth": 0.8,
    "lines.linewidth": 1.4,
    "legend.frameon": False,
    "pdf.fonttype": 42,
    "ps.fonttype": 42,
}


def use_style():
    plt.rcParams.update(STYLE)


def panel_label(ax, letter, dx=-0.16, dy=1.04):
    ax.text(dx, dy, letter, transform=ax.transAxes, fontsize=14,
            fontweight="bold", color="black", va="bottom", ha="left")


def save(fig, directory, stem):
    """Write the vector and raster copies of one figure."""
    for suffix in ("pdf", "png"):
        fig.savefig(directory / f"{stem}.{suffix}")
    plt.close(fig)
    return directory / f"{stem}.pdf"
```
