## Supplementary code for "Cancers relax selection on charge at the mitochondrial complex I quinone cavity": variants.py

```
"""Substitution universe, mutational null model and selection statistics."""
from __future__ import annotations

import collections
import re

import numpy as np
import pandas as pd
from Bio.Seq import Seq
from scipy import stats

import config as cfg

COMPLEMENT = {"A": "T", "C": "G", "G": "C", "T": "A"}


def read_rcrs(path):
    """Read the revised Cambridge Reference Sequence as a plain string."""
    seq = "".join(line.strip() for line in open(path)
                  if re.fullmatch(r"[ACGTN]+", line.strip() or "x"))
    if len(seq) != 16569:
        raise ValueError(f"expected 16,569 bp, read {len(seq)}")
    return seq


def translate_nd_genes(seq):
    """Translate the ND genes in the vertebrate mitochondrial code.

    Returns the protein sequences and, for each residue, the rCRS coordinates
    of its three codon positions. Several ND genes end in an incomplete stop
    codon completed by polyadenylation, so the final partial codon is dropped.
    """
    proteins, codons = {}, {}
    for gene, (start, end, strand) in cfg.ND_GENES.items():
        nt = seq[start - 1:end]
        positions = list(range(start, end + 1))
        if strand == -1:
            nt = str(Seq(nt).reverse_complement())
            positions = positions[::-1]
        n = len(nt) // 3
        protein = str(Seq(nt[:n * 3]).translate(table=cfg.MITO_CODE)).rstrip("*")
        if len(protein) != cfg.ND_PROTEIN_LENGTHS[gene]:
            raise ValueError(f"{gene}: translated {len(protein)} aa, "
                             f"expected {cfg.ND_PROTEIN_LENGTHS[gene]}")
        proteins[gene] = protein
        codons[gene] = [tuple(positions[3 * i:3 * i + 3]) for i in range(len(protein))]
    return proteins, codons


def build_universe(seq, proteins, codons, classes):
    """Every possible single-nucleotide substitution in the ND genes.

    Each substitution is classified by translating the mutant codon, so the
    synonymous set is complete rather than inferred from a missense-only
    annotation database. Channels are strand-aware trinucleotides on the rCRS.
    """
    rows = []
    for gene, (start, end, strand) in cfg.ND_GENES.items():
        protein = proteins[gene]
        for index, triplet in enumerate(codons[gene]):
            wt_aa = protein[index]
            codon = "".join(seq[p - 1] if strand == 1 else COMPLEMENT[seq[p - 1]]
                            for p in triplet)
            wire_class = classes.get((gene, index + 1), "other")
            for ci, pos in enumerate(triplet):
                ref = seq[pos - 1]
                if ref not in COMPLEMENT or not 2 <= pos <= 16568:
                    continue
                for alt in "ACGT":
                    if alt == ref:
                        continue
                    base = alt if strand == 1 else COMPLEMENT[alt]
                    mutant = codon[:ci] + base + codon[ci + 1:]
                    mut_aa = str(Seq(mutant).translate(table=cfg.MITO_CODE))
                    kind = ("synonymous" if mut_aa == wt_aa
                            else "nonsense" if mut_aa == "*" else "missense")
                    rows.append((gene, index + 1, ci + 1, pos, ref, alt, wt_aa,
                                 mut_aa, kind, seq[pos - 2:pos + 1] + ">" + alt,
                                 wire_class, strand))
    return pd.DataFrame(rows, columns=[
        "subunit", "aa_pos", "codon_pos", "pos", "ref", "alt", "wt_aa",
        "mut_aa", "kind", "channel", "wire_class", "strand"])


def trinucleotide_spectrum(calls, seq, nd_genes):
    """Strand-aware 192-channel spectrum of a cohort.

    Estimated from all mtDNA single-nucleotide variants except coding variants
    in the genes under test, so the null is independent of the data analysed.
    """
    coding = {"Missense_Mutation", "Silent", "Nonsense_Mutation"}
    d = calls[(calls.Variant_Type == "SNP")
              & ~(calls.Hugo_Symbol.isin(nd_genes)
                  & calls.Variant_Classification.isin(coding))]
    d = d[d.Start_Position.between(2, 16568)]
    tri = d.Start_Position.map(lambda p: seq[p - 2:p + 1])
    ok = [isinstance(t, str) and t[1] == r
          for t, r in zip(tri, d.Reference_Allele)]
    channels = (tri[ok] + ">" + d.Tumor_Seq_Allele2[ok]).value_counts()
    total = channels.sum() + cfg.SPECTRUM_PSEUDOCOUNT * cfg.N_CHANNELS
    return (channels + cfg.SPECTRUM_PSEUDOCOUNT) / total


def observed_counts(calls, nd_genes, kind):
    """Map observed variants of one consequence onto (position, ref, alt)."""
    classification = {"missense": "Missense_Mutation", "synonymous": "Silent"}[kind]
    d = calls[calls.Hugo_Symbol.isin(nd_genes)
              & (calls.Variant_Classification == classification)
              & (calls.Variant_Type == "SNP")]
    return collections.Counter(zip(d.Start_Position, d.Reference_Allele,
                                   d.Tumor_Seq_Allele2))


def observed_expected(universe, spectrum, count_column, mask):
    """Observed/expected for a subset, with an exact two-sided Poisson test."""
    weights = universe.channel.map(spectrum).fillna(spectrum.min())
    total = universe[count_column].sum()
    expected = total * weights[mask].sum() / weights.sum()
    observed = int(universe.loc[mask, count_column].sum())
    if expected <= 0:
        return observed, np.nan, np.nan, np.nan
    p = (stats.poisson.sf(observed - 1, expected) * 2 if observed > expected
         else stats.poisson.cdf(observed, expected) * 2)
    return observed, expected, observed / expected, min(p, 1.0)


def poisson_interval(observed, expected):
    """Exact 95% Poisson interval on an observed/expected ratio."""
    low = stats.chi2.ppf(0.025, 2 * observed) / 2 if observed > 0 else 0.0
    high = stats.chi2.ppf(0.975, 2 * (observed + 1)) / 2
    return low / expected, high / expected


def benjamini_hochberg(pvalues):
    """Benjamini-Hochberg adjusted p values, in the input order."""
    p = np.asarray(pvalues, float)
    order = np.argsort(p)
    m = len(p)
    q = np.empty(m)
    running = 1.0
    for rank in range(m - 1, -1, -1):
        running = min(running, p[order[rank]] * m / (rank + 1))
        q[order[rank]] = running
    return q
```
