## Supplementary code for "Cancers relax selection on charge at the mitochondrial complex I quinone cavity": wire.py

```
"""Construction of the hydrogen-bonded proton wire and its residue classes."""
from __future__ import annotations

import collections

import gemmi
import networkx as nx
import numpy as np
from scipy.spatial import cKDTree

import config as cfg


def load_model(path, chains):
    """Read a structure and return its first model with one conformer per atom."""
    st = gemmi.read_structure(str(path))
    st.setup_entities()
    st.remove_alternative_conformations()
    return st[0]


def membrane_frame(model):
    """Membrane normal and in-plane axis from the deposited lipids.

    The lipids and detergents spread within the bilayer, so the least-variance
    principal axis of their atoms is the membrane normal and the greatest is
    the long in-membrane axis of the arm.
    """
    pts = [[a.pos.x, a.pos.y, a.pos.z]
           for ch in model for r in ch if r.name in cfg.LIPIDS for a in r]
    P = np.asarray(pts, float)
    centre = P.mean(0)
    _, _, Vt = np.linalg.svd(P - centre, full_matrices=False)
    return centre, Vt[2], Vt[0], P


def matrix_side(model, centre, normal):
    """Return +1 if the matrix lies on the +normal side, else -1.

    The FMN cofactor sits in the peripheral arm, which is in the matrix.
    """
    for ch in model:
        for r in ch:
            if r.name == "FMN":
                c = np.mean([[a.pos.x, a.pos.y, a.pos.z] for a in r], axis=0)
                return 1 if (c - centre) @ normal > 0 else -1
    raise ValueError("FMN not found; cannot orient the membrane normal")


def collect_nodes(model, chains):
    """Nodes of the wire: Grotthuss-competent side-chain atoms and waters.

    Returns a list of (key, residue_name, atom_name, xyz). The key is
    (subunit, seqid) for an ND residue, ("HOH", i) for a water and
    ("OTH", chain, seqid) for a lining residue of another subunit.
    """
    nd_atoms, nd_xyz = [], []
    for ch in model:
        if ch.name not in chains:
            continue
        sub = chains[ch.name]
        for r in ch:
            name = "MET" if r.name == "FME" else r.name
            if r.het_flag == "H" and r.name != "FME":
                continue
            if name not in cfg.WIRE_ATOMS:
                continue
            for a in r:
                if a.name in cfg.WIRE_ATOMS[name]:
                    nd_atoms.append(((sub, r.seqid.num), name, a.name,
                                     np.array([a.pos.x, a.pos.y, a.pos.z])))
                    nd_xyz.append(nd_atoms[-1][3])
    nd_xyz = np.asarray(nd_xyz)
    tree = cKDTree(nd_xyz)

    waters, lining = [], []
    for ch in model:
        for r in ch:
            if r.name == "HOH":
                p = np.array([r[0].pos.x, r[0].pos.y, r[0].pos.z])
                if tree.query(p)[0] <= cfg.WATER_SHELL:
                    waters.append((("HOH", len(waters)), "HOH", "O", p))
            elif ch.name not in chains and r.het_flag != "H":
                name = r.name
                if name not in cfg.WIRE_ATOMS:
                    continue
                for a in r:
                    if a.name not in cfg.WIRE_ATOMS[name]:
                        continue
                    p = np.array([a.pos.x, a.pos.y, a.pos.z])
                    if tree.query(p)[0] <= cfg.LINING_SHELL:
                        lining.append((("OTH", ch.name, r.seqid.num), name, a.name, p))
    return nd_atoms + waters + lining


def build_graph(nodes, cutoff):
    """Hydrogen-bond graph over the node set.

    The model carries no hydrogens, so a heavy-atom distance criterion is used,
    as is standard for structures without modelled protons. Pairs within one
    residue are excluded.
    """
    X = np.asarray([n[3] for n in nodes])
    keys = [n[0] for n in nodes]
    pairs = cKDTree(X).query_pairs(cutoff, output_type="ndarray")
    d = np.linalg.norm(X[pairs[:, 0]] - X[pairs[:, 1]], axis=1)
    keep = (d >= cfg.HBOND_MIN) & np.fromiter(
        (keys[a] != keys[b] for a, b in pairs), bool, len(pairs))
    pairs, d = pairs[keep], d[keep]

    g = nx.Graph()
    g.add_nodes_from(range(len(nodes)))
    g.add_weighted_edges_from(
        [(int(a), int(b), float(w)) for (a, b), w in zip(pairs, d)])
    return g, pairs, d, X


def classify(nodes, graph, proj_normal):
    """Assign each ND residue to a proton-wire class.

    loading_site    titratable and in a major hydrated cluster
    cut_vertex      an articulation point of its cluster
    cluster_member  otherwise in a cluster of at least MIN_CLUSTER nodes

    The hierarchy is strict and evaluated per residue, so a residue that is
    both titratable in a major cluster and an articulation point is a loading
    site.
    """
    keys = [n[0] for n in nodes]
    names = [n[1] for n in nodes]
    comps = [c for c in nx.connected_components(graph) if len(c) >= cfg.MIN_CLUSTER]
    spans = {k: float(np.ptp(proj_normal[list(c)])) for k, c in enumerate(comps)}
    major = {k for k, c in enumerate(comps)
             if len(c) >= cfg.MAJOR_MIN_NODES or spans[k] >= cfg.MAJOR_MIN_SPAN}

    articulation = set()
    for c in comps:
        articulation |= set(nx.articulation_points(graph.subgraph(c)))

    in_cluster, is_cut, is_site = set(), set(), set()
    for k, c in enumerate(comps):
        for i in c:
            if not isinstance(keys[i], tuple) or keys[i][0] not in cfg.SUBUNITS:
                continue
            in_cluster.add(keys[i])
            if i in articulation:
                is_cut.add(keys[i])
            if names[i] in cfg.TITRATABLE and k in major:
                is_site.add(keys[i])

    classes = {}
    for key in in_cluster:
        classes[key] = ("loading_site" if key in is_site
                        else "cut_vertex" if key in is_cut
                        else "cluster_member")
    return classes, comps, major, articulation


def residue_geometry(model, chains):
    """Side-chain centroid and burial for every modelled ND residue."""
    allpos = np.asarray([[a.pos.x, a.pos.y, a.pos.z]
                         for ch in model for r in ch if r.het_flag != "H" for a in r])
    tree = cKDTree(allpos)
    out = {}
    for ch in model:
        if ch.name not in chains:
            continue
        sub = chains[ch.name]
        for r in ch:
            if r.het_flag == "H" and r.name != "FME":
                continue
            name = "MET" if r.name == "FME" else r.name
            side = [a for a in r if a.name not in ("N", "CA", "C", "O", "OXT")]
            ref = side or [a for a in r if a.name == "CA"]
            if not ref:
                continue
            centroid = np.mean([[a.pos.x, a.pos.y, a.pos.z] for a in ref], axis=0)
            out[(sub, r.seqid.num)] = {
                "residue": name,
                "centroid": centroid,
                "burial": len(tree.query_ball_point(centroid, 8.0)),
            }
    return out
```
